# MolClaw: An Autonomous Agent with Hierarchical Skills for Drug Molecule Evaluation, Screening, and Optimization

**DOI:** 10.64898/2026.04.03.716272

**Authors:** Lisheng Zhang, Lilong Wang, Xiangyu Sun, Wei Tang, Haoyang Su, Yuehui Qian, Qikui Yang, Qingsong Li, Zhenyu Tang, Haoran Sun, Yingnan Han, Yankai Jiang, Wenjie Lou, Bowen Zhou, Xiaosong Wang, Lei Bai, Zhengwei Xie

## Abstract

Computational drug discovery, particularly the complex workflows of drug molecule screening and optimization, requires orchestrating dozens of specialized tools in multi-step workflows, yet current AI agents struggle to maintain robust performance and consistently underperform in these high-complexity scenarios. Here we present **MolClaw**, an autonomous agent that leads drug molecule evaluation, screening, and optimization. It unifies over 30 specialized domain resources through a three-tier hierarchical skill architecture (70 skills in total) that facilitates agent long-term interaction at runtime: tool-level skills standardize atomic operations, workflow-level skills compose them into validated pipelines with quality check and reflection, and a discipline-level skill supplies scientific principles governing planning and verification across all scenarios in the field. Additionally, we introduce **MolBench**, a benchmark comprising molecular screening, optimization, and end-to-end discovery challenges spanning 8 to 50+ sequential tool calls. MolClaw achieves state-of-the-art performance across all metrics, and ablation studies confirm that gains concentrate on tasks that demand structured workflows while vanishing on those solvable with ad hoc scripting, establishing workflow orchestration competence as the primary capability bottleneck for AI-driven drug discovery.

## Introduction

The discovery of new therapeutic molecules remains one of the most resource-intensive endeavors in modern science. Bringing a single drug from initial concept to regulatory approval typically requires 10 to 15 years of effort and over US$1 to 2 billion in investment, yet fewer than one in ten candidates entering Phase I clinical trials ultimately reach the market [54, 66]. Central to this attrition is the early-stage computational drug discovery pipeline, a complex, multi-step workflow spanning target structure determination, binding pocket identification, virtual screening, *de novo* molecular generation, molecular docking, binding free energy estimation, ADMET profiling, and iterative lead optimization [50, 62]. Each step relies on specialized software tools with disparate input/output formats, heterogeneous parameter spaces, and steep learning curves: a typical structure-based virtual screening campaign may require coordinated use of ESMFold or AlphaFold for protein structure prediction [37, 32], fpocket or P2Rank for pocket detection [34, 33], AutoDock Vina or Glide for molecular docking [22, 24], RDKit for physicochemical property calculation [48], and GROMACS for molecular dynamics refinement [1], often demanding days of manual effort to orchestrate the format conversions, parameter tuning, and quality control between each stage [19]. This fragmentation not only slows the pace of discovery but also introduces reproducibility challenges, as subtle differences in preprocessing steps or parameter choices can substantially alter downstream results [8].

The rapid advances in large language models (LLMs) over the past two years have opened a compelling new avenue for automating such complex scientific workflows. Foundation model series, e.g., GPT [2], Claude [4], Gemini [57], and open-source alternatives such as LLaMA [60] and Qwen [7] have demonstrated remarkable capabilities in natural language understanding, multi-step reasoning, and code generation. When augmented with the ability to invoke external tools, a paradigm known as tool-augmented LLMs or AI agents [52, 42], these models can go beyond text generation to plan and execute multi-step computational tasks. The ReAct framework [68], which interleaves reasoning traces with action execution, has become a foundational architecture for scientific agents, enabling LLMs to decompose complex objectives into sequences of tool calls, interpret intermediate results, and adaptively revise their strategies. This agentic paradigm has been recognized as a transformative force for scientific discovery [65, 10], with a recent perspective envisioning AI agents as collaborative partners that integrate computational models, biomedical tools, and experimental platforms to accelerate research workflows [25]. In parallel, general-purpose scientific agent frameworks such as InternAgent-1.5 [23], which coordinates generation, verification, and evolution subsystems for long-horizon autonomous discovery across computational and empirical domains, have further validated the feasibility of sustained multi-step scientific reasoning by AI systems. On the software infrastructure side, the emergence of open-source agent platforms, notably OpenClaw [43] and Claude Code [5], has significantly lowered the barrier to building tool-augmented scientific agents. These platforms provide robust runtime environments for tool orchestration, error recovery, and multi-turn reasoning, making it increasingly practical to deploy LLM-driven agents in real-world computational pipelines.

Several pioneering systems have begun to explore the intersection of LLM agents and chemistry or drug discovery. ChemCrow [41] integrated 18 chemistry tools with GPT-4 to accomplish tasks spanning organic synthesis planning, molecular property prediction, and materials design, and demonstrated the autonomous synthesis of an insect repellent through a cloud-connected robotic platform. ChemCrow established the viability of tool-augmented LLMs for chemistry but was limited to relatively simple, single-step tool invocations; it did not address the multi-stage workflows characteristic of computational drug discovery (e.g., pocket identification followed by molecular generation, docking, ADMET filtering, and binding energy estimation), nor did it include specialized tools for protein structure prediction, molecular dynamics, or free energy calculations. Biomni [28] constructed a broader biomedical action space by mining tools, databases, and protocols from 2,500 publications across 25 biomedical subfields. While Biomni demonstrated impressive generalization across heterogeneous tasks including gene prioritization, drug repurposing, and molecular cloning, its code-based execution paradigm, where the agent writes and runs ad hoc Python scripts for each task, lacks the structured, reusable workflow abstractions needed for the complex, multi-tool pipelines central to computational drug discovery. DrugAgent [38] introduced a multi-agent framework for automating machine learning programming in drug discovery tasks such as drug–target interaction prediction, but its scope was confined to ML model training rather than the execution of computational chemistry workflows involving 3D structure manipulation, molecular docking, or dynamics simulation. TxAgent [26], built on a finetuned 8B-parameter LLM with 211 biomedical tools, excelled at therapeutic reasoning tasks including drug information retrieval and personalized treatment recommendation, but operated entirely within the domain of clinical pharmacology and did not address the molecular-level computational tasks that constitute early-stage drug discovery. ChatInvent [27], deployed within AstraZeneca’s discovery pipeline, demonstrated real-world utility for molecular design and synthesis planning but remains a closed-source, proprietary system. More recently, the concept of “Prompt-to-Pill” [63] proposed an end-to-end multi-agent architecture from target identification to virtual patient recruitment, but was presented as a conceptual framework without a publicly available implementation or systematic evaluation on standardized benchmarks. Concurrent works including FROGENT [44], a multi-agent system integrating database, tool, and model layers via the Model Context Protocol; Mozi [14], which introduces governed autonomy with supervisor and worker hierarchies for long-horizon reliability; LIDDiA [6], which demonstrated autonomous navigation of the drug discovery process on 30 therapeutic targets. Yet none of these systems integrates the full spectrum of professional computational chemistry tools required for structure-based drug discovery, nor do they provide a systematic multi-dimensional benchmark for evaluation.

Here, we present MolClaw (shown in Figure 1), an autonomous AI agent built on a hierarchical skill architecture designed to enable the fully automated execution of end-to-end early-stage drug discovery workflows. To support this broad scope, MolClaw integrates over 30 professional computational chemistry and structural biology tools into a unified ecosystem. This comprehensive toolkit includes ESMFold [37] and Chai-1 [56] for protein structure prediction; fpocket [34] and P2Rank [33] for pocket identification; Vina-GPU 2.0 [20], DiffDock [16], and KarmaDock [69] for molecular docking; REINVENT 4 [39] for de novo molecular generation and scaffold hopping; PLIP [51] and ProLIF [11] for protein–ligand interaction fingerprinting; EquiScore [13] for structure-based scoring; Boltz-2 [45] for binding affinity prediction; GROMACS [1] and OpenMM [21] for molecular dynamics simulation; gmx_MMPBSA [61] for MM-PBSA binding free energy decomposition; ADMET-AI [55] for pharmacokinetic property prediction; ProteinMPNN [17] for protein sequence design; and RDKit [48] for general cheminformatics operations. All these heterogeneous tools are orchestrated through the Science Context Protocol (SCP), which is compatible with the Model Context Protocol (MCP) and ensures seamless GPU cluster scheduling, concurrent task management, and standardized access control [30].

**Figure 1:**
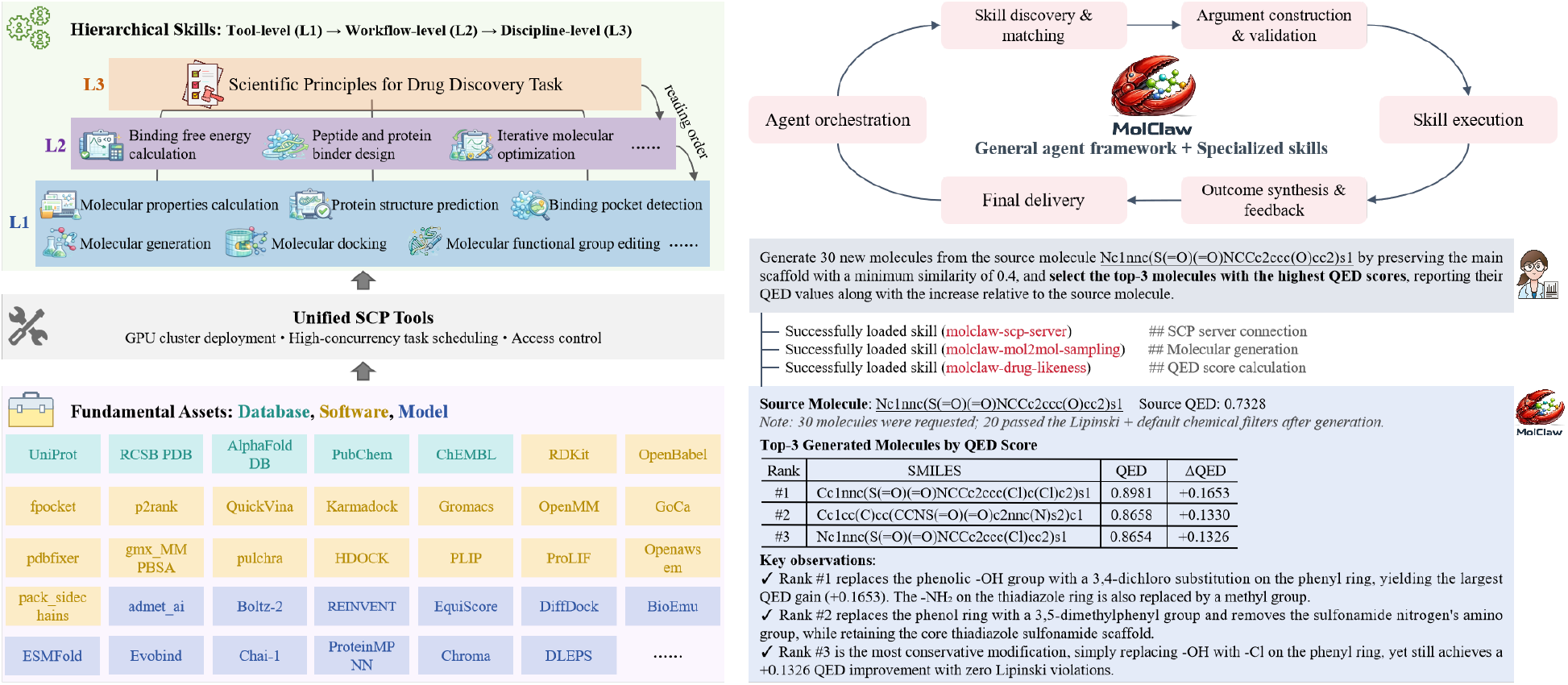
Technical architecture of MolClaw. The left side illustrates the construction of hierarchical skills, while the right side depicts the agent execution process and example interactions.

Building upon this robust SCP-standardized toolset, MolClaw employs a three-tier hierarchical skill architecture, comprising Tool-level (L1, foundational), Workflow-level (L2, intermediate), and Discipline-level (L3, highest strategic) skills, to encapsulate structure-based drug discovery domain expertise from concrete operational steps to overarching scientific governance. The foundational L1 includes over 58 fine-grained, task-specific templates wrapping individual tools or functionally related tool families, standardizing invocations, input/output validation, and quality control for core atomic operations. The intermediate L2 consists of 11 task-specific frameworks that compose validated L1 modules into end-to-end workflows for core drug discovery use cases (e.g., virtual screening, molecular optimization), defining tool selection, quality gates, and failure recovery while cross-referencing L3 principles to enforce scientific rigor. The highest L3 is a single comprehensive methodology document encoding 25 formal scientific principles for agent decision-making, quality verification, and reporting; loaded in full prior to any execution, it establishes universal scientific governance without dictating specific tool calls, ensuring robust methodology even for novel tasks. This hierarchical abstraction, analogous to the system call, library, and application layering in operating systems, serves three critical purposes: it encodes expert domain knowledge into reusable templates that are independent of the underlying LLM, it isolates errors at each level to prevent cascading failures, and it ensures reproducibility by standardizing the sequence and parameters of tool invocations across different runs and different LLM backbones. Importantly, MolClaw is designed to be both model-agnostic and framework-agnostic. The skill hierarchy is fully decoupled from LLM (tested with Claude Sonnet 4.6 and Kimi 2.5) as well as from the agent runtime. In practice, MolClaw has been deployed and evaluated on both OpenClaw [43] and Claude Code [5], two representative open-source agent platforms with distinct runtime architectures and skill-calling mechanisms. This cross-platform compatibility confirms that MolClaw’s capabilities improve automatically as both LLMs and agent platforms advance, without requiring modification to the skill definitions themselves.

To systematically evaluate MolClaw and establish a community resource for benchmarking drug discovery agents, we introduce MolBench, a multi-dimensional evaluation suite comprising three complementary tiers. MolBench-MS (Molecular Screening) evaluates the agent’s ability to correctly filter and rank molecules based on physicochemical constraints and activity predictions. MolBench-MO (Molecular Optimization) assesses the agent’s capacity to improve molecular properties through structural modifications while maintaining core scaffolds. MolBench-E2E (End-to-End Discovery) presents 3 comprehensive drug discovery challenges that each require the agent to autonomously execute extended, multi-phase workflows spanning 8–50+ sequential tool invocations, covering coarse-grained conformational sampling with dual force fields and all-atom reconstruction, multi-round closed-loop molecular property optimization, and structure-guided iterative lead optimization with docking-score-driven convergence. These scenarios closely mirror the complexity and the iterative, feedback-driven nature of real-world computational drug discovery campaigns.

We evaluate MolClaw against a comprehensive set of baselines, including frontier LLMs (GPT 5.2, Claude Sonnet 4.6, Gemini 3, DeepSeek v3.2, Qwen 3.5, Kimi 2.5, GLM 5, Minimax 2.5), the domain-specific biomedical agent Biomni [28], and vanilla agent frameworks (OpenClaw [43] and Claude Code without MolClaw skills). Across all three dimensions of MolBench, MolClaw achieves state-of-the-art performance, and ablation studies confirm that the performance gains derive primarily from the hierarchical skill architecture rather than the choice of underlying LLM.

In summary, this work makes the following contributions:

1. **A comprehensive tool ecosystem for computational drug discovery**. We integrate over 30 professional tools spanning the complete early-stage drug discovery pipeline, from protein structure prediction to molecular dynamics simulation, through a unified SCP infrastructure with GPU cluster scheduling, enabling fully automated execution of complex multi-tool workflows.
2. **An autonomous agent framework for drug discovery with a hierarchical skill scheme**. We propose the first three-level skill hierarchy (Tool, Workflow, and Pipeline) that encapsulates computational drug discovery expertise into reusable, composable, and model-agnostic skill templates, addressing the fundamental limitation of flat tool-calling in existing scientific agents.
3. **MolBench: the first multi-dimensional benchmark for drug discovery agents**. We construct and release MolBench, comprising molecular screening, molecular optimization, and 3 end-to-end drug discovery challenges that require genuine multi-step, multi-round tool execution across heterogeneous simulation and optimization pipelines, filling a critical gap in the evaluation of agentic systems for drug discovery.
4. **State-of-the-art performance with systematic evaluation**. We demonstrate that MolClaw substantially outperforms all baselines across MolBench dimensions, and through ablation studies establish that the hierarchical skill design, not the underlying LLM, is the primary driver of performance, validating the model-agnostic design philosophy.

## Results

### MolClaw achieves state-of-the-art performance across all MolBench evaluation dimensions

We evaluated MolClaw against eight frontier LLMs (GPT 5.2, Claude Sonnet 4.6, Gemini 3, DeepSeek v3.2, Qwen 3.5, Kimi 2.5, GLM 5, Minimax 2.5), the domain-specific biomedical agent Biomni, and two vanilla agent frameworks (Claude Code and OpenClaw without MolClaw skills). Across all seven metrics spanning the MolBench-MS and MolBench-MO benchmarks, MolClaw on Claude Code (MolClaw-CC) ranked first or tied for first on every metric, achieving sole best performance on four (binding affinity accuracy, docking hit count, molecule editing accuracy, and optimization delta) and tied best on three (property filtering accuracy and F1 score, optimization success rate) (Tables 1–2; Fig. 3). The multi-dimensional performance profile (Fig. 3I) shows that MolClaw-CC consistently encloses the largest area across all five normalized dimensions, a pattern that holds on both the Claude Code and OpenClaw platforms. Per-method confidence intervals, effect sizes, and pairwise statistical tests are presented in Fig. 4B–E, Fig. 5F–I, and Fig. 5J–M, respectively; complete numerical results are reported in the Supplementary Information.

**Table 1:** Performance comparison on the MolBench-MS benchmark across three sub-tasks: molecular property filtering, target-specific binding affinity comparison, and molecular docking screening. The best and second-best results are highlighted in bold and underline, respectively. Acc: accuracy. Here, Claude Code, OpenClaw and MolClaw all adopt Claude Sonnet 4.6 as their LLM backend.

| Method | Molecular Property Filtering |  | Binding Affinity Comparison | Molecular Docking Screening |
| --- | --- | --- | --- | --- |
|  | Acc (%) | F1 (%) | Acc (%) | Average hits@3 |
| *LLMs* |  |  |  |  |
| GPT 5.2 | 8.0 | 11.2 | 48.7 | 0.36 |
| Gemini 3 | 8.0 | 21.4 | 56.8 | 0.16 |
| Claude Sonnet 4.6 | 32.0 | 56.5 | 56.8 | 0.28 |
| Deepseek V3.2 | 20.0 | 34.9 | 43.2 | 0.08 |
| GLM 5 | 36.0 | 63.3 | 24.3 | 0.48 |
| Minimax 2.5 | 8.0 | 15.5 | 16.2 | 0.16 |
| Kimi 2.5 | 48.0 | 68.7 | 51.4 | 0.44 |
| Qwen 3.5 | 48.0 | 71.9 | 51.4 | 0.20 |
| *Agents* |  |  |  |  |
| Biomni | 74.0 | 82.7 | 24.3 | 0.00 |
| Claude Code (vanilla) | **98.0** | **99.3** | 51.4 | 0.56 |
| OpenClaw (vanilla) | 96.0 | 99.1 | 51.4 | 0.20 |
| MolClaw (Claude Code) | **98.0** | **99.3** | **81.1** | **0.80** |
| MolClaw (OpenClaw) | 96.0 | 98.9 | <u>73.0</u> | <u>0.64</u> |

**Table 2:** Performance comparison on the MolBench-MO benchmark across two sub-tasks: molecule editing and physicochemical property optimization. The best and second-best results are highlighted in bold and underline, respectively. Acc: accuracy, SR: success rate. Here, Claude Code, OpenClaw and MolClaw all adopt Claude Sonnet 4.6 as their LLM backend.

| Method | Molecule Editing | Molecule Optimization for Physicochemical Properties |  |
| --- | --- | --- | --- |
|  | Acc (%) | Delta | SR (%) |
| *LLMs* |  |  |  |
| GPT 5.2 | 84.6 | 0.428 | 82.1 |
| Gemini 3 | 51.0 | 0.574 | 89.7 |
| Claude Sonnet 4.6 | 94.9 | 1.023 | 97.4 |
| Deepseek V3.2 | 22.5 | 0.084 | 20.0 |
| GLM 5 | 74.4 | 0.697 | 84.6 |
| Minimax 2.5 | 30.8 | 0.003 | 5.1 |
| Kimi 2.5 | 89.7 | 0.783 | **100.0** |
| Qwen 3.5 | 90.0 | 0.716 | 87.5 |
| *Agents* |  |  |  |
| Biomni | 0.00 | 0.00 | 0.00 |
| Claude Code (vanilla) | <u>97.4</u> | 0.866 | 92.3 |
| OpenClaw (vanilla) | <u>97.4</u> | 1.293 | **100.0** |
| MolClaw (Claude Code) | **100.0** | **1.724** | **100.0** |
| MolClaw (OpenClaw) | <u>97.4</u> | <u>1.436</u> | **100.0** |

**Figure 2:**
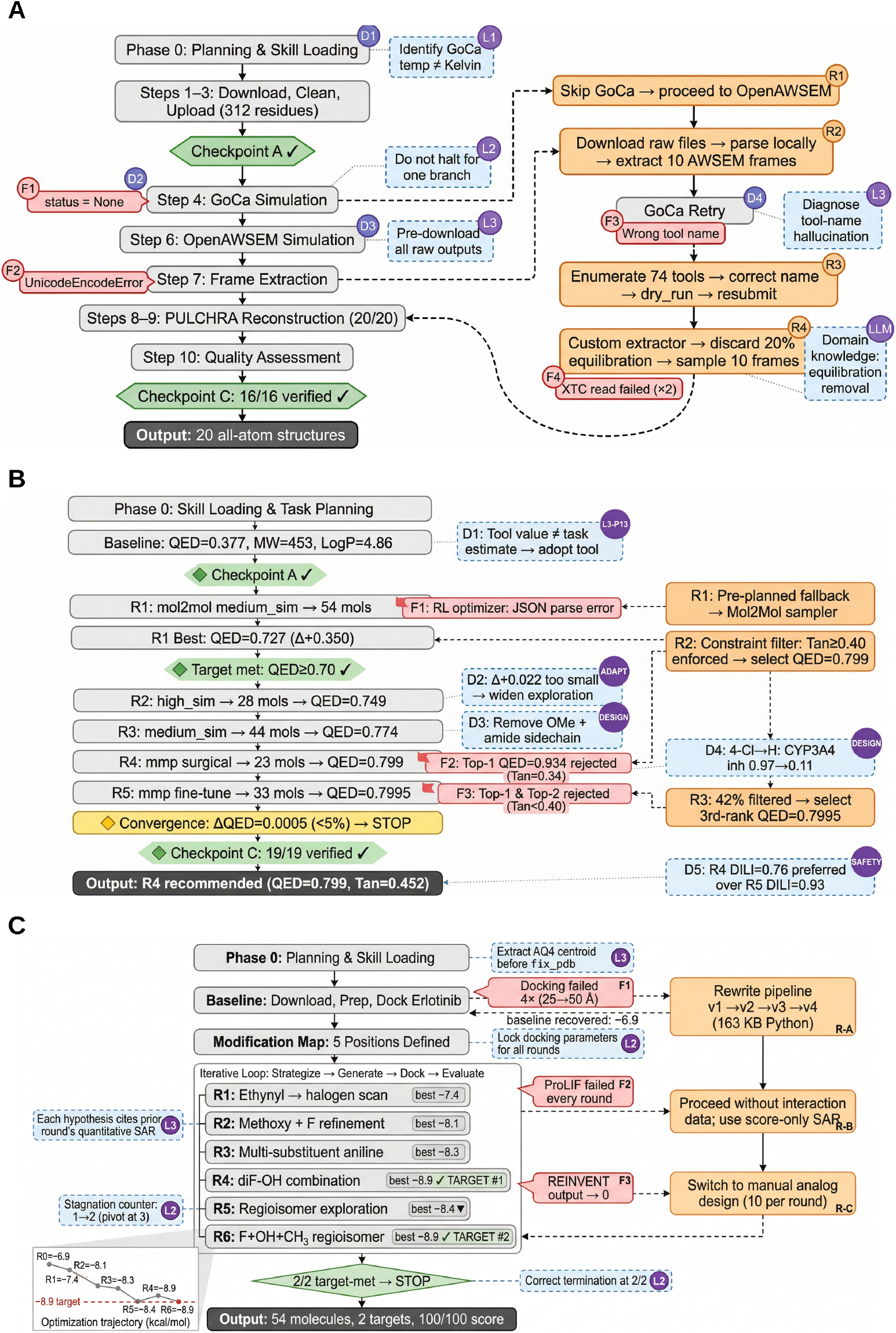
Agent execution traces for the three MolBench-E2E tasks. (**A**) E2E-Q1: coarse-grained conformational sampling. Five tool-level failures (red) were resolved via skill-governed recovery actions (orange), yielding 20 verified all-atom structures. (**B**) E2E-Q2: QED-driven iterative optimization. One tool fallback (F1), two constraint-driven rejections (F2–F3), and five strategy adaptations (D1–D5) were autonomously managed across five rounds, with all 19 reported values verified against source files. (**C**) E2E-Q3: structure-guided lead optimization of Erlotinib. Three failure categories—baseline docking crashes, persistent ProLIF errors, and generative-model collapse—were resolved via pipeline rewriting, score-only SAR, and manual analog design. Inset: optimization trajectory reaching the −8.9 kcal/mol target. Blue dashed boxes: agent decisions; purple badges: governing skill layer; green diamonds: verification check-points.

**Figure 3:**
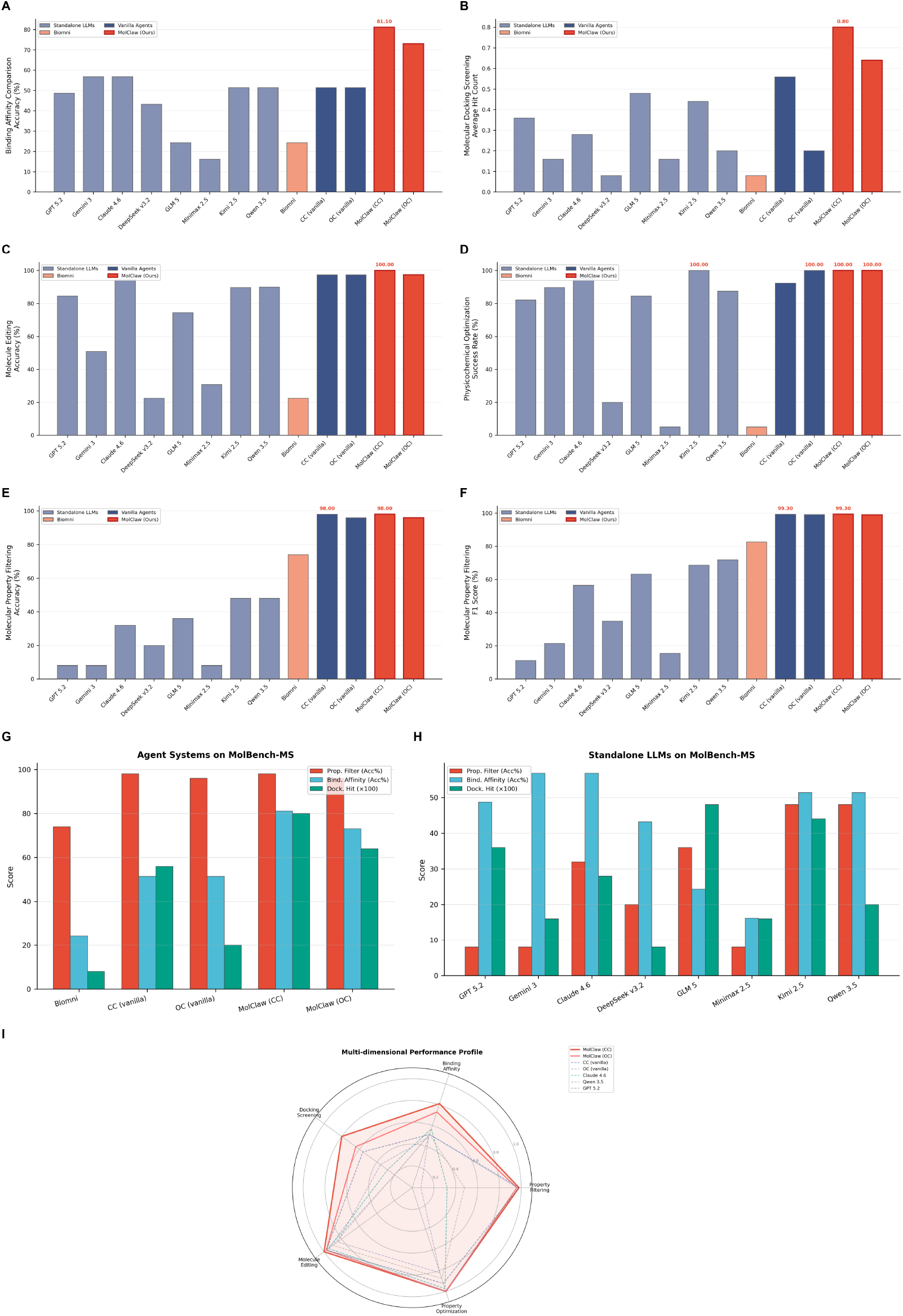
MolClaw achieves state-of-the-art performance across all MolBench evaluation dimensions. (**A**) Binding affinity comparison accuracy. MolClaw-CC achieves 81.1%. (**B**) Docking screening hit count. MolClaw-CC attains 0.80. (**C**) Molecule editing accuracy. MolClaw-CC reaches 100.0%. (**D**) Optimization success rate. (**E**) Property filtering accuracy. (**F**) Property filtering F1 score. (**G**) Agent systems grouped comparison across three MS sub-tasks. (**H**) Standalone LLMs grouped comparison. (**I**) Radar chart of normalized multi-dimensional performance for seven representative methods. Bars color-coded: LLMs (light blue-grey), Biomni (coral), vanilla agents (dark blue), MolClaw (red). “fail”: no valid output.

**Figure 4:**
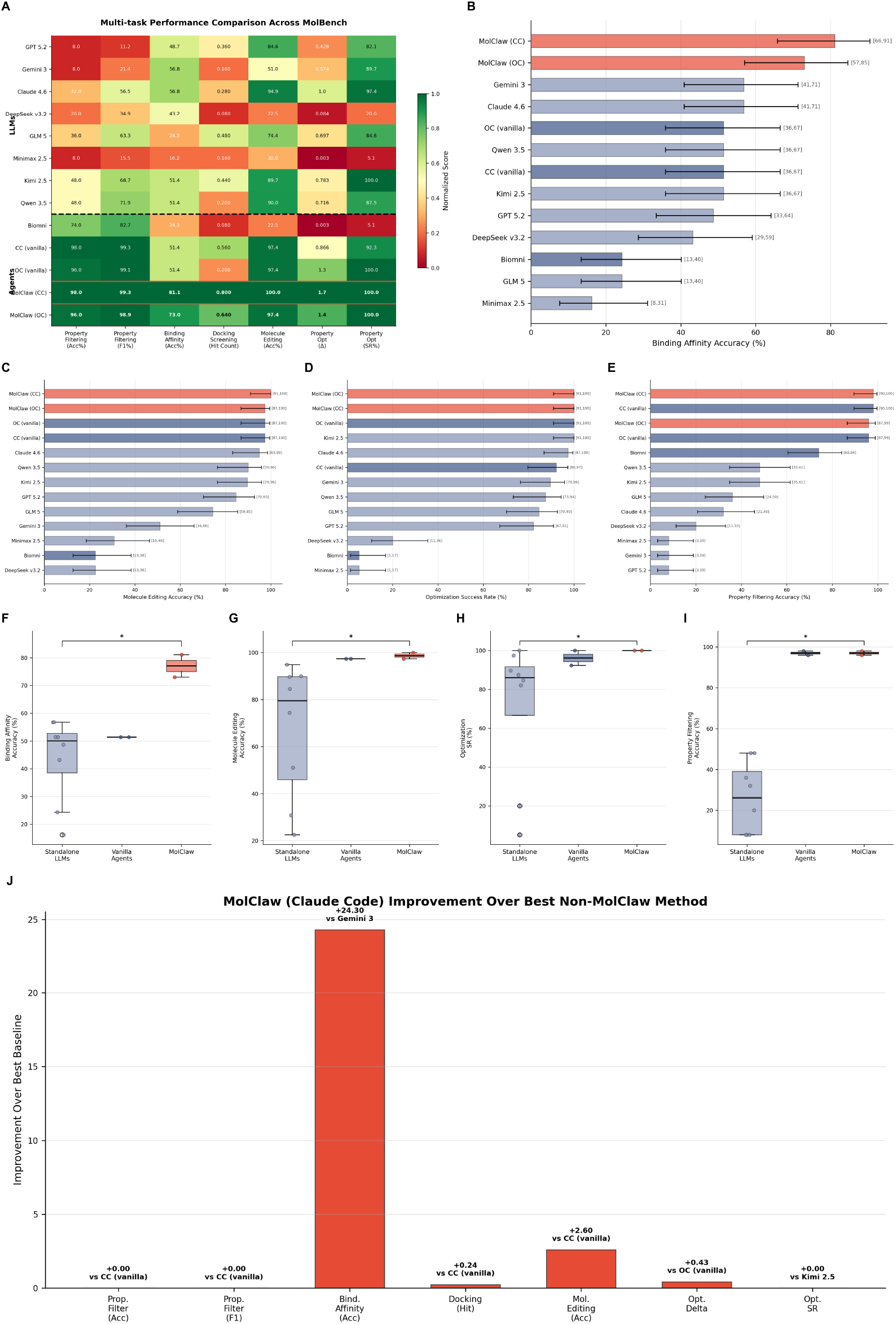
Statistical validation confirms the significance and reliability of MolClaw’s performance advantages. (**A**) Normalized performance heatmap across seven metrics for 13 methods. MolClaw variants highlighted by red borders. (**B–E**) Wilson score 95% CI forest plots for binding affinity accuracy (**B**), molecule editing accuracy (**C**), optimization success rate (**D**), and property filtering accuracy (**E**). (**F–I**) Category-level box-and-strip plots comparing LLMs (*n* = 8), vanilla agents (*n* = 2), and MolClaw (*n* = 2). ^∗^*P <* 0.05, Mann–Whitney *U*. (**J**) Waterfall chart of MolClaw-CC improvement over best non-MolClaw baseline per metric.

**Figure 5:**
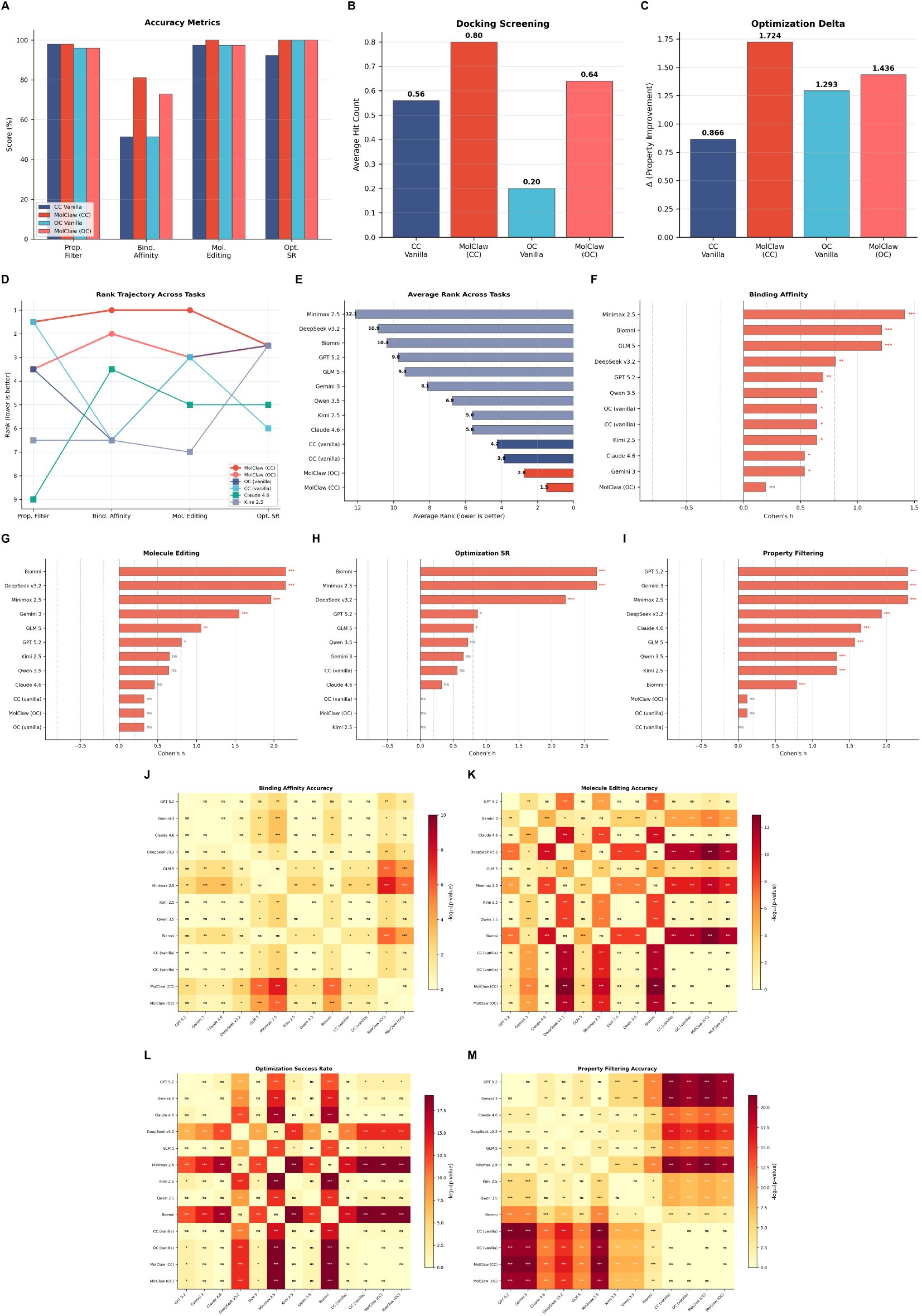
Ablation studies and in-depth statistical analyses reveal the mechanistic basis of MolClaw’s superiority. (**A–C**) Ablation on Claude Code and OpenClaw platforms: accuracy metrics (**A**), docking hit count (**B**), optimization delta (**C**). Largest skill-driven gain: binding affinity +29.7 pp (*P* = 0.013, *h* = 0.64). (**D**) Rank trajectory across four tasks for top six methods. (**E**) Average rank (Friedman *χ*^2^ = 35.35, *P* = 2.17 × 10^−4^); MolClaw-CC best at 1.5. (**F–I**) Cohen’s *h* effect sizes for MolClaw-CC vs. each baseline. Dashed lines: small/medium/large thresholds. (**J–M**) Pairwise Fisher’s exact test matrices for all 13 methods. ^∗^*P <* 0.05; ^∗∗^*P <* 0.01; ^∗∗∗^*P <* 0.001.

More importantly, the results reveal a fundamental insight about when hierarchical skills matter: MolClaw’s advantages are not uniform across tasks but are concentrated precisely on those that demand structured multi-step computational workflows, as detailed below.

### Molecular screening reveals a performance hierarchy governed by task complexity

The three MolBench-MS sub-tasks (Fig. 3A–B, E–H) expose a clear, three-tier performance hierarchy that directly reflects the value of hierarchical skill design.

On molecular property filtering (*n* = 50 queries), which requires computing physicochemical descriptors and applying threshold-based constraints, standalone LLMs performed poorly (mean accuracy 26.0%, range 8.0–48.0%; Fig. 3E), confirming that LLMs cannot reliably compute molecular descriptors through reasoning alone. However, all agent-based systems, including vanilla frameworks without MolClaw skills, achieved near-perfect accuracy (≥ 96.0%) by autonomously writing and executing RDKit-based scripts (Table 1). MolClaw-CC matched Claude Code vanilla at 98.0%, demonstrating that pre-defined skills confer no additional advantage when ad hoc scripting suffices. This task thus serves as an important control: tool access, not skill design, is the performance bottleneck.

The picture changes dramatically on binding affinity comparison (*n* = 37 molecular pairs), the most discriminating task in MolBench. Standalone LLMs achieved a mean accuracy of only 43.6%, near chance level for this binary task, with the best-performing LLMs (Gemini 3 and Claude Sonnet 4.6) reaching only 56.8% (Fig. 3A). Critically, vanilla agent frameworks showed no improvement over the best LLMs (both at 51.4%), indicating that unstructured code generation alone cannot address this challenge. In contrast, MolClaw-CC achieved 81.1% accuracy, a 24.3 percentage point gain over the best non-MolClaw baseline (Fisher’s exact test, *P* = 0.043, Cohen’s *h* = 0.54) and a 29.7 percentage point gain over its own vanilla backbone (*P* = 0.013, *h* = 0.64; Fig. 5A). This substantial improvement reflects MolClaw’s workflow for binding affinity prediction, which orchestrates target protein preparation, protein-ligand binding execution, and scoring into a validated multi-step pipeline that encodes domain expertise inaccessible to ad hoc approaches.

Molecular docking screening (*n* = 25 targets) confirmed this pattern: MolClaw-CC achieved the highest average hit count of 0.80, outperforming Claude Code vanilla (0.56) by 0.24 and OpenClaw vanilla (0.20) by a threefold margin on MolClaw-OC (0.64; Fig. 3B, Fig. 5B). Biomni failed to produce valid docking outputs entirely. These consistent improvements across both platforms indicate that MolClaw’s docking-specific skills, which chains receptor preparation, docking box definition, Vina-GPU execution, and result post-processing, confer a systematic advantage that scales beyond any single platform. Detailed sub-task breakdowns for each agent and LLM are shown in Fig. 3G–H.

### Molecular optimization confirms the value of iterative refinement skills

The MolBench-MO benchmark assesses two complementary molecular design capabilities (Table 2; Fig. 3C–D).

On molecule editing (*n* = 39 tasks), which evaluates the ability to apply specified chemical transformations such as functional group replacement and scaffold hopping, standalone LLMs showed high variance (accuracy range 22.5–94.9%, mean 67.2%). Both vanilla agent frameworks already achieved 97.4% accuracy, and MolClaw-CC reached a perfect 100.0% (a 2.6 percentage point gain), while not individually significant (*P* = 1.00), establishes MolClaw-CC as the only method to achieve perfect accuracy on this benchmark. Comparisons with standalone LLMs yielded large effect sizes (Cohen’s *h* up to 2.15 versus DeepSeek v3.2, *P* = 1.23 × 10^−13^; Fig. 5G). These results suggest that molecule editing is primarily a molecular representation manipulation task where code-generating agents already perform well, and hierarchical skills provide incremental rather than transformative improvement.

Physicochemical property optimization, by contrast, represents the most challenging task in MolBench and the clearest demonstration of MolClaw’s iterative molecular optimization skill. This task requires improving three key physicochemical properties (QED, LogP, LogS), demanding iterative cycles of structural modification, computational verification, and strategy revision. Standalone LLMs achieved a mean optimization delta (Δ) of only 0.538 (range 0.003–1.023), with success rates as low as 5.1% (Minimax 2.5), revealing that most LLMs lack the consistency required for reliable multi-objective optimization (Table 2). Claude Code vanilla achieved Δ = 0.866 with 92.3% success rate, while OpenClaw vanilla reached Δ = 1.293 with 100.0% success rate. MolClaw-CC achieved the highest Δ of 1.724, double that of Claude Code vanilla, with a perfect 100.0% success rate (Fig. 3D, Fig. 5C). This improvement reflects MolClaw’s L2 workflow skill for iterative molecular optimization, which integrates LLM-guided structural design, property computation, Lipinski violation checking, and automated evaluation into a dynamic loop that systematically explores the chemical space until all targets are simultaneously satisfied.

### Ablation and statistical analyses confirm hierarchical skills as the primary performance driver

To isolate the contribution of MolClaw’s skill architecture from platform-specific effects, we performed controlled ablation experiments on both agent platforms (Fig. 5A–C). The results reveal a consistent pattern: skills provide the greatest benefit on tasks requiring domain-specific computational workflows, while conferring negligible advantage on tasks solvable by general-purpose scripting.

On the Claude Code platform, MolClaw’s hierarchical skills elevated binding affinity accuracy from 51.4% to 81.1% (+29.7 percentage points, *P* = 0.013, *h* = 0.64), docking hit count from 0.56 to 0.80, and optimization delta from 0.866 to 1.724 (2.0 × increase). On the OpenClaw platform, analogous improvements were observed: binding affinity accuracy rose from 51.4% to 73.0% (+21.6 percentage points), docking hit count more than tripled from 0.20 to 0.64, and optimization delta increased from 1.293 to 1.436. The consistency of these results across two architecturally distinct platforms substantiates MolClaw’s model-agnostic design philosophy: the skill layer, not the underlying LLM or runtime, is the primary driver of performance.

Cross-task ranking analysis further supports this conclusion. A Friedman test over four accuracy-based metrics across 12 methods with complete data (Biomni excluded due to missing values) revealed significantly non-uniform rank distributions (*χ*^2^ = 35.35, *P* = 2.17 × 10^−4^). MolClaw-CC achieved the best average rank of 1.5 out of 12, maintaining rank 1 on three of four tasks (Fig. 5D–E). Cohen’s *h* effect sizes between MolClaw-CC and all baselines were uniformly positive, with the majority exceeding the large-effect threshold (|*h* | *>* 0.8), most notably on property filtering (*h* = 2.28 versus GPT 5.2, Gemini 3, and Minimax 2.5) and binding affinity (*h* = 1.41 versus Minimax 2.5) (Fig. 5F–I). Pairwise significance matrices (Fisher’s exact test; Fig. 5J–M) confirmed that MolClaw-CC achieved statistically significant superiority (*P <* 0.05) over the majority of baselines on binding affinity, molecule editing, and property filtering; the few non-significant comparisons occurred where baseline performance was already near ceiling. Wilson score 95% confidence intervals for all accuracy-based metrics are shown in Fig. 4B–E, and category-level distributions (standalone LLMs versus vanilla agents versus MolClaw) are compared in Fig. 4F–I, with all comparisons reaching significance (Mann–Whitney *U* test, *P <* 0.05). A complete description of statistical methodology and full numerical test results is provided in the Supplementary Information.

### MolClaw effectively addresses complex, long-horizon drug discovery tasks

#### Coarse-grained conformational sampling and all-atom reconstruction of the EGFR kinase domain (E2E-Q1)

To evaluate the agent’s capacity for conformational sampling, a prerequisite for cryptic allosteric site discovery, we tasked it with performing dual coarse-grained (CG) molecular dynamics simulations of the EGFR kinase domain (PDB: 1M17, chain A, 312 residues) using GoCa and OpenAWSEM force fields, followed by PULCHRA all-atom reconstruction. Superposition of the ten reconstructed ensembles onto the native crystal structure (Fig. 6A, B) revealed that both pipelines were executed end-to-end without manual intervention, producing physically plausible conformational ensembles suitable for downstream pocket analysis.

**Figure 6:**
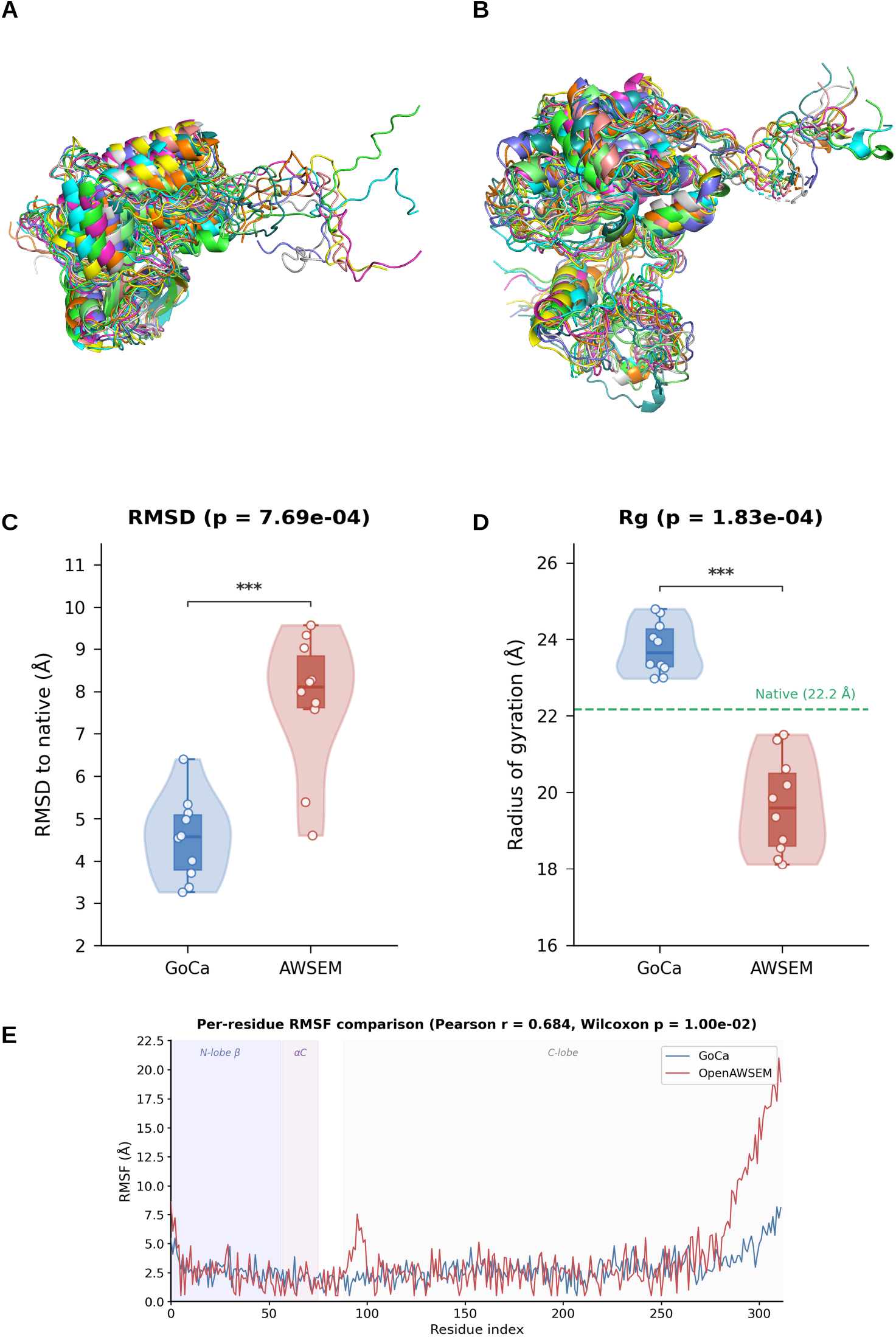
Coarse-grained conformational sampling of the EGFR kinase domain by OpenAWSEM and GoCa. (**A**) Superposition of 10 PULCHRA-reconstructed all-atom conformations from the OpenAWSEM ensemble, aligned to the 1M17 crystal structure. (**B**) Corresponding superposition for the GoCa ensemble. (**C**) C*α*-RMSD to native structure: GoCa 4.54 ± 0.93 Å versus AWSEM 7.78 ± 1.53 Å (*P* = 7.69 × 10^−4^). (**D**) Radius of gyration: GoCa *R*_g_ = 23.77 ± 0.65 Å, AWSEM *R*_g_ = 19.66 ± 1.18 Å, native *R*_g_ = 22.2 Å (dashed line; *P* = 1.83 × 10^−4^). (**E**) Pairwise C*α*-RMSD within each ensemble (*n* = 45 pairs): GoCa 4.28 ± 0.88 Å versus AWSEM 5.67 ± 1.60 Å (*P* = 2.44 × 10^−5^). (**F**) Per-residue RMSF profiles across 312 residues (Pearson *r* = 0.685); shaded regions denote the N-lobe *β*-sheet, *α*C-helix, and C-lobe. Violin plots show individual data points with box-and-whisker summaries; *P*-values from two-sided Mann–Whitney *U* tests; ∗∗∗*P <* 0.001; *n* = 10 conformations per method.

Quantitative assessment showed that GoCa, a native-topology Gō model, sampled conformations confined near the native basin (C*α*-RMSD 3.25–6.40 Å), whereas OpenAWSEM explored substantially broader conformational space (4.61–9.58 Å; Mann–Whitney *U, P* = 7.69 × 10^−4^; Fig. 6C). This broader sampling by OpenAWSEM is particularly relevant for cryptic site discovery, as transient pocket opening events typically require larger-amplitude backbone rearrangements that deviate significantly from the holo crystal structure.

The two force fields exhibited opposing trends in global compactness (Fig. 6D). GoCa structures were mildly expanded relative to the native state (mean *R*_g_ = 23.8 Å versus native 22.2 Å), while OpenAWSEM produced notably more compact conformations (mean *R*_g_ = 19.7 Å; *P* = 1.83 × 10^−4^). This compaction may reflect AWSEM’s cooperative folding energy landscape favoring tight hydrophobic packing, potentially exposing surface cryptic pockets through differential domain rearrangement rather than global swelling. Pairwise C*α*-RMSD among the ten sampled conformations further confirmed that OpenAWSEM generated greater structural diversity (mean 5.67 Å versus 4.28 Å; *P* = 2.44 × 10^−5^; Fig. 6E), providing a broader ensemble for subsequent pocket detection.

Despite these quantitative differences, per-residue RMSF profiles from the two methods were moderately correlated (Pearson *r* = 0.685; Fig. 6F), indicating consensus on the identity of flexible regions, including the N-/C-terminal tails, the *α*C–*β*4 loop, and the activation segment, consistent with experimentally characterized EGFR dynamics implicated in allosteric regulation. The agent autonomously completed the full workflow encompassing structure cleaning, CG model preparation, MD execution, trajectory extraction, and all-atom reconstruction with appropriate quality controls, demonstrating its competence in conformational sampling tasks essential for structure-based drug discovery beyond the static crystallographic snapshot.

Beyond the scientific outcomes, the execution trace of E2E-Q1 (Fig. 2A) reveals MolClaw’s realtime adaptive capabilities under cascading tool failures. The agent encountered five distinct failures across the two simulation engines: GoCa returned empty outputs due to an incorrect tool name (the agent had invoked `goca_pipeline` rather than the registered `run_goca_pipeline`), the OpenAWSEM frame extraction tool crashed with a Unicode encoding error, and two subsequent attempts to parse the GoCa trajectory via MDTraj failed due to topology mismatches. In each case, the hierarchical skill architecture guided recovery. Upon the initial GoCa failure, the L2 workflow skill directed the agent to proceed with OpenAWSEM first and defer the retry, a non-blocking strategy that preserved pipeline progress. When the OpenAWSEM extraction tool failed, the agent, following L3 Principle 14 (mandatory file collection) had already proactively downloaded all raw simulation outputs, enabling it to parse the trajectory PDB locally, map non-standard AWSEM residue names to canonical codes, and extract 10 frames without server-side tool support. For the GoCa retry, the agent autonomously enumerated all 74 tools registered on the server to diagnose the naming discrepancy, executed a dry-run validation before resubmission, and upon trajectory extraction failure, wrote a custom extractor that identified 45 frames in the XTC trajectory, discarded the first 20% as equilibration, and sampled 10 evenly spaced conformations. The entire workflow, spanning seven tool types and four recovery episodes, completed autonomously, producing 20 all-atom structures that passed all 16 pre-report data integrity checks.

#### QED-driven iterative molecular optimization reveals scaffold-imposed ceilings and agent blind spots (E2E-Q2)

We tasked the agent with iteratively optimizing the quantitative estimate of drug-likeness (QED) of a triazolo-benzodiazepine compound (MW = 452.9 Da, QED = 0.377) over five rounds, subject to dual constraints of QED ≥ 0.70 and Tanimoto similarity ≥ 0.40 to the starting molecule. The agent executed a complete Assess–Diagnose–Design–Verify loop at each round, using REINVENT4 mol2mol sampling for candidate generation, RDKit for property computation, and ADMET-AI for safety profiling. The full optimization trajectory is shown in Fig. 7.

**Figure 7:**
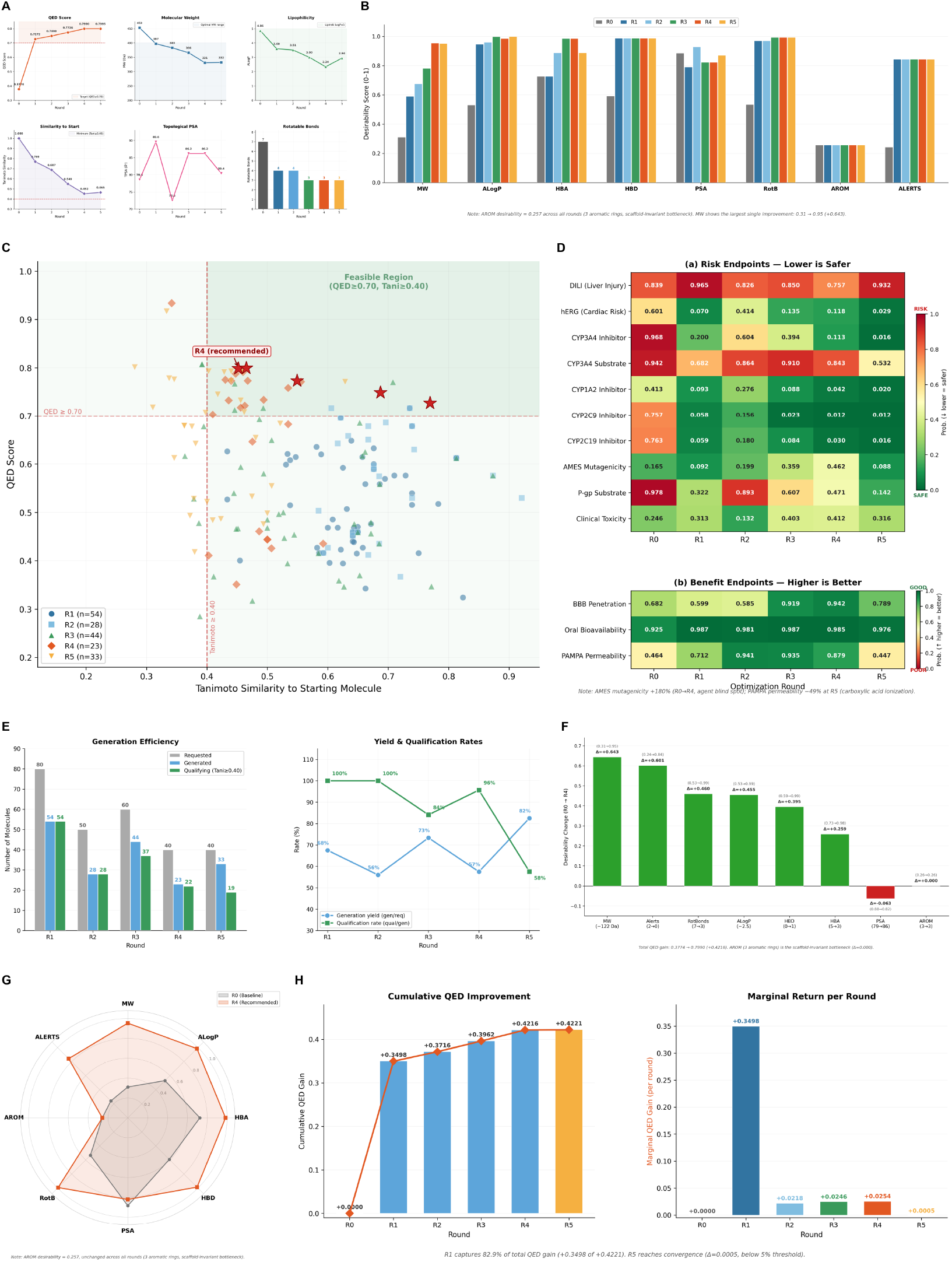
QED-driven iterative optimization of a triazolo-benzodiazepine scaffold by the AI agent. **(A)** Multi-dimensional property trajectory across five optimization rounds: QED score (target ≥ 0.70), MW, ALogP, Tanimoto similarity (constraint ≥ 0.40), TPSA and rotatable bonds. (**B**) QED desirability decomposition by component and round (R0–R5). (**C**) QED–Tanimoto trade-off for all 182 molecules; red stars, selected best per round; green quadrant, feasible region. (**D**) ADMET profile evolution (top: risk endpoints; bottom: benefit endpoints). (**E**) Generation efficiency: molecules requested, generated and qualifying per round (left); yield and qualification rates (right). (**F**) QED component contribution waterfall (R0 → R4). (**G**) Radar chart comparing R0 and R4 desirability profiles. (**H**) Cumulative QED gain (left) and marginal gain per round (right). QED and Tanimoto computed via RDKit (Morgan fingerprint, radius = 2, 2,048 bits); ADMET by ADMET-AI; *n* = 182 molecules (54 + 28 + 44 + 23 + 33).

The agent achieved the QED target in a single round (R1, QED = 0.727) by replacing the butyl ester side chain with a free carboxylic acid, simultaneously reducing molecular weight by 56 Da, removing three rotatable bonds, and clearing two Brenk structural alerts (Fig. 7A). This single modification captured 82.9% of the total five-round QED improvement (Fig. 7H, right panel). Subsequent rounds yielded progressively smaller gains through targeted modifications: carboxylic acid to ethanol (R2, +0.022), methoxy group removal combined with alcohol-to-amide conversion (R3, +0.025), and chlorine removal from the pendant phenyl ring (R4, +0.025). By R5, the marginal gain fell to +0.0005, and the agent correctly declared convergence (Fig. 7H). The final recommended molecule (R4; QED = 0.799, Tanimoto = 0.452) represents a 112% improvement over baseline.

To understand why QED plateaued near 0.80, we decomposed the score into its eight constituent desirability functions (Fig. 7B, F). Six components improved dramatically from R0 to R4: MW (+0.643), structural alerts (+0.601), rotatable bonds (+0.460), ALogP (+0.455), HBD (+0.395) and HBA (+0.259). However, the aromatic ring desirability (AROM) remained fixed at 0.257 across all rounds, the triazolo-benzodiazepine scaffold inherently contains three aromatic rings, and this cannot be altered without breaking the core pharmacophore or violating the Tanimoto constraint. The radar comparison of R0 versus R4 profiles (Fig. 7G) visualizes this ceiling effect: the R4 polygon approaches a near-regular octagon except for a persistent indentation at the AROM axis. Using the weighted geometric mean formulation of QED, we derived a theoretical ceiling of 0.846; the achieved value of 0.799 represents 94–95% of this bound.

The QED–Tanimoto scatter of all 182 generated molecules (Fig. 7C) reveals a fundamental trade-off: higher QED generally requires greater structural divergence. In rounds 3–5, the global highest-QED molecules (up to 0.934) fell below the Tanimoto threshold and were correctly excluded by the agent. Generation efficiency data (Fig. 7E) further quantify this tension: the fraction of molecules satisfying the Tanimoto constraint declined from 100% (R1) to 57.6% (R5) as the iterative seed progressively diverged from the starting structure.

Beyond QED optimization, the agent tracked multiple ADMET endpoints across rounds (Fig. 7D). CYP3A4 inhibition probability decreased from 0.968 to 0.113 (−88%), hERG cardiac risk from 0.601 to 0.118 (−80%), and aqueous solubility improved by 2.6 log units. The agent also demonstrated multi-objective judgement by selecting R4 over R5, correctly prioritizing the compound with the superior overall ADMET profile despite R5 having marginally higher QED. However, a critical blind spot emerged: AMES mutagenicity probability rose from 0.165 to 0.462 (+180%) between R0 and R4, approaching the positive threshold, yet this deterioration was never flagged in the agent’s reports. This omission likely reflects an attentional bias toward endpoints that were problematic at baseline while neglecting those that started in a safe range.

The execution trace of E2E-Q2 (Fig. 2B) further illustrates the agent’s real-time adaptive reasoning across the five-round optimization loop. In Round 1, the primary generation tool returned a JSON parsing error; the agent immediately executed its pre-planned fallback strategy, switching to an alternative tool without retrying, as this decision logic was encoded during Phase 0 planning. More subtly, the agent detected that the tool-computed baseline QED (0.377) diverged substantially from the task description’s estimate (0.50–0.55) and autonomously adopted the tool-derived value as ground truth throughout all subsequent rounds, following L3 Principle 13 (computation-first data authority). Strategy adaptation was explicitly conditioned on prior quantitative outcomes: when Round 2’s conservative high-similarity prior yielded only ΔQED = +0.022, the agent diagnosed the gain as insufficient and reverted to medium-similarity exploration for Round 3, subsequently escalating to matched molecular pair priors for surgical single-site modifications in Rounds 4–5. A critical constraint-enforcement pattern emerged in later rounds: in Round 4 the highest-QED molecule (0.934) was rejected because its Tanimoto similarity (0.34) fell below the 0.40 threshold, and in Round 5 the top two candidates were likewise excluded (42% rejection rate), demonstrating that the agent consistently prioritized constraint compliance over objective maximization. The three-tier verification system (Checkpoint A after each tool call, Checkpoint B per round, Checkpoint C pre-report) audited all 19 key numerical values against their source files with zero discrepancies.

#### Structure-guided iterative lead optimization of Erlotinib targeting EGFR (E2E-Q3)

### Baseline establishment and locked docking parameters

The AI agent downloaded the EGFR kinase domain co-crystal structure (PDB: 1M17, chain A, 2.6 Å resolution) and extracted the AQ4 ligand centroid (22.014, 0.253, 52.794) from HETATM records prior to receptor preparation. After removing heterogens and waters and adding hydrogens at pH 7.0 via PDBFixer, the receptor was converted to PDBQT format and Erlotinib was docked using QuickVina2 with a 25 × 25 × 25 Å box centered on the AQ4 centroid. The resulting baseline docking score was −6.9 kcal/mol, establishing a target threshold of −8.9 kcal/mol (ΔScore ≤ −2.0). Erlotinib’s computed molecular properties (MW = 393.4 Da, LogP = 3.41, HBD = 1, HBA = 7, rotatable bonds = 10, TPSA = 74.7 Å^2^) confirmed zero Lipinski violations and Veber compliance. These docking parameters were locked for all subsequent rounds (Fig. 8A).

**Figure 8:**
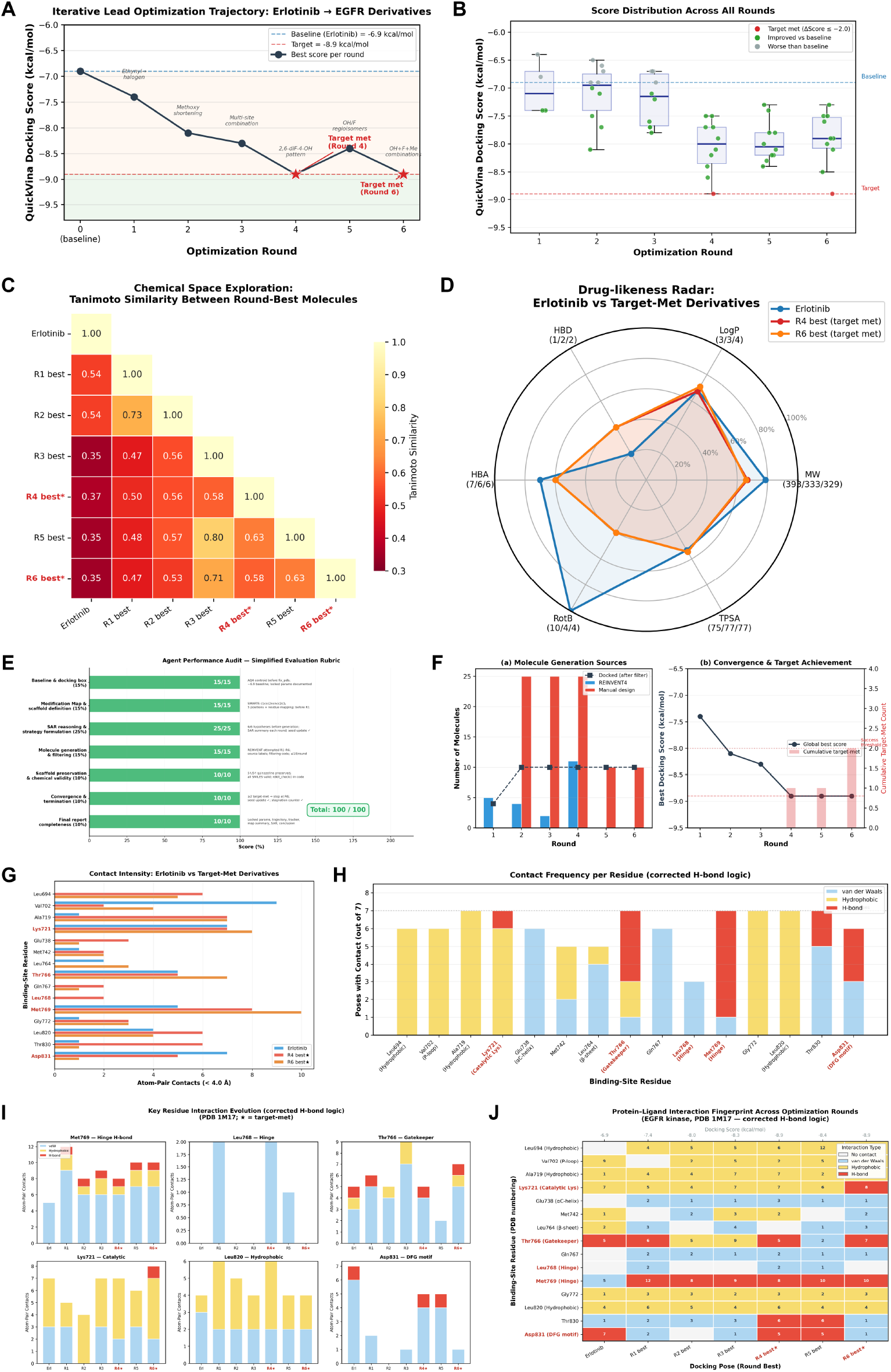
Comprehensive evaluation of AI-agent-driven iterative lead optimization of Erlotinib targeting the EGFR kinase domain. (**A**) Optimization trajectory showing best QuickVina docking score per round; blue dashed line: Erlotinib baseline (−6.9 kcal/mol); red dashed line: −8.9 kcal/mol target. (**B**) Docking score distributions across Rounds 1–6 (box-and-strip plot, *n* = 54). (**C**) Tanimoto similarity heatmap between round-best molecules. (**D**) Drug-likeness radar plot comparing Erlotinib with two target-met derivatives. (**E**) Agent performance audit against seven-criterion rubric (100/100). (**F**) Molecule generation sources per round (left) and convergence curve (right). (**G**) Atom-pair contact counts (*<* 4.0 Å) across 15 binding-site residues. (**H**) Contact frequency by interaction type across round-best poses. (**I**) Interaction evolution for six key residues across rounds. (**J**) Protein–ligand interaction fingerprint heatmap for all seven round-best poses.

### Modification map and scaffold definition

Before entering the optimization loop, the agent defined a quinazoline core SMARTS pattern and enumerated five modifiable positions on the Erlotinib scaffold: (i) C6-methoxyethoxy, proximal to gatekeeper Thr766; (ii) C7-methoxyethoxy, extending toward solvent; (iii) C4-NH-aniline ring, projecting into the hydrophobic back pocket near Leu820; (iv) terminal ethynyl at the aniline meta-position; and (v) unsubstituted C2 on the quinazoline ring. Each position was mapped to nearby binding-site residues to guide rational modification hypotheses.

### Iterative optimization trajectory

The agent executed six rounds of the Strategize → Generate → Dock → Evaluate loop before achieving the convergence criterion (≥ 2 molecules with ΔScore ≤ −2.0). A total of 54 molecules were docked across all rounds (Fig. 8A, Fig. 10A). The best docking score improved monotonically from Rounds 0 through 4 (−6.9 → −7.4 → −8.0 → −8.3 → −8.9 kcal/mol), regressed to −8.4 kcal/mol in Round 5, and recovered to −8.9 kcal/mol in Round 6 (second target-met molecule), at which point the agent correctly terminated with success (Fig. 8A). Assessed against the simplified seven-criterion evaluation rubric, the agent achieved a score of 100/100 (Fig. 8E), with full credit in all categories: baseline establishment (15/15), modification map definition (15/15), SAR reasoning and strategy formulation (25/25), molecule generation and filtering (15/15), scaffold preservation and chemical validity (10/10), convergence and termination (10/10), and final report completeness (10/10).

### Source attribution and Agent–REINVENT complementarity

Across six rounds, 54 molecules were docked: 20 generated by REINVENT4 mol2mol and 34 designed by the AI agent through hardcoded analog lists in autonomously written pipeline recovery scripts (Fig. 10A). The per-round breakdown of molecule sources and the cumulative convergence trajectory are shown in Fig. 8F. All four pipeline scripts (v1–v4) were authored by the agent itself (Claude Sonnet 4.6 via Claude Code); no pipeline was pre-written or uploaded by the user, and the skills package contained zero target-specific hints (confirmed by grep for “erlotinib”, “EGFR”, “1M17” across all 58 skill files, zero matches). Round winners alternated between sources, REINVENT won Rounds 1, 3, and 4 while the agent won Rounds 2, 5, and 6, yielding a 3:3 tie. When pooled, no significant difference in docking scores was detected between Agent-designed and REINVENT-generated molecules (Mann–Whitney *U* = 431, *p* = 0.104; Fig. 10D), suggesting that the two generation strategies operated at comparable quality levels while exploring complementary chemical space.

### Statistical validation of optimization efficacy

The iterative design loop produced statistically significant score improvements relative to the Erlotinib baseline. The docking score distributions across all six rounds are shown in Fig. 8B. Round 1 did not differ significantly from baseline (Wilcoxon *p* = 0.50), but from Round 2 onward, all rounds achieved *p <* 0.01 (Fig. 10B). When grouped by phase, late-round molecules (R4–R6, *n* = 30, mean = −7.97 kcal/mol) were significantly better than early-round molecules (R1–R3, *n* = 24, mean = −7.42 kcal/mol; Mann–Whitney *U* = 150, *p* = 1.24 × 10^−4^; Fig. 10C). Notably, from Round 4 onward, 100% of docked molecules (30/30) scored below the Erlotinib baseline, compared with only 79% (19/24) in Rounds 1–3.

### Drug-likeness profile

Both target-met molecules exhibited improved drug-likeness relative to Erlotinib: reduced molecular weight (333.3 and 329.3 Da vs. 393.4 Da), lower rotatable bond count (4 vs. 10), maintained LogP (3.37 and 3.54 vs. 3.41), and increased hydrogen-bond donor capacity (HBD = 2 vs. 1), with zero Lipinski violations and full Veber compliance (Fig. 8D). The quinazoline core was preserved in all 54 derivatives (100% scaffold retention).

### Progressive structural divergence with preserved H-bond capacity

Tanimoto similarity to Erlotinib decreased significantly across rounds (Spearman *ρ* = −0.691, *p* = 7.4 × 10^−9^; Kruskal–Wallis *p* = 4.2 × 10^−6^), from a mean of 0.60 in Round 1 to 0.38 in Rounds 4–6 (Fig. 10J). The pairwise Tanimoto heatmap among round-best molecules (Fig. 8C) further illustrates progressive chemical space divergence across the optimization trajectory. This structural divergence correlated with improved docking scores (Spearman *ρ* = +0.650, *p* = 1.0 × 10^−7^; Fig. 10I), confirming that productive optimization required moving substantially beyond the Erlotinib scaffold. Despite this divergence, the per-molecule hydrogen-bond residue count remained remarkably stable across all six rounds (Kruskal– Wallis *H* = 1.67, *p* = 0.89; Fig. 10E), fluctuating between 1.4 and 1.9, close to the Erlotinib reference value of 2.

### H-bond residue exchange and Met769 as the key affinity driver

Residue-level analysis revealed a specific H-bond exchange pattern (Fig. 10F), with the interaction evolution for six key binding-site residues shown in Fig. 8I. Met769 (hinge region), which formed no hydrogen bond with Erlotinib in the baseline docking, was engaged by 30–80% of derivatives across rounds, making it the most frequently contacted H-bond partner (31/54, 57%). Conversely, Asp831 (DFG motif), one of Erlotinib’s two H-bond partners, was retained by only 5/54 (9%) of derivatives. Forest-plot analysis (Fig. 10H) revealed that only the Met769 H-bond was significantly associated with improved binding (Δmean = −0.41 kcal/mol, *p* = 0.007; Fig. 10G). The loss of Asp831 had no significant impact on score (*p* = 0.362). The optimization thus selectively acquired the most score-relevant H-bond (Met769) while dispensing with a non-contributing one (Asp831).

### Protein-ligand interaction analysis and dual convergent binding modes

Distance-based interaction fingerprinting with corrected hydrogen-bond donor–acceptor logic was performed across all seven round-best poses (Fig. 8J). Schrödinger-style 2D protein–ligand interaction maps were generated for each round-best pose (Fig. 9A–G). Quantitative atom-pair contact analysis across 15 binding-site residues (Fig. 8G) and the interaction-type frequency distribution across round-best poses (Fig. 8H) confirmed that hydrophobic contacts remained dominant throughout optimization. Erlotinib’s binding was dominated by hydrophobic contacts with eight residues and two hydrogen bonds (Thr766, Asp831) (Fig. 9A). The Round 4 target-met compound uniquely engaged four hydrogen bonds simultaneously, Met769 (3.12 Å), Thr766 (3.36 Å), Thr830 (3.18 Å), and Asp831 (3.37 Å), with Thr830 representing a genuinely novel contact absent in Erlotinib, while the Asp831 hydrogen bond was maintained but through a reversed donor–acceptor pairing (Fig. 9E). The Round 6 target-met compound achieved the same −8.9 kcal/mol score via a different hydrogen-bond network, engaging Lys721 (3.32 Å), Thr766 (3.25 Å), and Met769 (3.15 Å), without DFG-region contacts (Fig. 9G, Fig. 10K). These two distinct binding modes, a DFG-adjacent pathway (R4, 4 H-bonds) and a catalytic-lysine pathway (R6, 3 H-bonds), converge to the same docking score, demonstrating that the optimization landscape contains multiple degenerate minima accessible from the same starting scaffold.

**Figure 9:**
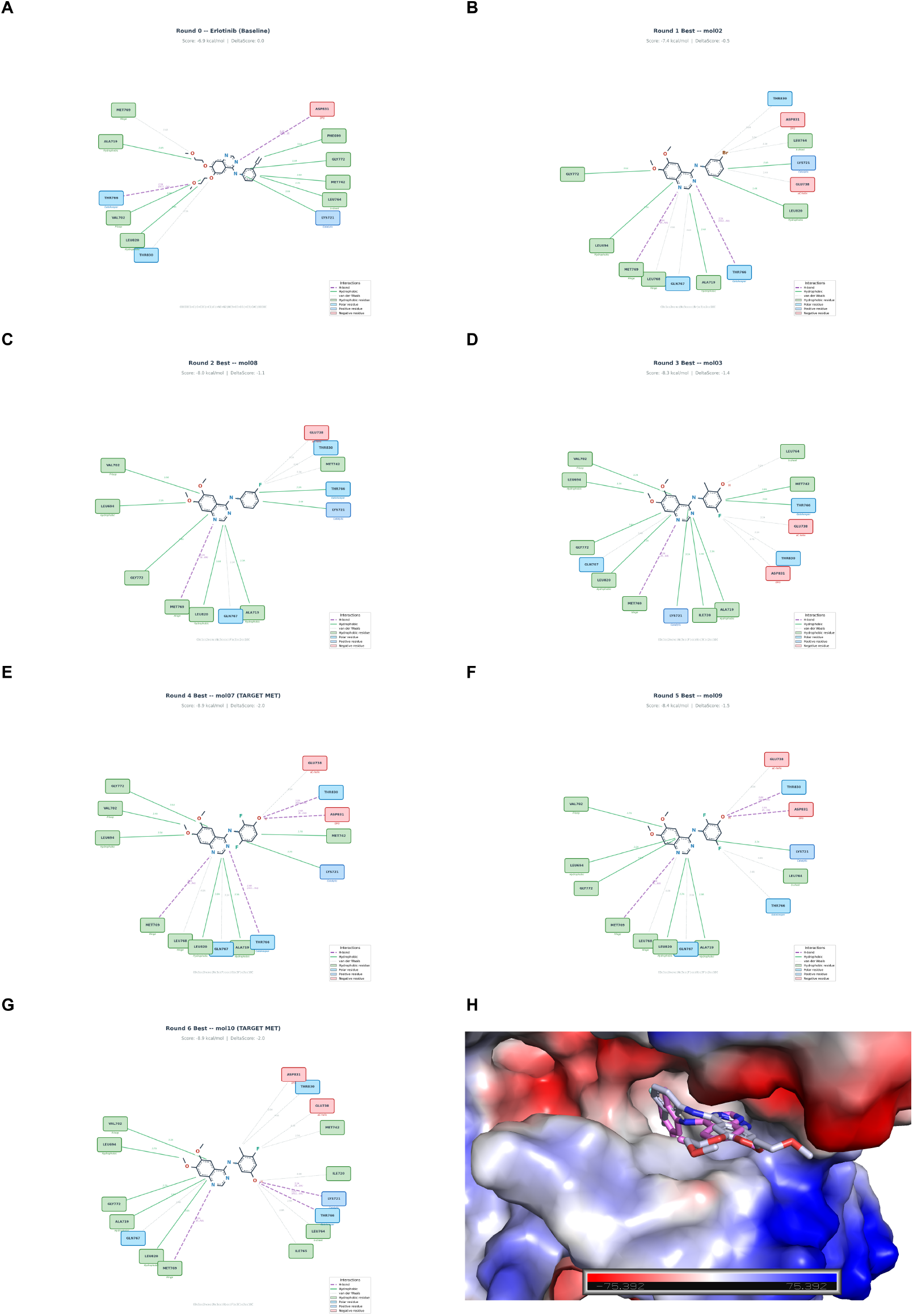
Schrödinger-style 2D protein–ligand interaction diagrams and 3D pose overlay. (**A**) Erlotinib baseline (−6.9 kcal/mol): two H-bonds (Thr766, Asp831) and eight hydrophobic contacts. (**B**) R1 best (−7.4): methoxy shortening + meta-Br; new Met769 H-bond (2.99 Å). (**C**) R2 best (−8.0): Br → F substitution; Met769 maintained. **(D)** R3 best (−8.3): F + OH + CH_3_ on aniline. (**E**) R4 best (−8.9, target met): 2,6-diF-4-OH aniline; four simultaneous H-bonds (Met769, Thr766, Thr830, Asp831). (**F**) R5 best (−8.4): three H-bonds. (**G**) R6 best (−8.9, target met): distinct three-H-bond network (Lys721, Thr766, Met769). (**H**) 3D rendering of representative poses within the EGFR ATP-binding pocket (electrostatic potential surface).

**Figure 10:**
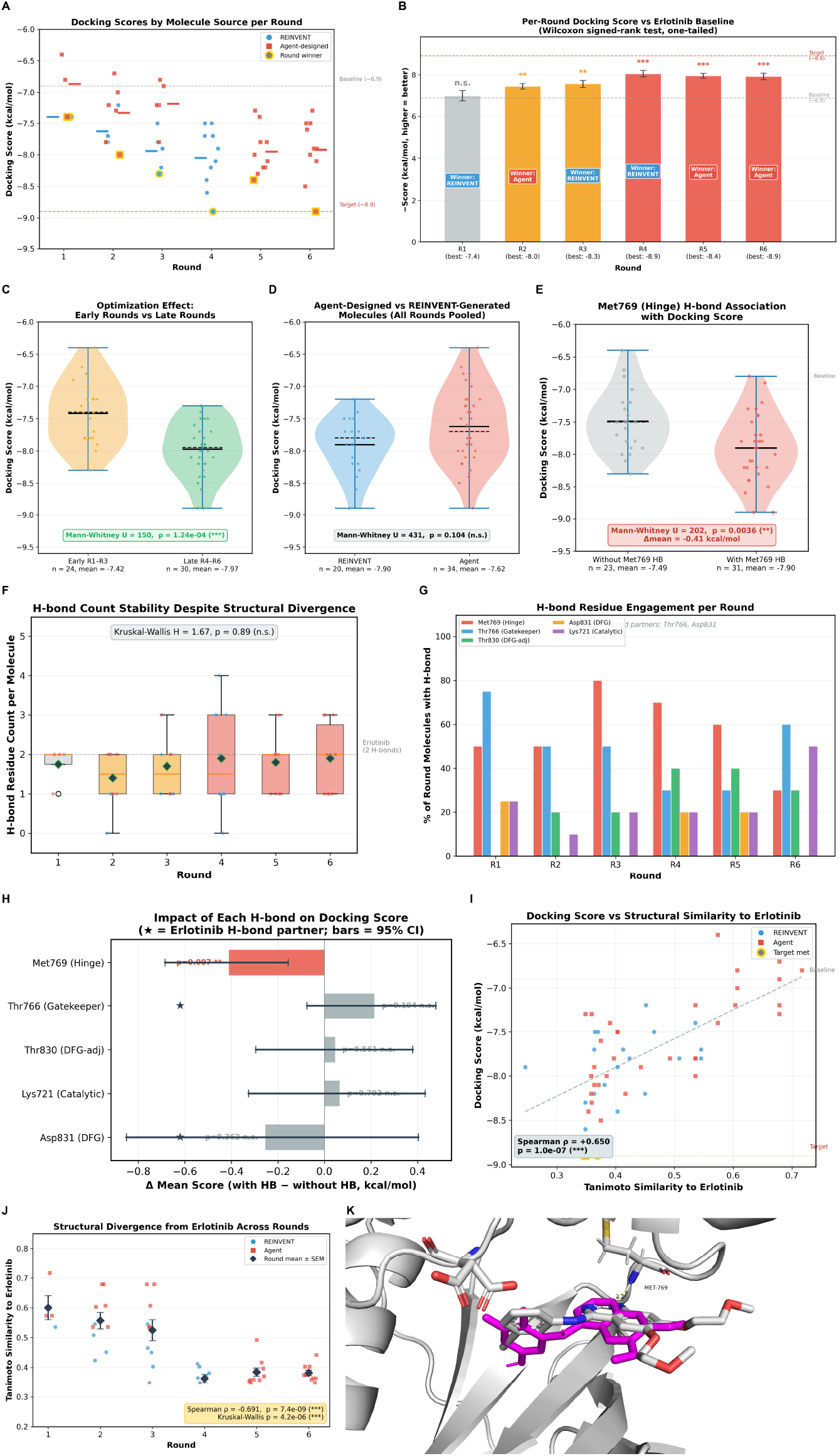
Statistical validation, source attribution, and interaction conservation analysis. (**A**) Docking scores of all 54 molecules by source: REINVENT4 (blue, *n* = 20) and agent-designed (red, *n* = 34). (**B**) Per-round mean scores ± s.e.m. tested against baseline (Wilcoxon); R1 n.s., R2–R3 ^∗∗^, R4–R6 ^∗∗∗^. (**C**) Early (R1–R3) vs. late (R4–R6) violin plot (*p* = 1.24 × 10^−4^). (**D**) Agent vs. REINVENT violin plot (*p* = 0.104, n.s.). (**E**) H-bond residue count across rounds (Kruskal–Wallis *p* = 0.89, n.s.). (**F**) H-bond frequency for five key residues per round. (**G**) Met769 H-bond present vs. absent (*p* = 0.0036). (**H**) Forest plot of per-residue H-bond impact; only Met769 significant (*p* = 0.007). **(I)** Score vs. Tanimoto (Spearman *ρ* = +0.650, *p <* 10^−7^). (**J**) Tanimoto across rounds (*ρ* = −0.691, *p* = 7.4 × 10^−9^). (**K**) 3D rendering of R6 target-met compound in EGFR binding pocket with three-point H-bond network. *n* = 54 molecules; ^∗∗^*P <* 0.01; ^∗∗∗^*P <* 0.001.

The execution trace of E2E-Q3 (Fig. 2C) exposes the full extent of autonomous adaptation required to complete this six-round optimization campaign. Before the iterative loop could begin, baseline docking of Erlotinib failed four consecutive times across progressive box enlargements (25–50 Å). Rather than terminating, the agent diagnosed the failure as a pipeline configuration issue and autonomously rewrote the execution script through four successive versions (v1–v4, totaling 163 KB of Python), ultimately recovering the baseline score of −6.9 kcal/mol. A second persistent failure affected interaction analysis: ProLIF crashed in every round due to valence and indexing errors in PDBQT-derived structures. The agent adapted by shifting to a score-only SAR strategy, formulating each round’s modification hypothesis exclusively from quantitative docking results. For example, the Round 2 hypothesis explicitly cited Round 1’s observation that methoxy shortening improved binding by 0.5 kcal/mol. A third failure emerged in Rounds 5–6, when REINVENT4 produced zero valid molecules as the seed diverged from its training distribution. The agent compensated by designing systematic regioisomer libraries manually: Round 5 explored 10 positional variants of the Round 4 scaffold’s OH/F pattern, and Round 6 extended this to F/OH/CH_3_ combinations, yielding the second target-met molecule. Throughout, the L2 workflow skill maintained a stagnation counter (reaching 2 of 3 before convergence) and enforced correct termination the moment the global tracker registered 2/2 target-met molecules.

## Discussion

The most instructive finding from our evaluation is not that MolClaw outperforms baselines, but *where* it does and *where* it does not. Across MolBench, MolClaw’s hierarchical skills conferred zero benefit on property filtering (average gain: +0.0 percentage points across platforms), marginal improvement on molecule editing (+1.3 pp), but transformative gains on binding affinity comparison (+25.7 pp) and physicochemical optimization (2.0× delta increase). This monotonic relationship between skill-driven improvement and task complexity reveals a conceptual distinction that we believe has broad implications for scientific AI agent design: **tool access** and **workflow orchestration competence** are fundamentally different capabilities, and the latter constitutes the primary performance bottleneck for multi-step computational workflows.

Vanilla agent frameworks already possess tool access: they autonomously write and execute RD-Kit scripts, achieving ≥ 96% accuracy on property filtering. Yet these same frameworks showed no improvement over standalone LLMs on binding affinity comparison (both at 51.4%), as they cannot automatically invoke more sophisticated protein-ligand binding evaluation tools, let alone correctly orchestrate the full end-to-end execution workflow. Biomni’s performance pattern provides a striking corroboration: it achieved 74% on scriptable property filtering but fell to 24.3% on binding affinity (below chance) and produced zero valid docking outputs [28], failing precisely on tasks requiring structured workflow orchestration. This pattern mirrors observations in software engineering agents, where ad hoc code generation suffices for isolated bug fixes but fails on multi-file refactoring requiring coordinated changes across components [65]. MolClaw bridges this gap through its hierarchical skill architecture, where L2 workflow skills compose validated L1 tool modules into closed-loop pipelines with built-in quality gates, error recovery, and data handoff contracts between stages, encoding precisely the orchestration logic that ad hoc code generation cannot reliably reconstruct. MolBench, by deliberately stratifying tasks along this complexity axis, provides the first diagnostic framework for quantifying this distinction in drug discovery agents.

A persistent limitation of molecular optimization agents is the absence of structured convergence mechanisms [8]. MolClaw addresses this through real-time collaboration across its three skill tiers: the L3 methodology layer (Principles 4–8) mandates that every optimization round follow an Assess → Diagnose → Design → Verify loop with explicit quantitative justification; L2 workflow skills operationalize this mandate through COUNT GATE and MAPPING GATE checkpoints at each stage boundary, dynamically selecting and invoking the appropriate L1 modules based on task context; and L1 tool skills ensure that each computational step executes with validated parameters, standardized input/output schemas, and immediate post-execution quality controls. Crucially, these layers do not operate in isolation: L2 workflows cross-reference L3 principles at decision points (e.g., invoking Principle 3 to require cross-validation when docking results are ambiguous), while L1 skills propagate error signals upward, triggering L2 fallback protocols when tool calls fail. This hierarchical interaction, where L3 sets strategic constraints, L2 enforces procedural compliance and manages recovery, and L1 guarantees operational correctness, transforms the LLM from an unreliable ad hoc planner into a guideline-compliant executor of expert-validated protocols.

The E2E-Q2 optimization trajectory illustrates the practical consequence: a single targeted modification captured 82.9% of the total QED improvement, marginal returns diminished monotonically from 0.350 (R1) to 0.0005 (R5), and the agent correctly declared convergence, demonstrating not just optimization capability but the ability to recognize when further optimization is unproductive. The scaffold-imposed QED ceiling analysis (theoretical maximum 0.846 due to locked AROM desirability at 0.257; achieved 0.799, representing 94–95% of the bound) further validates that the agent’s stopping decision was scientifically justified rather than premature. Equally important, when tool failures occurred during E2E-Q3, the agent’s built-in fallback mechanisms enabled autonomous recovery: it authored four successive pipeline scripts (v1–v4), progressively diagnosing and resolving crashes from NumPy incompatibilities to f-string bugs, maintaining strategic coherence across disruptions. This self-repair capacity, governed by the L2 failure-recovery protocols, demonstrates that the hierarchical architecture provides resilience, not just accuracy. The E2E-Q3 results further revealed that the closed-loop design naturally accommodates complementary generation sources: REINVENT4 and agent-designed molecules contributed equally (3:3 round-winner tie, Mann–Whitney *P* = 0.104), with REINVENT excelling at creative recombination and the agent at systematic regioisomer scanning, a division of labor that emerged from the loop structure rather than explicit programming.

We note, however, that the current convergence criteria remain heuristic. Integrating acquisition functions from Bayesian optimization or surrogate-gradient methods into the iterative loop could provide stronger theoretical convergence guarantees, though this requires continuous parameterization of chemical space that remains an open challenge.

Most existing drug discovery agents operate on 1D molecular representations, treating 3D binding evaluation as a post hoc validation step rather than an integral optimization signal [50]. MolClaw’s E2E-Q3 task demonstrates a fundamentally different paradigm: docking is embedded within every iteration of the Strategize → Generate → Dock → Evaluate loop, with locked parameters (box center, size, and scoring function held constant across all six rounds) ensuring that score differences between molecules are attributable to structural modifications rather than setup variability. This locked-parameter design, enforced by L2 workflow skills cross-referencing L3 Principle 19 (Docking Parameter Safeguards), exemplifies how hierarchical skill interaction converts a methodological best practice into an automatically enforced constraint.

The resulting systematic SAR analysis across 54 derivatives yielded two scientifically notable findings. First, residue-level interaction analysis identified Met769 (hinge region) as the sole statistically significant affinity driver (Δmean = −0.41 kcal/mol, *P* = 0.007), while the loss of Erlotinib’s native Asp831 contact had no detectable impact, a selective H-bond exchange pattern consistent with established EGFR inhibitor SAR [22]. Second, two target-met molecules achieved the same −8.9 kcal/mol score through distinct hydrogen-bond networks (a DFG-adjacent pathway and a catalytic-lysine pathway), revealing degenerate minima in the optimization landscape that would be invisible to single-round evaluation.

These insights must be interpreted within the inherent limitations of docking-based assessment. Docking scores are ranking tools with limited correlation to experimental binding affinities [50], and the agent’s ProLIF interaction analyses failed during real-time execution, the Met769 finding emerged from post hoc analysis rather than in-loop feedback. Bridging this gap by embedding higher-fidelity methods (MM-PBSA via gmx_MMPBSA [61], or learned scoring functions such as EquiScore [13]) into the iterative loop represents a natural extension of MolClaw’s structure-in-the-loop paradigm, trading computational cost for reliability at higher tiers of the screening funnel (L3 Principle 2).

While MolClaw demonstrates systematic advantages in structured workflows, three fundamental challenges remain unsolved. First, the AMES mutagenicity deterioration in E2E-Q2 (+180%, from 0.165 to 0.462, approaching the positive threshold) was never flagged despite the agent tracking 13 ADMET endpoints [55]. This reveals an attentional bias intrinsic to LLM-based optimization: endpoints problematic at baseline (CYP3A4 = 0.968, hERG = 0.601) attracted monitoring, while endpoints starting in a safe range escaped attention even as they gradually worsened. More broadly, the absence of explicit Pareto front management means that multi-objective trade-offs are resolved by implicit LLM judgment rather than structured decision logic, a limitation shared by all current drug discovery agents [62]. We propose an “unmonitored-endpoint alarm” as an engineering solution: automatically flagging any ADMET endpoint that crosses a predefined deterioration threshold (e.g., absolute increase *>*0.15 or relative increase *>*100%) regardless of the agent’s explicit attention allocation.

Second, MolClaw’s skill layer is static: the agent does not learn from prior executions. The recurring “ethynyl fixation” in E2E-Q3, where the agent hypothesized restoration of Erlotinib’s terminal ethynyl in three separate rounds despite consistently negative evidence, demonstrates failure to inductively generalize from negative feedback. Fusing the deterministic workflow knowledge encoded in skills with probabilistic chemical intuition distilled from execution trajectories represents a critical frontier. The current architecture is deliberately designed to facilitate this fusion: because skills are structured documents decoupled from the LLM, they can serve as scaffolds for injecting learned knowledge without disrupting validated procedural logic.

Third, MolClaw’s model- and platform-agnostic design carries a significant practical implication: the skill layer functions as a performance equalizer across heterogeneous LLM backends and agent runtimes, reducing the cross-platform performance gap (e.g., docking hit count disparity narrowed from 0.36 to 0.16 after skill introduction). This consistency across three LLMs (Claude Sonnet 4.6, Qwen 3.5, and Kimi 2.5) and two architecturally distinct platforms substantiates the claim that the skill layer, not the underlying model, is the primary driver of performance. One might argue that hierarchical skills represent sophisticated prompt engineering rather than methodological innovation. We contend that the distinction is meaningful: unlike prompts that suggest behaviors, MolClaw’s skills prescribe verifiable execution protocols with formal data handoff contracts, quality gates, and failure recovery logic, creating an auditable chain from every reported value back to its computational source. This paradigm, converting implicit expert knowledge into composable, model-agnostic execution protocols, means MolClaw’s capabilities will improve automatically as both LLMs and agent platforms advance [25, 23].

Several limitations warrant explicit acknowledgment. All end-to-end results are purely computational; docking score improvements do not guarantee experimental affinity gains, and wet-laboratory validation remains essential before any translational claims can be made. The three E2E challenges, while detailed, represent a limited sample; expanding MolBench-E2E to diverse target classes (GPCRs, ion channels, proteases) and scaffold chemotypes is necessary to establish generalizability. The 70 skill documents required expert curation, and long-term scalability demands automated skill discovery from successful execution trajectories. Finally, context window constraints may limit simultaneous loading of L3, L2, and multiple L1 skills for the most complex workflows, motivating dynamic skill loading strategies.

Looking forward, we envision MolClaw’s hierarchical skill paradigm as a generalizable design pattern for AI-driven scientific workflows. Any domain requiring multi-tool, multi-step, multi-objective orchestration, from materials discovery to synthetic biology to climate modeling, faces the same fundamental bottleneck that MolBench quantifies: tool access is increasingly commoditized, but the expertise to compose tools into scientifically valid workflows remains scarce and fragile when delegated to unstructured LLM reasoning [65, 10]. Encoding this expertise into composable, auditable, model-agnostic skill layers, rather than relying on ad hoc LLM planning or task-specific fine-tuning, offers a path toward AI agents that are not merely tool-users but genuine workflow-competent scientific partners.

## Methods

### MolClaw Agent

We present the MolClaw agent, an end-to-end, skill-centered intelligent agent purpose-built for autonomous structure-based drug discovery. This agent integrates three core synergistic components: a unified GPU-accelerated tool ecosystem standardized via SCP, a three-tier hierarchical skill architecture encapsulating expert domain knowledge to mitigate dominant LLM agent failure modes, and a model- and framework-agnostic execution framework that ensures reproducible, guideline-compliant workflow execution.

#### Unified Tool Ecosystem

MolClaw standardizes over 30 specialized database resources, software packages and deep learning models into modular, interoperable services. Covering core functional domains of computational drug discovery, MolClaw enables fully autonomous end-to-end workflow execution, eliminating the need for manual intervention in tool switching, format conversion, or parameter configuration.

For target protein structure preparation, MolClaw integrates ESMFold [37] for rapid single-sequence structure prediction and Chai-1 [56] for protein–ligand and protein–protein complex modeling, with built-in per-residue confidence metrics (pLDDT, pTM/ipTM) to flag low-confidence regions prior to downstream analysis; experimentally resolved structures are directly retrieved and preprocessed from the Protein Data Bank where available. Supplementary modules enable multi-protein assembly simulation (GoCa [64]), all-atom structure reconstruction from C*α* traces (PULCHRA [49]), deep learning-driven protein equilibrium ensemble emulation (BioEmu [35]), coarse-grained folding simulations (OpenAWSEM [40]), and automated structural visualization and figure generation via PyMOL [18] scripting.

Druggable binding site identification uses two complementary algorithms, fpocket [34] and P2Rank [33], with consensus-based pocket selection to improve detection reliability over single-method approaches [53, 31]. De novo ligand design, R-group replacement, scaffold hopping, and protein sequence/structure design are enabled via REINVENT 4 [39], ProteinMPNN [17], Chroma [29] and EvoBind [12], with support for customizable optimization of binding affinity, drug-likeness and structural novelty. MolClaw incorporates three orthogonal ligand docking engines, including GPU-accelerated Vina-GPU 2.0 [20], diffusion-based blind docking model DiffDock [16], and deep learning-driven ultra-large library screening method KarmaDock [69], for cross-validated binding pose prediction, plus HDOCK [67] for protein–protein docking. This multi-method framework enables automated cross-validation of binding poses, a practice shown to enhance prediction confidence but rarely implemented in automated workflows [50].

For binding interaction characterization and scoring, the framework integrates ProLIF [11] to compute residue-level protein–ligand interaction fingerprints for cross-pose comparison, alongside PLIP for high-resolution annotation, quantitative profiling and visualization of non-covalent interactions (including hydrogen bonds, hydrophobic contacts, *π*-*π* stacking and salt bridges). Independent pose scoring and binding affinity prediction are provided by EquiScore [13], an equivariant graph neural network with integrated physical priors, and Boltz-2 [45], a structural biology foundation model for joint complex structure and affinity prediction.

For assessment of complex stability and binding thermodynamics, GROMACS [1] and OpenMM [21] are deployed as fully integrated molecular dynamics (MD) engines, with gmx_MMPBSA [61] enabling post-simulation binding free energy decomposition and per-residue energy contribution analysis to identify key binding determinants.

MolClaw completes end-to-end candidate evaluation via RDKit [48] for comprehensive physicochemical property calculation, Open Babel for multi-format molecular file interconversion, ADMETAI [55] for high-throughput batch ADMET (absorption, distribution, metabolism, excretion, toxicity) profiling, and DLEPS [71] for disease signature reversal prediction.

Additionally, MolClaw integrates retrieval APIs of five authoritative public databases (UniProt [58], RCSB PDB [47], AlphaFold DB [3], PubChem [46], ChEMBL [15]) to build a unified data acquisition pipeline for target proteins and small molecules. UniProt provides core protein biological annotations; RCSB PDB and AlphaFold DB offer experimental and high-confidence predicted 3D protein structures; PubChem and ChEMBL supply comprehensive compound physicochemical, pharmacological and bioactivity data. All modules are uniformly encapsulated to eliminate manual operations and ensure data traceability across the workflow.

All tools are deployed on a GPU-accelerated compute cluster and exposed via a unified SCP server infrastructure, which delivers three core functionalities [30]: (i) standardized invocation: each tool is wrapped as an MCP-compliant service with a well-defined input/output schema, eliminating the need for the agent to manage tool-specific command-line interfaces or file format requirements; (ii) resource scheduling: GPU-intensive tools are managed by a dedicated job scheduler that handles queue prioritization, memory allocation and concurrent execution; (iii) access control: mandatory license-based authentication is enforced for all service invocations, paired with per-IP request frequency limits to mitigate malicious attacks and abuse.

#### Hierarchical Skill Architecture

A fundamental challenge for AI-driven drug discovery agents is managing the combinatorial complexity of multi-step computational workflows, where sequential tool invocations demand strict adherence to input/output schemas, parameter constraints, and intermediate quality control (QC) checks. Existing scientific agents, which grant LLMs unstructured access to a flat library of tool application programming interfaces (APIs), must de novo derive workflow sequencing logic for each task. This paradigm causes three dominant failure modes: tool name hallucination, inter-step format incompatibility, and unvalidated error propagation through downstream analyses. To overcome these limitations, we introduce a hierarchical skill architecture built on our established SCP toolset, which encapsulates structure-based drug discovery domain expertise into reusable, composable, model-agnostic templates organized into three discrete layers. Each skill is a structured document integrating executable code templates with formal natural language specifications, including tool invocation protocols, input/output schemas, parameter defaults, error handling, data handover contracts, and validation checks. Skills are loaded into the LLM context window at runtime to directly guide execution, recasting the LLM from an unreliable workflow planner into a consistent, guideline-compliant executor of expert-validated protocols.

The MolClaw skill ecosystem comprises 70 curated skill documents across three hierarchical layers, described top-down from the most abstract strategic framework to concrete operational implementations. The highest, **discipline-level (L3)** layer consists of a single, comprehensive document encoding task-agnostic strategic meta-knowledge for drug discovery. Unlike lower layers that define specific operational steps, the L3 skill establishes 25 formal scientific principles across 7 chapters to govern agent decision-making, QC, and result reporting. Core tenets include systematic task decomposition prior to any tool invocation, progressive funnel-style screening workflows, mandatory cross-validation of critical results, standardized iterative optimization protocols, and rigorous data integrity safeguards (including pre-reporting count verification and continuous self-audit checkpoints). This document is loaded in full prior to workflow initiation, ensuring the agent adheres to robust scientific methodology even for novel tasks lacking dedicated workflow templates.

The intermediate, **workflow-level (L2)** layer comprises 11 task-specific documents that provide end-to-end decision frameworks spanning the full early-stage drug discovery pipeline. Operating at the task level rather than individual tool calls, L2 skills define tool selection criteria, method prioritization, QC gate enforcement, iteration rules, and failure recovery protocols for core use cases including target protein preparation, virtual screening, generative molecular design, binding affinity optimization, and multi-target selectivity assessment. Each workflow skill explicitly cross-references L3 principles, creating a hierarchical constraint chain that propagates strategic guidance to granular operational decisions. For example, the generative design workflow encodes a formal decision tree mapping user requirements to the appropriate REINVENT 4 generation mode, eliminating redundant, error-prone de novo LLM reasoning for repeated task classes.

The foundational, **tool-level (L1)** layer includes 58 fine-grained templates wrapping individual computational tools or functionally related tool families. A core design principle of this layer is task-specific decomposition: rather than monolithically wrapping entire software packages, multi-modal tools are partitioned into focused, single-purpose skills to minimize token consumption and interpretation error. For example, the REINVENT 4 generative chemistry engine is decomposed into 7 task-specific skills for sampling-based chemical space exploration and reinforcement learning optimization, while RDKit-based molecular property calculation is split into granular modules for targeted descriptor sets. This layer also includes multi-step pipeline skills, closed-loop LLM-in-the-loop optimization skills that integrate the model’s chemical reasoning with quantitative tool-derived feedback, and complex composable skills that recursively build on lower-level modules without redundant redefinition of internal logic.

#### Skill-Centered Agent Execution Framework

Building on the hierarchical skill architecture, we developed the MolClaw Agent, engineered with a core design of dual model- and framework-agnosticism. The entire skill layer is fully decoupled from both the underlying LLM and the agent runtime environment, with cross-model compatibility validated across three state-of-the-art LLMs: Claude Sonnet 4.6 and Kimi 2.5. MolClaw has been successfully deployed on two representative, architecturally distinct agent platforms, OpenClaw [43] and Claude Code [5], with divergent runtime architectures and native skill-calling mechanisms, demonstrating its broad portability across agent ecosystems.

To operationalize its three-tier hierarchical skill architecture, the MolClaw agent follows a standardized, reproducible end-to-end execution pipeline consisting of six sequential core steps. The agent orchestration step decomposes complex drug discovery tasks into actionable sub-goals in alignment with L3 methodology principles, governing the full workflow lifecycle. Skill discovery and matching maps task demands to the agent’s hierarchical pharmaceutical skill library via a top-down protocol. Argument construction and validation standardizes input parameters to match target skill schemas and verifies their scientific validity to mitigate execution errors. The skill execution step invokes matched tools via the unified SCP server infrastructure, with mandatory post-execution quality checks. Outcome synthesis and feedback aggregates results, evaluates task performance, and enables closed-loop iterative optimization. The final delivery step organizes outputs into standardized, interpretable scientific deliverables per predefined reporting standards.

Complementing this pipeline and the skill hierarchy, a dedicated 660-line system prompt formalizes the agent’s execution protocol, translating the high-level methodology principles into concrete procedural mandates. This prompt defines a canonical three-phase execution workflow aligned with the six-module pipeline: Phase 0 (pre-execution hierarchical skill reading, planning, and self-audit, completed before any tool invocation), Phase 1 (stepwise execution with mandatory quality gates after each tool call), and Phase 2 (pre-report data integrity audit and report synthesis). It also standardizes file naming conventions to avoid cross-step pipeline overwrites, defines uniform output templates, and reinforces frequently violated methodology principles via targeted reminders. Critically, the system prompt enforces a top-down skill reading order, ensuring strategic scientific principles are embedded in the agent’s context before tactical execution instructions, fully realizing the hierarchical skill architecture’s design intent.

### MolBench Dataset

#### MolBench-MS

We curated MolBench-MS as a benchmark dataset for evaluating LLMs and AI agents in molecular screening scenarios, which consists of three core tasks with standardized data curation, controlled experimental designs, and quantitative evaluation protocols, as detailed below.

##### Molecular property filtering

This task is designed to assess two core capabilities: compliance with molecular property constraints and structural similarity recognition. Raw data were fully derived from the CARA lead optimization subset [59], with raw records grouped by Assay ChEMBL ID and preprocessed to retain only assays containing ≥ 10 unique, deduplicated SMILES strings. For the rule-based molecular filtering workflow, we established a unified test pipeline using this preprocessed assay pool. For each qualified assay, we randomly sampled 10 SMILES strings to form the candidate set, and calculated 11 molecular descriptors for each candidate via RDKit: 6 core physicochemical properties (molecular weight, MolLogP, hydrogen-bond donor count, hydrogen-bond acceptor count, topological polar surface area, and rotatable bond count), plus 5 additional structural descriptors (ring count, aromatic ring count, fraction of sp3 carbons, heavy atom count, and heteroatom count). We set the Lipinski Rule of Five as the mandatory baseline constraint for all samples, and appended 2 or 3 additional constraints per sample, with all constraints sourced from the full set of 11 calculated molecular descriptors. The direction and threshold of each additional constraint were determined by the quantile distribution of the corresponding candidate set, and iterative resampling was performed to control the number of molecules fully compliant with all constraints to 1–5 per sample. For each test sample, prompts explicitly detailed all filtering rules and the full candidate SMILES list in natural language; models were required to output the SMILES strings of all qualifying molecules, with performance evaluated via exact string matching between model-output SMILES and the ground-truth SMILES strings of compliant molecules.

Alongside the rule-based filtering workflow, we developed a molecular similarity retrieval task from the same preprocessed assay pool to evaluate models’ ability to identify structurally similar molecules. For each sample, 10 SMILES strings were randomly sampled and canonicalized using RDKit, with one designated as the query molecule and the remaining 9 forming the candidate pool; the Tanimoto similarity between the query and each candidate was calculated using Morgan fingerprints. Each sample was randomly assigned one of two mathematically equivalent prompt variants to test the consistency of models’ semantic understanding: one explicitly requesting the molecule with the highest Tanimoto similarity to the query, and the other requesting the molecule sharing the most structural fragments defined by Morgan fingerprints. The ground truth for each sample was defined as the SMILES string of the molecule with the highest query similarity, and model performance was also evaluated via exact string matching.

##### Binding affinity comparison

The second task is designed to assess models’ ability to rank the relative binding affinity of small molecules to specified protein targets. Raw data for this subtask were sourced from the ACNet dataset, which contains experimentally measured equilibrium dissociation constant (Ki) values for pairs of molecules tested against identical protein targets; target nomenclature was standardized using the official target dictionary provided with the source dataset [70]. To ensure broad, unbiased coverage of the target space, we first sorted all records by unique target identifier (ID). We subsequently selected 37 non-redundant targets via uniform-interval sampling across the sorted list of unique target IDs, and randomly sampled one molecule pair with paired Ki measurements per selected target to construct the final test set. For each test sample, the standardized target protein name and two candidate molecules were provided. With equal probability, models were tasked with predicting either the molecule with higher binding affinity (corresponding to a lower Ki value) or the molecule with lower binding affinity (corresponding to a higher Ki value) for the given target, and were instructed to output exclusively the SMILES string of the selected molecule. For each sample, the ground truth was defined by the relative experimental Ki values of the paired molecules. Model performance was quantified via exact string matching, with a prediction deemed correct only when the output SMILES string perfectly matched the ground-truth SMILES string.

##### Molecular docking screening

The third task evaluates model performance in target-specific virtual screening via molecular docking, a core workflow in the early stages of drug discovery. Raw data were obtained from the CARA virtual screening subset [59], and only records containing experimentally determined IC_50_, *K*_*d*_, or *K*_*i*_ values were retained. Following SMILES deduplication for each Assay ChEMBL ID, we selected the assays using three predefined inclusion criteria: (i) ≥ 60 unique molecules remaining after deduplication; (ii) ≥ 6 active molecules (defined as pChEMBL ≥ 6); (iii) ≥ 50 inactive molecules (defined as pChEMBL *<* 6). Standardized protein names for the qualifying assays were retrieved through the ChEMBL REST API, and invalid entries were excluded. This process yielded a final set of 25 non-redundant assays. For each assay, we randomly sampled 6–10 active molecules and a complementary number of inactive molecules to form a fixed candidate set of 60 molecules per sample. The order of candidates was fully randomized to mitigate potential ordering bias.

Each test sample was formatted as a JSON object containing the target ChEMBL ID, standardized protein name, and 60 candidate SMILES. Models were instructed to rank all candidates by predicted docking strength (descending order) and return the full ranked list as a JSON array. The ground truth for each sample was defined as all active molecules sorted by experimental pChEMBL values in descending order. Model performance was quantified via Hits@3, the average number of ground-truth active molecules recovered in the top 3 of the model’s ranked list across all samples.

#### MolBench-MO

The MolBench-MO benchmark was curated from the ChemCoTBench dataset [36]: we first randomly sampled a subset of questions from two subtasks (molecular editing and physicochemical property-focused molecular optimization) within the source dataset, with all ambiguous or ill-defined entries systematically removed during preprocessing. The first subtask, functional group modification, requires models to perform targeted addition, deletion or substitution of functional groups on a given source molecule in strict adherence to task-specified constraints, with performance quantified by operational accuracy (the proportion of outputs that fully meet all modification requirements). The second subtask assesses the targeted optimization of drug-relevant physicochemical properties, in which models must optimize input molecules against three key metrics for drug development: QED, LogP and LogS. Performance on this subtask is evaluated via two primary endpoints: the absolute change (delta) in the target property after optimization, and the success rate of achieving pre-specified optimization criteria.

Molecular optimization tasks targeting three specific therapeutic targets were excluded from the final MolBench-MO benchmark. This decision was informed by preliminary validation experiments, which demonstrated that LLMs solved these target-specific tasks predominantly by reproducing established medicinal chemistry strategies memorized from their training data, rather than through de novo rational chemical reasoning. As the primary goal of MolBench-MO is to benchmark true reasoning ability rather than the recall of pre-existing chemical knowledge, inclusion of these tasks would have confounded performance metrics and undermined the validity of cross-model comparative evaluations.

#### MolBench-E2E

While MolBench-MS and MolBench-MO evaluate isolated molecular screening and single-metric optimization capabilities respectively, real-world computational drug discovery campaigns demand the autonomous orchestration of multi-tool, multi-phase workflows in which each step’s output conditions the next step’s input and execution logic. MolBench-E2E was designed to benchmark this long-horizon agentic competence: the ability to plan, execute, self-monitor, and adaptively revise complex scientific workflows that span 8–50+ sequential tool invocations across heterogeneous software ecosystems. The benchmark comprises three end-to-end challenges, each representing a distinct workflow archetype of early-stage structure-based drug discovery.

The first task (E2E-Q1) evaluates the agent’s ability to perform coarse-grained conformational sampling using two force fields with fundamentally different theoretical foundations, GoCa (Gō-model-based C*α* coarse-grained dynamics) and OpenAWSEM (Associative-memory, Water-mediated, Structure and Energy Model), followed by all-atom reconstruction using PULCHRA. Starting from the EGFR kinase domain (PDB: 1M17), the agent must preprocess the crystal structure, prepare force-field-specific inputs for both simulation engines, execute independent coarse-grained molecular dynamics runs, extract representative conformational ensembles, and reconstruct full atomic detail from the reduced representations. This task tests the agent’s capacity to manage format conversions and parameter configurations across fundamentally different simulation paradigms within a single coordinated pipeline.

The second task (E2E-Q2) assesses closed-loop, multi-round molecular property optimization driven by quantitative feedback. E2E-Q2 mandates a structured iterative protocol over up to five rounds: each round requires explicit property assessment, diagnostic reasoning identifying which structural features limit the target metric (QED), generation of candidate modifications with chemical rationale, and quantitative verification before seed selection. The task imposes convergence detection rules (early stopping upon target achievement; convergence declaration after consecutive rounds without improvement) and structural similarity constraints (Tanimoto ≥ 0.40), requiring the agent to balance exploitation of promising modifications against exploration of alternative strategies. The evaluation focus is on the quality of the Assess → Diagnose → Design → Verify reasoning loop across rounds, rather than raw optimization magnitude.

The third task (E2E-Q3), the most complex, requires the agent to integrate receptor preparation, binding site characterization, baseline docking, generative molecular design (via REINVENT 4, supplemented by manual analog design), molecular docking with locked parameters, drug-likeness filtering, and structure–activity relationship (SAR) interpretation within a multi-round optimization loop of up to 15 rounds. The agent must establish and maintain a persistent experimental context, docking box coordinates, baseline scores, a scaffold modification map linking modifiable positions to nearby binding-site residues, across all rounds. Termination is governed by dual criteria: success (≥ 2 molecules achieving a docking score improvement of ≥ 2 kcal/mol relative to Erlotinib) or failure (round 15 reached without meeting the target). A mandatory strategy pivot rule requires the agent to shift to a different modifiable position or multi-site modification if the best score stagnates for three consecutive rounds. This task tests the full integration of tool orchestration, scientific reasoning, and autonomous decision-making that characterizes real medicinal chemistry campaigns.

All three tasks share a common evaluation protocol. Each provides the agent with a target protein specification, a starting molecule or structure, explicit success/failure criteria with quantitative thresholds, and a required output format including structured log files and a final summary report. Agent performance is assessed via task-specific rubrics that decompose each challenge into weighted criteria spanning: (1) correct tool selection and invocation sequence; (2) scientific validity of intermediate reasoning; (3) data integrity and parameter consistency across workflow steps; (4) appropriate termination behavior; and (5) completeness of output deliverables. Full task specifications, per-phase workflow details, and evaluation rubrics are provided in Supplementary Tables S5–S7.

## Data Availability

Both the MolBench dataset (CSV format) and associated evaluation code can be accessed from Github (https://github.com/InternScience/MolClaw).

## Code Availability

The skills adopted in MolClaw, together with the corresponding configuration instructions, are also available at GitHub (https://github.com/InternScience/MolClaw).

## Appendix

### Methods

### Supplementary Note S1: Skill Inventory

**Table 3:**
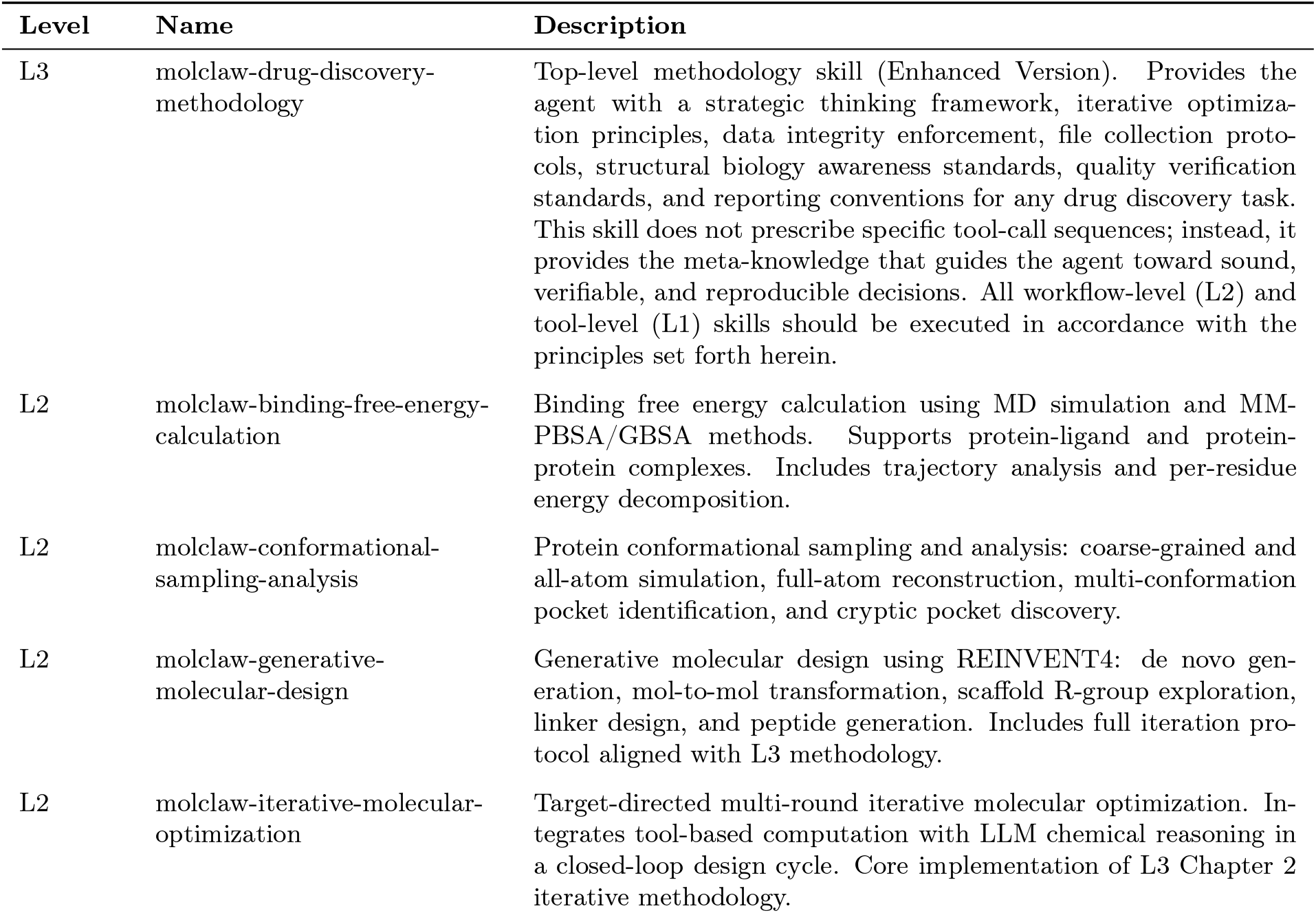
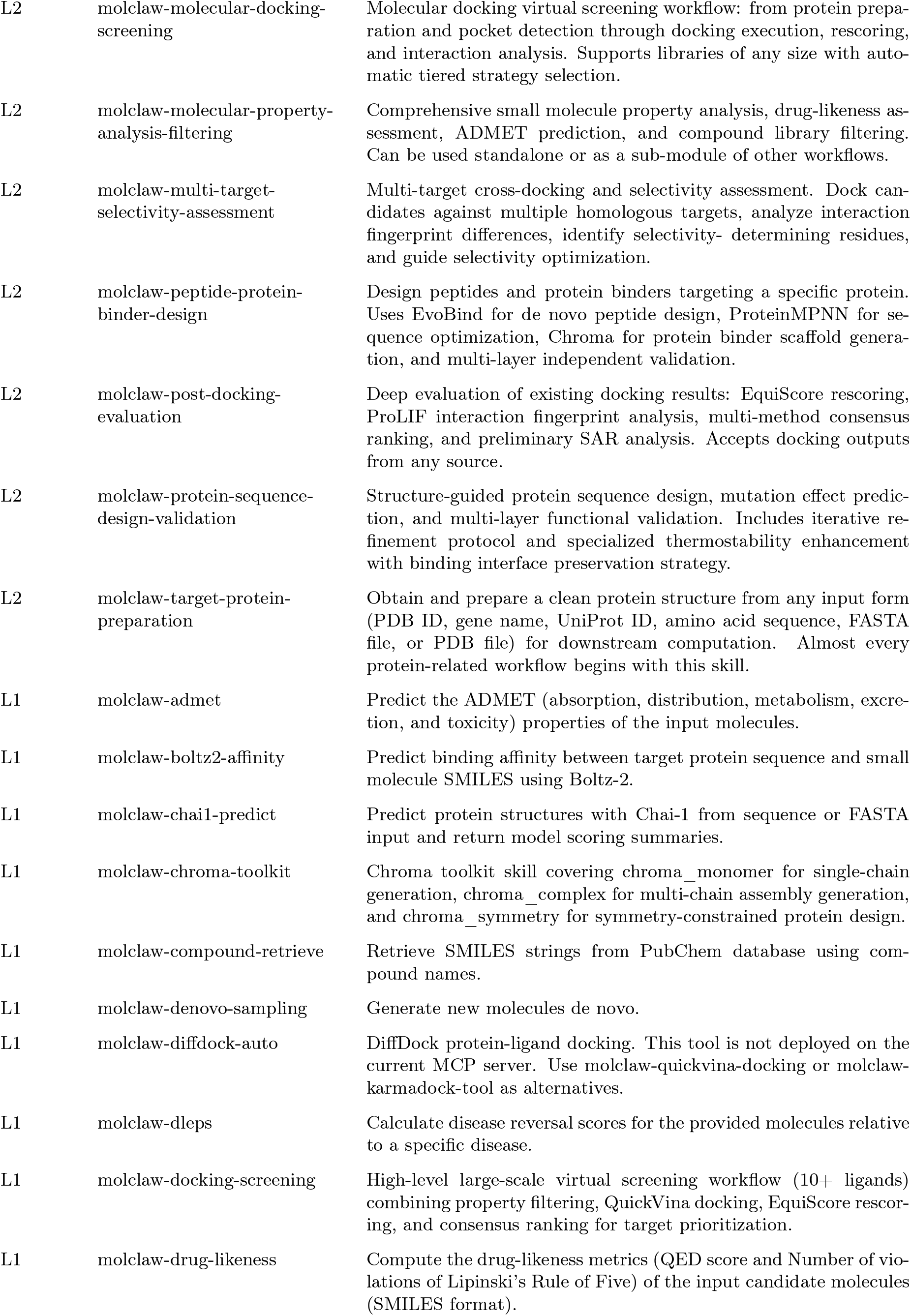
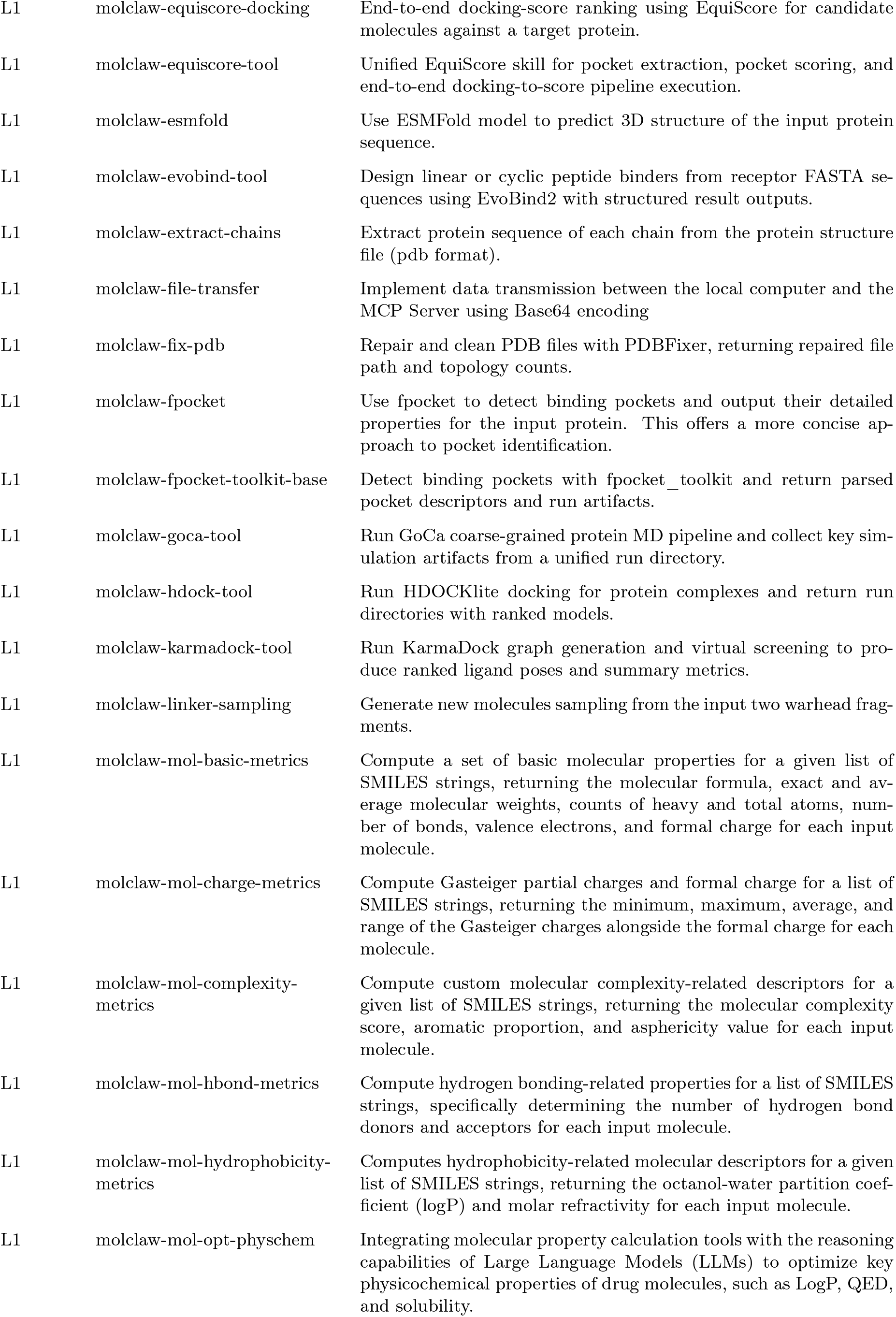
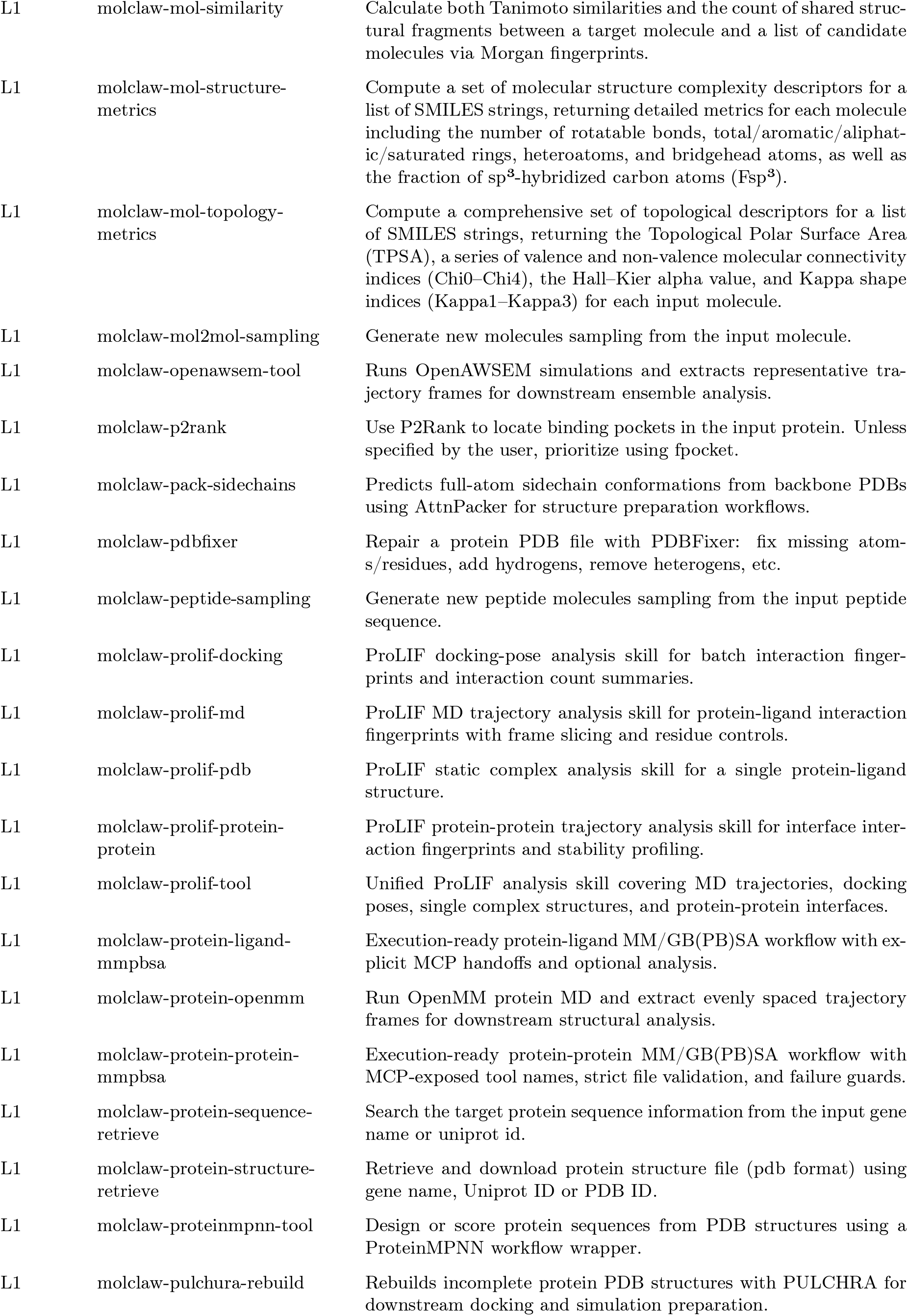
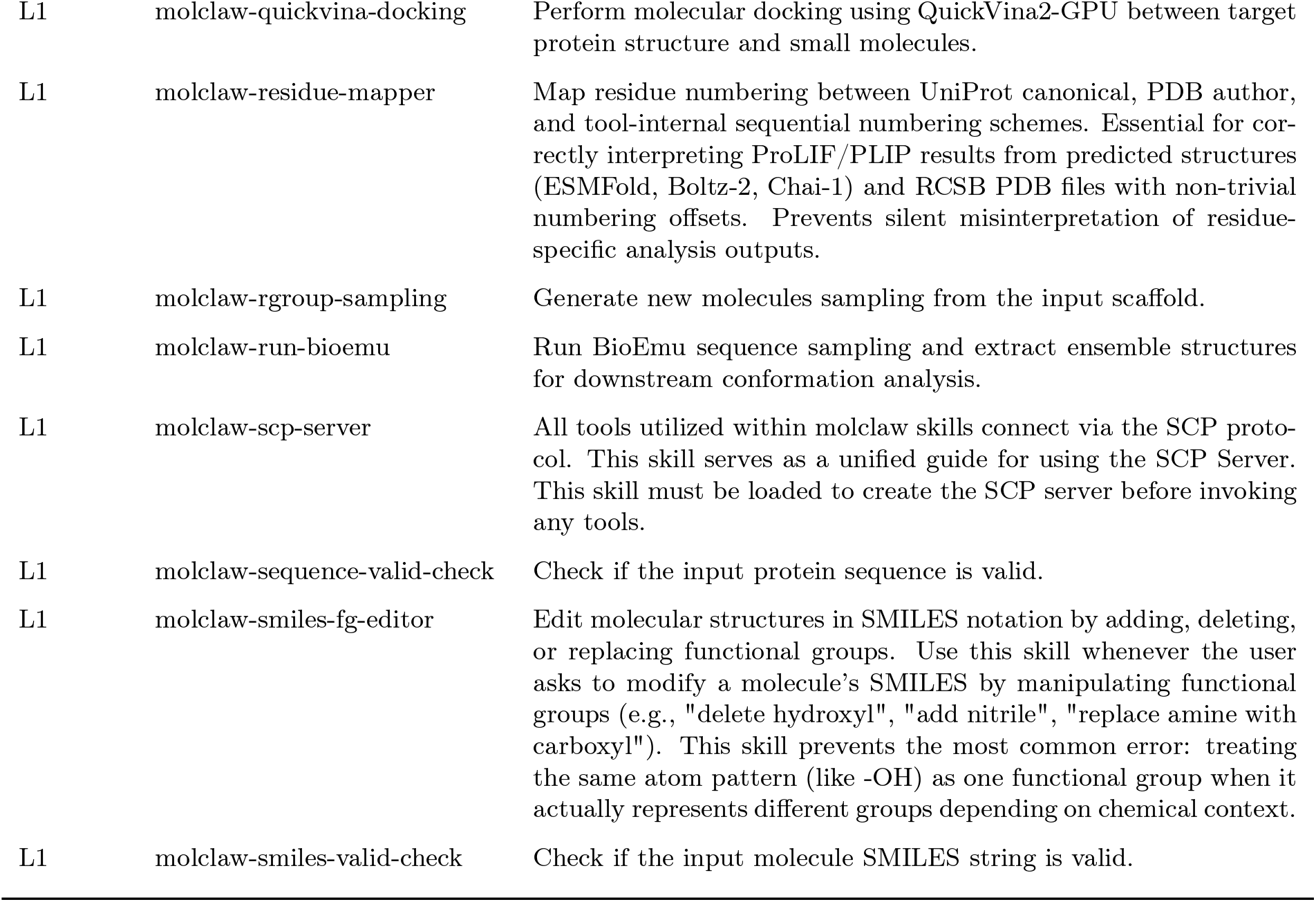
Complete hierarchical skill inventory of the MolClaw agent, including skill level, name, and description.

### Supplementary Note S2: Three-Tier Architecture: Design Rationale

#### S2.1 Problem Statement

An LLM-based drug discovery agent must orchestrate dozens of heterogeneous computational tools— from molecular docking engines and molecular dynamics simulators to deep-learning scoring functions and generative molecular models—within scientifically rigorous workflows. Three categories of failure commonly arise in such systems:

1. **Hallucination and data fabrication**. The LLM may invent plausible but fictitious numerical results, substitute training-knowledge values for tool-computed outputs, or miscount generated molecules.
2. **Domain-logic errors**. Silent failures occur when results from different tools use incompatible conventions (e.g., residue numbering schemes), when docking parameters are set outside physically valid ranges, or when the agent applies a single-point evaluation where an iterative optimization is required.
3. **Incomplete provenance**. Structure files, visualization images, and intermediate results generated on the remote SCP server may not be downloaded to the local workspace, severing the audit trail between reported conclusions and verifiable data.

#### S2.2 Architectural Response

The three-layer architecture addresses these failure modes through separation of concerns:

The layers are connected by cross-references: each L2 workflow declares in its YAML metadata which L3 principles it instantiates, and each L2 workflow specifies which L1 tools it consumes. This ensures consistency: a change to an L3 principle propagates to all L2 workflows that reference it.

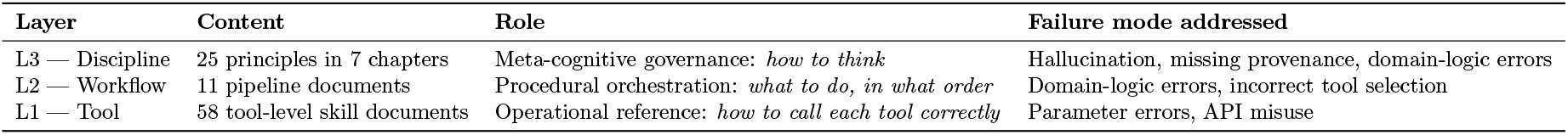

| Layer | Content | Role | Failure mode addressed |
| --- | --- | --- | --- |
| L3 — Discipline | 25 principles in 7 chapters | Meta-cognitive governance: *how to think* | Hallucination, missing provenance, domain-logic errors |
| L2 — Workflow | 11 pipeline documents | Procedural orchestration: *what to do, in what order* | Domain-logic errors, incorrect tool selection |
| L1 — Tool | 58 tool-level skill documents | Operational reference: *how to call each tool correctly* | Parameter errors, API misuse |

#### S2.3 Separation of Naming Conventions

To prevent a frequent runtime failure—the agent passing a *skill name* (kebab-case, e.g., `molclaw-qui ckvina-docking`) where an *SCP tool name* (snake_case, e.g., `molecule_docking_quickvina_ful lprocess`) is expected—the architecture enforces a strict naming convention distinction (L3 Principle 24). Only SCP tool names may appear in `call_tool()` invocations. Every L1 skill document contains the exact `call_tool(“actual_tool_name”,…)` code block to eliminate ambiguity.

#### S2.4 Data-Handoff Contracts

Each L2 workflow terminates with a standardized **Output Specification (Data Handoff Contract)** table that declares, for each output artifact: the variable name, file format, which downstream skills consume it, and the download policy (Category A: must download; Category B: should download; Category C: may skip). This design imports the API-contract paradigm from software engineering into scientific workflow orchestration, enabling the 11 workflows to compose freely without data-transfer failures at workflow boundaries.

### Supplementary Note S3: L3 Discipline Layer: The 25 Principles

The L3 discipline document contains 25 principles organized into seven chapters. Below we describe each chapter’s rationale and key mechanisms in detail.

#### S3.1 Task Understanding and Planning (Principles 1–3)

**Principle 1 (Understand Before Acting)** requires the agent to complete a six-step analysis before invoking the first tool: (1) identify the task type (virtual screening, molecular design, evaluation, structural analysis, or protein/peptide design); (2) distinguish hard constraints from soft constraints; (3) plan the execution path with stage dependencies, parallelization opportunities, and computational bottlenecks; (4) anticipate file collection needs at each step; (5) anticipate residue numbering issues and plan mapping steps; (6) flag computationally required deliverables that must not be replaced by literature values or LLM inference.

Steps (4)–(6) are novel additions that directly target the three failure modes identified in S2.1: step (4) prevents incomplete provenance, step (5) prevents numbering mismatches, and step (6) prevents hallucinated results.

**Principle 2 (Tiered Screening with Progressive Refinement)** encodes the funnel logic of drug discovery as an explicit four-tier strategy:

| Tier | Time per molecule | Method | Typical elimination rate |
| --- | --- | --- | --- |
| 1 | Seconds | Physicochemical filtering (RDKit, drug-likeness) | 20–40% |
| 2 | Seconds–minutes | Molecular docking (QuickVina2-GPU, DiffDock) | Ranking-based selection |
| 3 | Minutes | Multi-dimensional rescoring (EquiScore, ProLIF, Boltz-2) | Cross-validation filtering |
| 4 | Hours | Dynamics-based validation (MD + MM-PBSA) | Final $\leq 5$ candidates |

Critically, Principle 2 mandates that **at every tier boundary, the agent must programmatically count how many molecules survived** and record the screening funnel as verified statistics— not estimates.

**Principle 3 (More Than One Method)** requires cross-validation for all critical computational steps: dual-tool pocket detection (fpocket + P2Rank with consensus logic), multi-method docking scoring (QuickVina + EquiScore or Boltz-2), and at least two independent affinity prediction methods. When methods disagree, the disagreement must be flagged rather than silently resolved—disagreement itself is treated as valuable scientific information.

#### S3.2 Iterative Optimization Methodology (Principles 4–8)

This chapter formalizes the iterative design process that is central to molecular optimization, generative design, and protein engineering tasks.

**Principle 4 (All Design Tasks Should Be Iterative)** declares that molecular generation/optimization, peptide/protein sequence design, and candidate selection tasks must never be completed in a single round. The core logic is a closed loop of evaluate–diagnose–correct–re-evaluate, where each round’s output becomes the next round’s input and each round’s strategy is adjusted based on quantitative analysis of the previous round.

**Principle 5 (Three Questions Per Round)** is the operational gate for each iteration. Before starting any optimization round, the agent must explicitly answer:

1. *What is this round trying to improve?* — Must cite specific data from the previous round (e.g., “4 of the Top 5 molecules had CYP3A4 inhibition probability *>* 0.7”). Generic statements like “continue optimizing” are rejected.
2. *What strategy will be used?* — Must provide a chemical or structural rationale (e.g., “replace methoxyethoxy with cyclopropyloxy to block O-demethylation”).
3. *How will improvement be measured?* — Must define a quantifiable success criterion (e.g., “CYP3A4 probability *<* 0.5 with docking score change *<* 1.5 kcal/mol”).

This design prevents the common failure mode of “blind iteration,” where an agent repeats the same operation without learning from previous results.

**Principle 6 (Exploration–Exploitation Strategy)** defines a rhythm across rounds:

| Phase | Rounds | Similarity constraint | Generation volume | Goal |
| --- | --- | --- | --- | --- |
| Exploration | 1–2 | Low (0.4–0.5) | 30–50 molecules | Discover promising chemical space regions |
| Targeted optimization | 3–4 | Medium (0.6–0.7) | 10–20 molecules | Focus on specific modification sites |
| Convergence | 5+ | High ($\geq 0.8$) | 5–10 molecules | Fine-tune peripheral groups |

This encodes the exploration–exploitation trade-off from reinforcement learning as explicit, schedule-based rules.

**Principle 7 (Convergence Criteria)** defines four stopping conditions: target-met (all objectives achieved), convergence (*<* 5% change in key metrics for two consecutive rounds), trade-off (Pareto frontier reached), and resource limit. The report must state which condition triggered termination.

**Principle 8 (Complete Iteration Records)** requires saving per-round records including all generated SMILES with evaluation results, the strategy and its rationale, delta metrics versus the previous round, and the cumulative global best molecule. The final report must include an optimization trajectory table with verified values.

#### S3.3 Data Integrity, Verification, and Quality Control (Principles 9–13)

This chapter constitutes the most critical guardrail against result fabrication and is the centerpiece of our anti-hallucination design.

**Principle 9 (Never Trust a Single Tool’s Output at Face Value)** establishes immediate plausibility checks: deterministic computations (MW, LogP) must be internally consistent; docking scores must be negative (positive values indicate failure); predictions must be compared against known experimental values when available; and different physical quantities (docking scores, Δ*G, K*_*d*_, IC_50_) must never be conflated.

**Principle 10 (Three-Category Information Distinction)** requires all reported values to be annotated as one of three categories:

- **Category 1 — Tool-computed facts:** Precise numerical values from tool calls, citing the source tool (e.g., “QuickVina docking score: −8.3 kcal/mol”).
- **Category 2 — Agent interpretations:** Inferences drawn by the agent from computed data, explicitly labeled (e.g., “This score suggests moderate binding potential (agent analysis)”).
- **Category 3 — Literature-derived values:** Data from published sources, marked with a warning label: “LITERATURE VALUE — not computed in this session,” with full citation.

This three-category system ensures that a downstream researcher can immediately identify which values are independently reproducible from tool outputs, which are analytical judgments, and which require literature verification.

**Principle 11 (Count-Before-Report)** is the primary anti-fabrication mechanism. Before writing ANY numerical claim—in run logs, round summaries, or the final report—the agent must execute an explicit programmatic verification against the actual source file. The principle specifies concrete verification methods for each claim type:

| Claim type | Verification method | Example |
| --- | --- | --- |
| “Generated $N$ molecules” | Count entries in output file | `python3 -c "import json; print(len(json.load(open('file.json'))))"` |
| “$M$ molecules passed filter” | Count entries in filtered output | `grep -c "PASS" results.csv` |
| “Docking score of $X$” | Re-read exact tool return | Direct citation of tool return value |

The verification command and its result must be recorded in the execution log. If the verified count differs from expectations, the *actual* count must be reported—not the expected count. This principle is declared non-negotiable and applies to every number in every report.

**Principle 12 (Three-Checkpoint Self-Audit)** distributes verification across three mandatory checkpoints:

- **Checkpoint A (after each tool call):** Immediate sanity checks—docking score sign, ADMET probability range [0, 1], MW–formula consistency, output file existence and non-zero size, structure prediction confidence thresholds.
- **Checkpoint B (before each round summary):** Re-read ALL tool output files from the round. Verify molecule counts, scores, and selected candidates against actual file contents.
- **Checkpoint C (before the final report):** Create a Data Integrity Verification table mapping every key reported value to its source file and verification command:

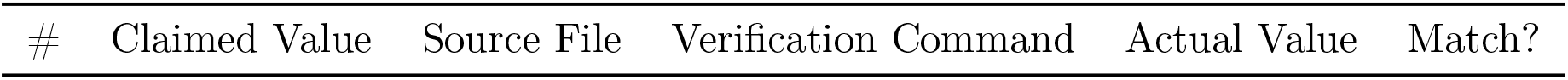

Any mismatch must be resolved by correcting the report, not the data.

**Principle 13 (Computation-First Hierarchy)** establishes a strict four-level fallback:

Any mismatch must be resolved by correcting the report, not the data. **Principle 13 (Computation-First Hierarchy)** establishes a strict four-level fallback:
| Level | Authority | Source | Action |
| --- | --- | --- | --- |
| 1 (highest) | Direct tool computation | Primary tool computes the value | Always prefer |
| 2 | Alternative tool computation | Backup tool if primary fails | Try before literature |
| 3 | Approximate computation | Indirect proxy (e.g., LogP for solubility trends) | Try before literature |
| 4 (lowest) | Literature reference | Published values | Last resort only, with full protocol |

Level 4 (literature) requires: (a) pause and re-examine whether any tool can compute even an approximation; (b) label with “LITERATURE VALUE”; (c) cite with authors, year, DOI; (d) cross-check against a second independent source; (e) assess context match; (f) explain why computation was impossible.

The following are strictly forbidden: presenting literature values as computational results; generating complex analyses (interaction maps, selectivity profiles) from LLM training knowledge; using LLM chemical intuition to fill in numbers that a tool should have produced.

#### S3.4 File Collection and Data Provenance (Principles 14–17)

This chapter addresses the systematic problem of incomplete file collection from the remote SCP server.

**Principle 14 (Mandatory Collection of ALL Structure Files)** requires downloading every molecular structure file generated during execution—PDB, PDBQT, SDF, MOL2, CIF, GRO, XTC, and 10+ other formats—using a standard download-verify protocol: `server_file_to_base64` → local decode → verify `os.path.getsize()>0`. The principle includes a comprehensive table mapping every tool category (docking, affinity prediction, structure prediction, MD simulation, etc.) to its expected structure output fields. The guiding principle is stated as “over-download rather than under-download.”

**Principle 15 (Mandatory Collection of ALL Image Files)** extends the same protocol to visualization outputs (PNG, SVG, TIFF, etc.), with a table mapping analysis types to expected image outputs.

**Principle 16 (User-Critical File Identification)** classifies files into three categories:

| Category | Policy | Examples |
| --- | --- | --- |
| A — User-critical | MUST download | Structure files, result tables, visualization images, molecule lists |
| B — Diagnostic | SHOULD download | Tool log files, configuration files, parameter records |
| C — Ephemeral | MAY skip | Temporary intermediates, cache files |

The default policy is to download all Category A and B files.

**Principle 17 (Universal File Download Rule)** states the overarching rule: every SCP tool call that returns a file path must be followed by downloading that file before proceeding to the next step. A tool call is not considered complete until its output files are downloaded and verified locally.

#### S3.5 Structural Biology Awareness (Principles 18–19)

**Principle 18 (Residue Numbering Reconciliation)** addresses the single most common source of silent errors in automated computational structural biology pipelines. The principle identifies four numbering schemes that coexist in typical tasks:

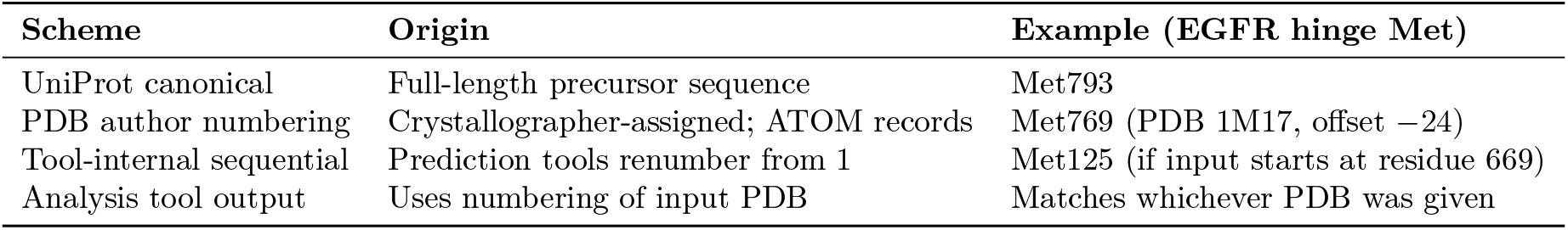

| Scheme | Origin | Example (EGFR hinge Met) |
| --- | --- | --- |
| UniProt canonical | Full-length precursor sequence | Met793 |
| PDB author numbering | Crystallographer-assigned; ATOM records | Met769 (PDB 1M17, offset −24) |
| Tool-internal sequential | Prediction tools renumber from 1 | Met125 (if input starts at residue 669) |
| Analysis tool output | Uses numbering of input PDB | Matches whichever PDB was given |

The mandatory mapping protocol consists of five steps: (a) flag mapping needs at planning time; (b) build a complete mapping table before the first residue-specific analysis, using DBREF records, arithmetic offset, or sequence alignment; (c) record the mapping table in the execution log; (d) translate all tool outputs to the task’s reference scheme before drawing conclusions; (e) include the mapping table in the final report.

To implement this protocol, we developed a dedicated SCP tool, `residue_mapper`, that automates all three mapping strategies (arithmetic for predicted structures, DBREF-based for RCSB PDB files, Needleman–Wunsch alignment as fallback), supports forward and reverse lookups with a mixed query syntax (`Met793`, `pdb:769`, `tool:76` can coexist in one query), and outputs a standardized CSV mapping table. The tool is described in detail in the L1 skill `molclaw-residue-mapper`.

The principle further specifies common traps: searching for UniProt residue numbers in prediction-tool outputs (they will not be found); reporting ProLIF residue IDs without noting they are in tool-internal numbering; assuming two PDB files for the same protein use the same numbering.

**Principle 19 (Docking Parameter Safeguards)** imposes hard constraints on docking box parameters: a minimum of 25.0 Å per dimension (non-negotiable floor), and a progressive box enlargement protocol on failure (25 → 30 → 40 → 50 Å → switch method). Additional checks include: pocket center must not be (0, 0, 0) or default values; identical scores across all molecules signal systematic setup errors; batch-wide docking failure suggests receptor preparation issues rather than ligand issues.

#### S3.6 Reporting Standards (Principles 20–23)

**Principle 20** defines a standardized report structure including: task summary, methods overview, residue numbering reference, results (with source-tool annotations and three-category distinctions), iteration history, final recommendations, downloaded files summary, limitations discussion, and suggested experimental next steps.

**Principle 21 (Honest Annotation of Uncertainty)** enumerates ten specific situations that must be flagged: predicted vs. experimental structure used; divergent pocket detection results; inconsistent docking/rescoring rankings; ADMET confidence limitations; lack of experimental validation; literature values used as fallback; residue numbering mapping applied; generation count discrepancies; docking box enlargement retries; any failed computational steps.

**Principles 22–23** mandate a data provenance audit table (mapping every key value to its source file and verification command) and a complete file inventory at execution end.

#### S3.7 System Architecture Awareness (Principles 24–25)

**Principle 24** enforces the distinction between SCP tool names (snake_case, used in `call_tool()`) and skill names (kebab-case, used as documentation identifiers), preventing the most common runtime error in the three-layer architecture. It also maintains a list of currently unavailable tools to prevent calls to undeployed endpoints.

**Principle 25 (SCP File-Transfer Enforcement)** operationalizes the file collection mandates of Principles 14–17 at the system architecture level, requiring that every SCP tool call returning a file path be followed by a download-and-verify step before the agent proceeds. While Principle 17 states this as a universal methodological rule, Principle 25 encodes it as a system-level constraint tied to the SCP server’s file-transfer mechanism, ensuring enforcement even when the agent’s methodological context is not loaded.

### Supplementary Note S4: L2 Workflow Layer: Pipeline Design

The 11 L2 workflows collectively cover the major computational pipelines in small-molecule and biologic drug discovery. Each workflow follows a standardized structure consisting of: applicability conditions (when to use / when not to use), prerequisites (input dependencies from other workflows), a phase-by-phase execution protocol with embedded quality-gate checkpoints, common failure-and-recovery tables, and an output specification (data handoff contract).

#### S4.1 Workflow Coverage

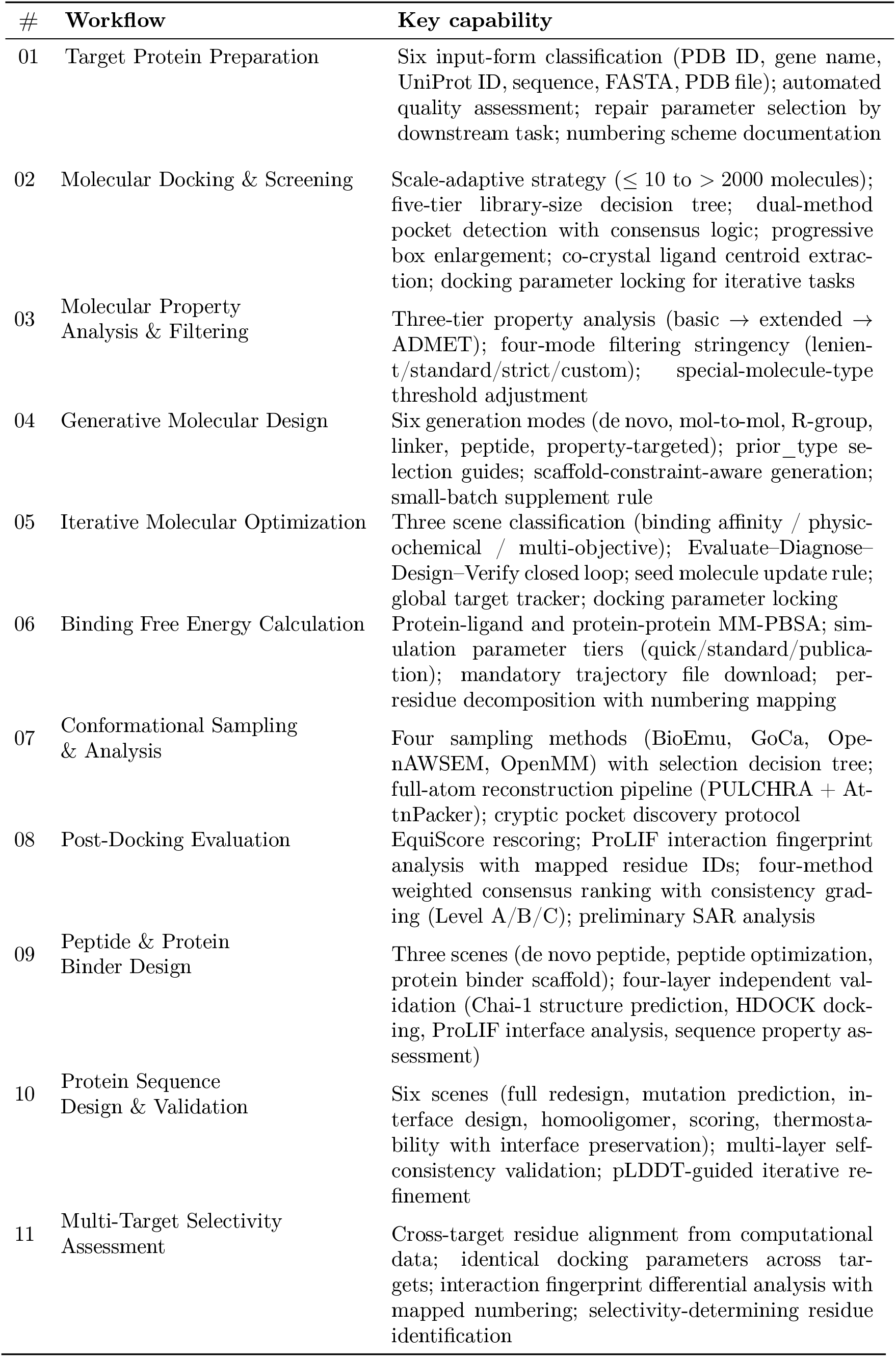

| # | Workflow | Key capability |
| --- | --- | --- |
| 01 | Target Protein Preparation | Six input-form classification (PDB ID, gene name, UniProt ID, sequence, FASTA, PDB file); automated quality assessment; repair parameter selection by downstream task; numbering scheme documentation |
| 02 | Molecular Docking & Screening | Scale-adaptive strategy ($\leq 10$ to $> 2000$ molecules); five-tier library-size decision tree; dual-method pocket detection with consensus logic; progressive box enlargement; co-crystal ligand centroid extraction; docking parameter locking for iterative tasks |
| 03 | Molecular Property Analysis & Filtering | Three-tier property analysis (basic → extended → ADMET); four-mode filtering stringency (lenient/standard/strict/custom); special-molecule-type threshold adjustment |
| 04 | Generative Molecular Design | Six generation modes (de novo, mol-to-mol, R-group, linker, peptide, property-targeted); prior_type selection guides; scaffold-constraint-aware generation; small-batch supplement rule |
| 05 | Iterative Molecular Optimization | Three scene classification (binding affinity / physicochemical / multi-objective); Evaluate-Diagnose-Design-Verify closed loop; seed molecule update rule; global target tracker; docking parameter locking |
| 06 | Binding Free Energy Calculation | Protein-ligand and protein-protein MM-PBSA; simulation parameter tiers (quick/standard/publication); mandatory trajectory file download; per-residue decomposition with numbering mapping |
| 07 | Conformational Sampling & Analysis | Four sampling methods (BioEmu, GoCa, OpenAWSEM, OpenMM) with selection decision tree; full-atom reconstruction pipeline (PULCHRA + AttnPacker); cryptic pocket discovery protocol |
| 08 | Post-Docking Evaluation | EquiScore rescoring; ProLIF interaction fingerprint analysis with mapped residue IDs; four-method weighted consensus ranking with consistency grading (Level A/B/C); preliminary SAR analysis |
| 09 | Peptide & Protein Binder Design | Three scenes (de novo peptide, peptide optimization, protein binder scaffold); four-layer independent validation (Chai-1 structure prediction, HDOCK docking, ProLIF interface analysis, sequence property assessment) |
| 10 | Protein Sequence Design & Validation | Six scenes (full redesign, mutation prediction, interface design, homooligomer, scoring, thermostability with interface preservation); multi-layer self-consistency validation; pLDDT-guided iterative refinement |
| 11 | Multi-Target Selectivity Assessment | Cross-target residue alignment from computational data; identical docking parameters across targets; interaction fingerprint differential analysis with mapped numbering; selectivity-determining residue identification |

#### S4.2 Key Workflow Design Patterns

Several design patterns recur across the 11 workflows:

##### Decision-tree entry points

Rather than a single linear protocol, each workflow begins with a classification step that routes execution to the appropriate sub-protocol. For example, Workflow 02 selects one of five screening strategies based on library size (≤ 10, 11–100, 101–500, 501–2000, *>* 2000); Workflow 05 classifies optimization tasks into three scenes (binding, physicochemical, multi-objective); Workflow 10 offers six scenes covering different protein design goals.

##### Quality-gate checkpoints embedded in the pipeline

Each workflow contains multiple CHECKPOINT blocks—implemented as checklists—that halt execution until verification criteria are satisfied. These checkpoints are the operational instantiation of L3 Principle 12 (Three-Checkpoint Self-Audit). For example, Workflow 02 includes checkpoints after molecule preparation (counts verified), after pocket detection (box ≥ 25 Å, center not at origin), after each docking call (score negative, pose file downloaded), and after ranking (counts consistent across scoring methods, ProLIF residues mapped).

##### COUNT GATE protocol

At every stage where the number of molecules changes (validation, filtering, generation, docking, selection), a COUNT GATE block requires programmatic verification of the actual count from source files. This is the workflow-level operationalization of L3 Principle 11.

##### MAPPING GATE protocol

Before any residue-specific analysis interpretation, a MAPPING GATE block requires execution of the residue numbering mapping if the analysis structure uses a different numbering scheme from the task description. This is the workflow-level operationalization of L3 Principle 18.

##### Failure-and-recovery tables

Each workflow includes a structured table of common failures, their likely causes, and specific recovery actions. These encode domain-expert troubleshooting knowledge in a machine-readable format. For instance, Workflow 02 specifies: “All docking scores positive → likely wrong pocket or box too small → re-detect pocket; try progressive box enlargement 25 → 30 → 40 → 50 Å.” Workflow 05 specifies: “LLM designs invalid SMILES 3 times → switch to REINVENT mol2mol sampling with high_similarity prior as fallback generator.”

##### Explicit boundary conditions

Each workflow declares not only when to use it, but also when NOT to use it and which workflow to use instead. For example, Workflow 02 states: “Do NOT use when the user already has docking results and only wants post-hoc evaluation—use Workflow 08 instead.” This prevents incorrect workflow selection.

### Supplementary Note S5: L1 Tool Layer: Coverage and Organization

The 58 L1 tool skill documents cover the following functional categories:

| Category | Tool count | Representative tools |
| --- | --- | --- |
| Molecular docking & scoring | 6 | QuickVina2-GPU, KarmaDock, DiffDock, EquiScore (pocket + screen + pipeline), docking-screening |
| Binding affinity prediction | 1 | Boltz-2 protein-ligand affinity |
| Molecular generation (REINVENT4) | 5 | De novo, mol-to-mol, R-group, linker, peptide sampling |
| Molecular property computation | 8 | Basic metrics, hydrophobicity, H-bond, charge, topology, complexity, structure, drug-likeness |
| ADMET prediction | 1 | ADMET-AI (90+ endpoints) |
| Molecular similarity & editing | 3 | Morgan fingerprint similarity, SMILES functional-group editor, SMILES validity check |
| Protein structure retrieval | 3 | By PDB ID, gene name, UniProt ID |
| Protein structure prediction | 3 | ESMFold, Chai-1, Boltz-2 |
| Protein structure processing | 6 | PDB repair (fix_pdb, pdbfixer), chain extraction, PULCHRA rebuild, AttnPacker side-chain packing, compound retrieval |
| Pocket detection | 3 | fpocket, fpocket-toolkit-base, P2Rank |
| Interaction analysis (ProLIF) | 5 | MD trajectory, docking poses, single PDB, protein-protein, unified toolkit |
| MD simulation | 2 | OpenMM protein MD, BioEmu conformational sampling |
| Coarse-grained simulation | 2 | GoCa, OpenAWSEM |
| Free energy calculation (MM-PBSA) | 2 | Protein-ligand MM-PBSA, protein-protein MM-PBSA |
| Protein design | 3 | ProteinMPNN, EvoBind2, Chroma toolkit (monomer + complex + symmetry) |
| Protein-protein docking | 1 | HDOCK |
| Residue numbering | 1 | residue_mapper |
| Molecular optimization (composite) | 2 | Target-directed optimization, physicochemical optimization |
| File management | 2 | File transfer (base64), SCP server connection |
| Sequence validation | 2 | Protein sequence validity, SMILES validity |
| Deep-learning electrostatic potential | 1 | DLEPS |

Each L1 document follows a consistent structure: YAML metadata header (name, description, license, author), prerequisite notes (file upload, PDB preprocessing), SCP server connection boilerplate, tool description block (arguments, return values), usage code examples with exact `call_tool()` invocations, and for composite tools, multi-step workflow guidance with inter-skill references.

For complex tool families (ProLIF, EquiScore, Chroma, MM-PBSA), a unified toolkit skill document covers all related tools in one place, with scenario-based routing guidance. The two MM-PBSA skills additionally include separate reference documents for each sub-step (fix_pdb, prepare_complex, run_mmpbsa, analyze_mmpbsa), providing fine-grained parameter guidance.

### Supplementary Note S6: Key Design Innovations: Detailed Analysis

#### S6.1 Anti-Hallucination as Institutional Design

Existing approaches to reducing LLM hallucination in agent systems primarily rely on model-level interventions (RLHF, prompt engineering) or output-level checks (fact-verification tools). Our skill library takes a fundamentally different approach: it treats anti-hallucination as an **institutional design problem**—building verification requirements into the workflow infrastructure itself so that fabrication becomes procedurally impossible rather than merely discouraged.

The anti-hallucination mechanism consists of four interlocking components:

1. **Programmatic verification gates (Principle 11):** Every numerical claim must be preceded by an explicit verification command against the source file. The command and its result are recorded in the execution log, creating an audit trail that an external reviewer can independently check.
2. **Three-checkpoint continuous audit (Principle 12):** Verification is not deferred to the end; it occurs after every tool call (A), before every round summary (B), and before the final report (C). This distribution prevents error accumulation.
3. **Three-category information labeling (Principle 10):** By requiring explicit annotation of each value’s provenance (tool-computed / agent-inferred / literature-derived), the system eliminates the ambiguity that enables hallucination—the agent cannot present an LLM-generated value without visibly violating the labeling protocol.
4. **Computation-first hierarchy (Principle 13):** By establishing a strict four-level preference order (direct computation → alternative computation → approximate computation → literature), the system structurally prevents the agent from taking the path of least resistance (generating values from training knowledge).

These four components form a **trust chain**: any reported value can be traced backward through the audit table (Principle 12, Checkpoint C) to a verification command (Principle 11) that reads a source file (Principle 14) produced by a tool call (Principle 10, Category 1) that was the agent’s first choice in the computation hierarchy (Principle 13). Breaking any link in this chain produces a visible, auditable violation.

#### S6.2 Residue Numbering: An Engineering Solution to a Pervasive Scientific Problem

Residue numbering mismatches are a well-known problem in computational structural biology, but they are rarely addressed systematically in automated pipelines. In our system, the problem is particularly acute because a single task may involve:

- An RCSB PDB file with author numbering (e.g., Met769 in PDB 1M17)
- A Boltz-2 prediction with sequential numbering (e.g., Met76)
- ProLIF interaction analysis outputting the Boltz-2 numbering
- A task description referencing UniProt numbering (e.g., Met793)

Without explicit mapping, the agent would conclude “Met793 interaction absent” when it is actually present as Met76—a false negative with downstream consequences for molecular optimization decisions.

Our solution operates at three levels:

1. **Discipline level (L3 Principle 18):** Establishes the mandatory mapping protocol as a universal principle, with five concrete steps and detailed examples of correct vs. incorrect interpretation.
2. **Tool level (L1** `residue_mapper`): A dedicated SCP tool implementing three mapping strategies in priority order: arithmetic offset (instant, exact) for predicted structures; DBREF-based offset (instant, exact) for RCSB PDB files; Needleman–Wunsch alignment (seconds, robust) as universal fallback. The tool supports seven usage scenarios, mixed query syntax, reverse lookups, multi-chain complexes, and offline mode.
3. **Workflow level (L2 MAPPING GATEs):** At every point in every workflow where residue-specific results are interpreted—ProLIF analysis in Workflow 02/08, per-residue energy decomposition in Workflow 06, interface contacts in Workflow 09/10, cross-target comparison in Workflow 11—a MAPPING GATE block enforces execution of the mapping protocol before conclusions are drawn.

This three-level design ensures that even if the agent “forgets” the L3 principle during a long execution, the L2 workflow’s MAPPING GATE will catch the omission at the point of use.

#### S6.3 Structured Iteration as Scientific Method

Many agent systems implement iteration as a simple retry loop (“if the result is unsatisfactory, try again”). Our skill library transforms iteration from ad hoc repetition into a formalized scientific process through four mechanisms:

1. **Mandatory diagnostic gates (Principle 5):** Each round must begin by answering three questions grounded in data, preventing blind repetition.
2. **Scheduled strategy evolution (Principle 6):** The exploration–exploitation rhythm prevents the agent from either exploring randomly throughout (never converging) or converging prematurely on a suboptimal solution.
3. **Formal convergence criteria (Principle 7):** Four explicit stopping conditions prevent infinite loops while ensuring the agent does not stop prematurely.
4. **State management mechanisms:** The seed molecule update rule (Workflow 05) ensures the correct starting point for each round. The global target tracker (Workflow 05) maintains cumulative progress across rounds. The docking parameter locking protocol (Workflows 02/05) ensures comparability across rounds by preventing inadvertent parameter drift.

Together, these mechanisms transform iteration from a brute-force loop into an auditable Evaluate– Diagnose–Design–Verify cycle where each round’s strategy is a response to the previous round’s quantitative analysis—closely mirroring the design-make-test-analyze (DMTA) cycle in experimental drug discovery.

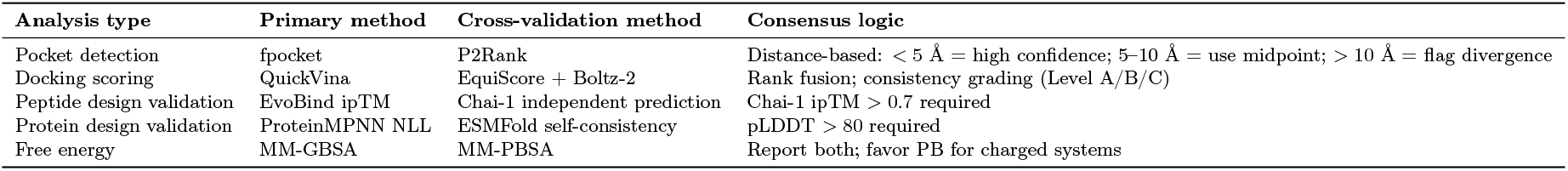

#### S6.4 Multi-Method Cross-Validation as Default

Rather than treating cross-validation as optional, the skill library makes it the default operating mode:

This systematic cross-validation means that false positives from any single method are caught before they propagate downstream.

#### S6.5 Domain Knowledge Operationalization

The skill library encodes numerous pieces of domain-expert tacit knowledge as explicit, executable rules:

- DiffDock’s confidence score is valid only for comparing poses of the **same** molecule; it must never be used to rank different molecules against each other (Workflow 02).
- Docking scores are ranking tools, not thermodynamically exact free energies; converting ΔScore to selectivity fold-change via Δ*G* = *RT* · ln(*K*_*d*_) is explicitly forbidden (Principle 9, Workflow 11).
- EvoBind’s ipTM is a design score, not an independent assessment; third-party validation (Chai-1) is mandatory (Workflow 09).
- ESMFold quality degrades beyond 800 residues; longer sequences should use Chai-1 (Workflow 01).
- When co-crystal ligand centroid extraction is needed, it must occur *before* `fix_pdb` removes heteroatoms—order of operations is critical (Workflow 02).
- Peptide drugs have stability, permeability, and immunogenicity concerns not covered by standard ADMET tools; these must be discussed qualitatively based on sequence features (Workflow 09).

Each of these represents a failure mode that an LLM without domain-specific guidance would likely encounter, potentially producing scientifically invalid results.

### Results and Discussion

### Supplementary Information: Statistical Methodology and Detailed Analyses

#### Supplementary Note 1. Statistical methodology

All statistical analyses were performed in Python 3.11 using SciPy (v1.11), statsmodels (v0.14), and scikit-posthocs (v0.9). This note describes the rationale and procedures for the tests referenced in the main text and throughout this Supplementary Information.

##### Proportion-based comparisons

For accuracy-based metrics (property filtering accuracy, binding affinity accuracy, molecule editing accuracy, and optimization success rate), each method’s performance on a given task was treated as a binomial outcome (correct or incorrect), yielding a proportion k/n where k is the number of correct responses, and n is the total number of test items. Pairwise comparisons between MolClaw-CC and each baseline were conducted using Fisher’s exact test (two-sided), which is appropriate for small-to-moderate sample sizes (*n* = 37–50 in this study) where the chi-squared approximation may be unreliable. Effect sizes were quantified using Cohen’s h, defined as 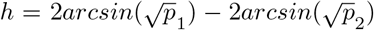, where *p*_1_ and *p*_2_ are the two proportions being compared. By convention, | *h* | *<* 0.2 is considered negligible, 0.2–0.5 small, 0.5–0.8 medium, and > 0.8 large. To control the family-wise error rate across multiple comparisons, we applied both Bonferroni correction and Benjamini–Hochberg false discovery rate (FDR) correction; conclusions reported in the main text remain significant under FDR correction in all cases where significance is claimed.

##### Confidence intervals

Wilson score 95% confidence intervals were computed for all proportion-based metrics. The Wilson interval is preferred over the Wald (normal approximation) interval for proportions near 0 or 1, which is the case for several methods on property filtering (for example, GPT at 8.0%) and molecule editing (for example, MolClaw-CC at 100.0%). Confidence intervals are reported in Fig. 4B–E and referenced in the main text.

##### Multi-method ranking

To assess whether the overall performance ranking across methods was significantly non-uniform, we applied the Friedman test, a non-parametric alternative to repeated-measures ANOVA suitable for ranked data. The test was conducted over four accuracy-based tasks (property filtering, binding affinity, molecule editing, and optimization success rate) as blocks, with 12 methods as treatments (Biomni was excluded due to missing values on three of four tasks). Post-hoc pairwise comparisons were performed using the Nemenyi test.

##### Category-level comparisons

To compare the aggregate performance of standalone LLMs (*n* = 8 methods) against agent-based systems (*n* = 4 methods, excluding Biomni), we applied one-sided Mann–Whitney U tests (agents > LLMs). The results are presented in Fig. 4F–I.

##### Non-proportion metrics

Docking hit count and optimization delta (Δ) are continuous-valued and were compared descriptively, as the single aggregate value per method precludes conventional parametric or non-parametric testing at the method level. We report absolute and relative differences for these metrics.

#### Supplementary Note 2. The relationship between task complexity and skill-driven performance gain

A central finding of this study is that MolClaw’s hierarchical skills do not provide uniform improvement across all tasks (Fig. 5A–C). Rather, the magnitude of skill-driven performance gain varies systematically with the degree to which a task depends on structured, domain-specific computational workflows. This pattern, observed consistently across both the Claude Code and OpenClaw platforms, provides direct evidence for the functional necessity of the three-level skill hierarchy (L1–L2–L3) proposed in MolClaw’s architecture.

To formalize this observation, we computed the average absolute skill gain for each metric as the mean of the MolClaw-versus-vanilla differences on the two platforms:

Property filtering accuracy: average gain = 0.0 percentage points (CC: +0.0, OC: +0.0). This task is fully solvable by ad hoc RDKit scripting; hierarchical skills add no measurable value.

Molecule editing accuracy: average gain = +1.3 percentage points (CC: +2.6, OC: +0.0). Both vanilla agents already achieve ≥ 97.4% accuracy; the remaining gains are marginal and statistically non-significant.

Optimization success rate: average gain = +3.9 percentage points (CC: +7.7, OC: +0.0). On the OpenClaw platform, both vanilla and MolClaw achieve 100.0%, creating a ceiling that masks potential differences.

Binding affinity comparison accuracy: average gain = +25.7 percentage points (CC: +29.7, OC: +21.6). This is the largest absolute gain on any metric and the only one reaching statistical significance on the Claude Code platform (Fisher’s exact test, P = 0.013, Cohen’s h = 0.64; Fig. 5A).

Docking screening hit count: average gain = +0.34 (CC: +0.24, OC: +0.44). Relative to the vanilla baselines, this represents a 43% improvement on the Claude Code platform and a 220% improvement on the OpenClaw platform.

Optimization delta (Δ): average gain = +0.50 (CC: +0.858, OC: +0.143). On the Claude Code platform, MolClaw-CC achieves nearly double the optimization magnitude of its vanilla counterpart (1.724 versus 0.866, a ratio of 2.0 ×; Fig. 5C).

This ordering is not coincidental. Property filtering requires only the correct invocation of a cheminformatics library—a single L1-level operation that any code-generating agent can reproduce. Molecule editing similarly relies on well-defined SMILES transformations that LLMs can express directly. In contrast, binding affinity comparison requires orchestrating a multi-step computational pipeline (molecular format conversion, receptor preparation, docking box definition, docking execution, and score interpretation)—precisely the type of validated workflow encoded in MolClaw’s L2 skills. Physicochemical property optimization goes further, demanding an iterative loop of structural design, computational verification, and strategy adaptation that corresponds to MolClaw’s L3 pipeline-level skill. The progressive increase in skill-driven gain from L1-amenable tasks to L2-dependent tasks to L3-dependent tasks thus provides empirical validation of the three-level hierarchy itself, demonstrating that each level addresses a genuine capability gap rather than serving as mere organizational abstraction.

#### Supplementary Note 3. Binding affinity comparison as the primary discriminative task in MolBench

Among all MolBench sub-tasks, binding affinity comparison exhibits the widest performance spread between method categories and provides the clearest separation between tool-augmented and skill-augmented agents. This section examines the task in detail.

The task presents matched molecular pairs with experimentally measured Ki values and asks models to identify which molecule binds more strongly to a given target (*n* = 37 pairs). As a binary classification task, the chance-level accuracy is 50%. Notably, four of eight standalone LLMs performed at or below this chance level: GPT 5.2 at 48.7% (18/37), DeepSeek v3.2 at 43.2% (16/37), GLM 5 at 24.3% (9/37), and Minimax 2.5 at 16.2% (6/37) (Table 1; Fig. 3A). The remaining four LLMs exceeded 50% only modestly: Gemini 3 and Claude Sonnet 4.6 at 56.8% (21/37), Kimi 2.5 and Qwen 3.5 at 51.4% (19/37). The overall LLM mean of 43.6% does not significantly differ from chance, indicating that current frontier LLMs possess no reliable internal representation of structure–activity relationships sufficient for pairwise binding affinity judgment.

A critical observation is that vanilla agent frameworks (Claude Code and OpenClaw, both at 51.4%) failed to improve upon the best standalone LLMs on this task—in contrast to property filtering, where vanilla agents dramatically outperformed standalone LLMs (98.0% versus 26.0% mean; Fig. 3E versus Fig. 3A). This discrepancy is informative: for property filtering, the performance bottleneck is tool access (the ability to invoke RDKit), which vanilla agents solve through ad hoc code generation. For binding affinity comparison, the bottleneck is not tool access per se but the correct orchestration of a multi-step computational pipeline—preparing molecular structures, configuring docking parameters, executing the docking calculation, and interpreting the results within a validated workflow. Ad hoc scripting can invoke individual tools but cannot reliably assemble them into a correct and complete pipeline, as evidenced by the vanilla agents’ failure to exceed LLM-level performance.

MolClaw’s L2 workflow for binding affinity prediction addresses this gap directly. By encoding the validated sequence of operations (format conversion ? receptor preparation ? docking box definition ? Vina-GPU docking ? score extraction and comparison) as a reusable skill template, MolClaw ensures that each step is executed with correct parameters and proper error handling, regardless of the underlying LLM. The resulting 81.1% accuracy on MolClaw-CC (30/37 correct; 95% Wilson CI: [65.8%, 90.5%]; Fig. 4B) represents a qualitative shift from near-chance performance to reliable prediction.

This finding has broader implications for the design of scientific AI agents: it suggests that the primary value proposition of hierarchical skill architectures lies not in accelerating tasks that are already solvable through ad hoc scripting, but in enabling tasks that require validated multi-step computational expertise—a capability class that is fundamentally different from code generation.

#### Supplementary Note 4. Heterogeneity and unpredictability of LLM performance across tasks

The eight standalone LLMs evaluated in this study exhibit marked performance heterogeneity both within and across tasks (Tables 1–2; Fig. 3A–F, Fig. 4A). This heterogeneity has important implications for the design of drug discovery agents.

##### Within-task variance

The coefficient of variation (CV) across LLMs ranges from 33% (binding affinity accuracy) to 62% (property filtering accuracy), with molecule editing accuracy (CV = 40%) and optimization success rate (CV = 48%) falling in between. For optimization success rate, the absolute range spans from 5.1% (Minimax 2.5) to 100.0% (Kimi 2.5)—a 94.9 percentage point gap— indicating that LLM capabilities for molecular property optimization are highly model-dependent (Table 2; Fig. 3D).

##### Cross-task inconsistency

No single LLM ranks consistently at the top across all tasks. The Spearman rank correlation between LLM performance on property filtering accuracy and binding affinity accuracy is *ρ* = 0.174 (P = 0.68), indicating near-zero association. This absence of correlation is exemplified by two striking cases: GLM 5 achieves the highest docking hit count among all LLMs (0.48; Fig. 3B) but ranks seventh of eight on binding affinity accuracy (24.3%; Fig. 3A), while Gemini 3 exhibits the reverse pattern—ranking first on binding affinity (56.8%) but last among LLMs on docking hit count (0.16). This dissociation suggests that the molecular reasoning capabilities tested by different MolBench sub-tasks are largely independent and cannot be predicted from a model’s performance on any single task.

Correlations between other task pairs are somewhat stronger but remain mostly non-significant at the *n* = 8 sample size: property filtering versus molecule editing (*ρ* = 0.577, P = 0.13), binding affinity versus molecule editing (*ρ* = 0.602, P = 0.11). The two significant correlations observed—molecule editing versus optimization success rate (*ρ* = 0.714, P = 0.047) and binding affinity versus optimization success rate (*ρ* = 0.819, P = 0.013)—likely reflect a shared dependence on chemical structure reasoning ability.

##### Implications for agent design

The unpredictability of LLM performance profiles reinforces the rationale for MolClaw’s model-agnostic architecture. Because no single LLM excels uniformly across all drug discovery sub-tasks, a system that relies on the native reasoning capabilities of a specific LLM will inevitably exhibit blind spots. MolClaw’s hierarchical skill layer externalizes domain expertise into reusable, validated workflow templates, rendering the system’s overall performance less sensitive to the idiosyncratic strengths and weaknesses of the underlying LLM. This is empirically confirmed by the observation that MolClaw achieves top-ranked performance on both the Claude Code platform (powered by Claude Sonnet 4.6) and the OpenClaw platform (powered by a different LLM backbone), despite the substantial performance differences between these LLMs when used without skills (Fig. 5A– C).

#### Supplementary Note 5. Biomni and the limitations of code-based agent paradigms

Biomni, a general-purpose biomedical agent that operates by generating and executing ad hoc Python scripts for each task, exhibits a distinctive pattern of selective failure on MolBench that illuminates the limitations of purely code-based agent paradigms.

On the two sub-tasks that can be solved through single-library scripting—property filtering accuracy (74.0%) and property filtering F1 score (82.7%)—Biomni substantially outperformed all standalone LLMs (best LLM: 48.0% accuracy, 71.9% F1; Table 1; Fig. 3E–F), confirming its ability to generate and execute functional RDKit-based code. However, on binding affinity comparison, Biomni achieved only 24.3%—well below the chance level of 50% and 32.5 percentage points below the best-performing LLM (Gemini 3 at 56.8%; Fig. 3A). This below-chance performance suggests that Biomni’s code-generation approach not only fails to improve upon raw LLM reasoning but actively interferes with it, possibly by introducing computational artifacts or by executing an incorrectly configured pipeline whose outputs are systematically misleading.

More critically, Biomni failed entirely on four of seven metrics: docking screening hit count, molecule editing accuracy, optimization delta, and optimization success rate (Table 1, Table 2). These failures are not random: they occur precisely on tasks that require either multi-step tool orchestration (docking screening) or precise molecular representation manipulation (molecule editing, property optimization). The code-based paradigm, in which the agent writes a fresh script for each query without access to validated workflow templates, is fundamentally brittle for these tasks—a single incorrect parameter, missing preprocessing step, or format incompatibility can cause the entire pipeline to fail silently or produce invalid outputs.

This pattern stands in sharp contrast to MolClaw, which succeeded on every task and every metric across both platforms (Tables 1–2). The key architectural difference is that MolClaw’s hierarchical skills pre-encode the correct sequence and parameterization of tool invocations, eliminating the need for the LLM to independently discover and correctly implement complex computational workflows from scratch. The comparison between Biomni and MolClaw thus provides a controlled demonstration of the difference between “having tools” and “having skills”: both systems have access to computational chemistry tools, but only the latter embeds the domain expertise required to use them correctly in multi-step pipelines.

#### Supplementary Note 6. Validation of platform-agnostic design through cross-platform consistency

A core design principle of MolClaw is platform agnosticism: the skill layer is fully decoupled from both the underlying LLM and the agent runtime, allowing deployment on architecturally distinct platforms without modification. The dual-platform evaluation on Claude Code and OpenClaw provides a rigorous test of this principle.

##### Vanilla baselines reveal substantial platform-dependent variability

Without MolClaw skills, the two platforms exhibit divergent performance on several metrics. On docking screening, Claude Code vanilla achieves a hit count of 0.56 while OpenClaw vanilla reaches only 0.20—a 2.8 × gap (Fig. 5B). On optimization delta, the direction reverses: OpenClaw vanilla (Δ = 1.293) outperforms Claude Code vanilla (Δ = 0.866) by 49% (Fig. 5C). On optimization success rate, OpenClaw vanilla achieves 100.0% while Claude Code vanilla reaches 92.3% (Table 2). These disparities reflect differences in the platforms’ code generation strategies, error recovery mechanisms, and runtime environments, demonstrating that vanilla agent performance is highly platform-dependent.

##### MolClaw’s skills substantially reduce cross-platform variability

After equipping both platforms with MolClaw’s hierarchical skills, the performance gap narrows on most metrics. On docking screening, the gap decreases from 0.36 (0.56 versus 0.20) to 0.16 (0.80 versus 0.64)—a 56% reduction in absolute disparity (Fig. 5B). On optimization delta, the gap decreases from 0.427 (favoring OpenClaw) to 0.288 (Fig. 5C). On optimization success rate, both platforms achieve 100.0% with MolClaw skills, completely eliminating the 7.7 percentage point vanilla gap (Table 2). On property filtering, the pre-existing 2.0 percentage point gap (98.0% versus 96.0%) is preserved, which is expected since MolClaw’s skills do not intervene in this task.

The one metric where the cross-platform gap widens after skill introduction is binding affinity accuracy: vanilla platforms show identical performance (both 51.4%), while MolClaw-CC (81.1%) outperforms MolClaw-OC (73.0%) by 8.1 percentage points (Fig. 5A). This widening likely reflects an interaction between the L2 binding affinity workflow and platform-specific differences in how intermediate tool outputs are parsed and forwarded—a behavior that merits further investigation in future work but does not undermine the overall trend toward convergence.

These results support the conclusion that MolClaw’s skill architecture serves as an equalizer across platforms: by standardizing the sequence and parameterization of computational operations, the skill layer reduces the extent to which overall performance depends on platform-specific implementation details. This property is practically valuable because it means that improvements in either the LLM or the agent platform will translate into improved MolClaw performance without requiring modifications to the skill definitions themselves.

#### Supplementary Note 7. Ceiling effects, discriminative power, and the complementarity of evaluation metrics

Several MolBench metrics exhibit ceiling effects that merit discussion, both for their implications on statistical interpretation and for the methodological insights they provide about benchmark design.

##### Ceiling effects on three metrics

On property filtering accuracy, four of thirteen methods achieve ≥ 95% accuracy (Claude Code vanilla, OpenClaw vanilla, MolClaw-CC, MolClaw-OC; Table 1). On molecule editing accuracy, four methods achieve ≥ 95% accuracy (the same four agent-based systems). On optimization success rate, five methods achieve 100.0% (Kimi 2.5, OpenClaw vanilla, MolClaw-CC, MolClaw-OC, and OpenClaw vanilla). These ceiling effects reduce the statistical power of pairwise comparisons on these metrics—for example, the comparison between MolClaw-CC (100.0%) and Claude Code vanilla (97.4%) on molecule editing yields P = 1.00 by Fisher’s exact test (Fig. 5J– M)—and explain why several comparisons in Fig. 5F–I and Fig. 5J–M fail to reach significance despite positive effect sizes.

##### Binding affinity and optimization delta as high-discriminative metrics

In contrast, binding affinity comparison accuracy and optimization delta exhibit no ceiling effects and provide clear separation between method categories. On binding affinity, only MolClaw-CC exceeds 75% accuracy (Fig. 4B), and the pairwise significance matrix (Fig. 5J) shows that MolClaw-CC achieves significant superiority over 10 of 12 baselines. On optimization delta, the range spans from 0.003 (Minimax 2.5) to 1.724 (MolClaw-CC)—a 575-fold difference (Table 2). These two metrics thus carry the greatest discriminative burden in MolBench and should be prioritized in future benchmark extensions.

##### The complementarity of success rate and optimization delta

A noteworthy observation is the decoupling between optimization success rate and optimization delta among top-performing methods. Kimi 2.5, OpenClaw vanilla, MolClaw-OC, and MolClaw-CC all achieve 100.0% success rate, yet their optimization deltas differ substantially: 0.783, 1.293, 1.436, and 1.724, respectively (Table 2; Fig. 5C). This decoupling reveals that success rate—a binary measure of whether all property targets are met—does not capture the margin by which targets are exceeded. Optimization delta provides this complementary information, distinguishing between methods that barely satisfy constraints and those that achieve substantial improvement beyond the minimum thresholds. The 2.2× ratio between MolClaw-CC (Δ = 1.724) and Kimi 2.5 (Δ = 0.783), despite identical success rates, underscores the practical importance of reporting both metrics. For lead optimization campaigns where safety margins above property thresholds are desirable, optimization delta is the more informative indicator of real-world utility.

##### Implications for future benchmark design

The observation that different metrics exhibit different levels of discriminative power suggests that future versions of MolBench should include additional tasks calibrated to the capability frontier of current agent systems. Specifically, tasks in the binding affinity and multi-step workflow category—where the largest performance gaps were observed—should be expanded and diversified to provide finer-grained assessment of skill-driven capabilities. Conversely, tasks that are already saturated (property filtering, molecule editing) may benefit from increased difficulty through more complex constraint sets or larger candidate pools.

#### E2E: Design Principles and Scope

MolBench-E2E comprises three end-to-end challenges, each representing a maximally distinct workflow archetype of early-stage structure-based drug discovery. Tasks were designed according to three guiding principles:

1. **Ecological validity**. Each task mirrors a workflow that a computational chemist would encounter in a real drug discovery campaign. Task specifications were drafted by domain experts with experience in structure-based drug discovery and subsequently reviewed for scientific plausibility.
2. **Minimal ambiguity with maximal autonomy**. Tasks specify success criteria and required outputs but do not prescribe the exact sequence of tool invocations, allowing the agent to exercise autonomous planning. Where parameter choices could introduce uncontrolled variance (e.g., docking box coordinates), tasks mandate a deterministic derivation procedure (e.g., centroid of co-crystal ligand heavy atoms).
3. **Workflow archetype coverage**. The three tasks were selected to span the major categories of long-horizon computational workflows: physics-based conformational sampling, iterative property-driven molecular optimization, and structure-guided lead optimization with docking feedback. This design ensures that benchmark performance reflects generalizable workflow orchestration competence rather than task-specific memorization.

#### Distinction Between MolBench-MO and MolBench-E2E Optimization Tasks

MolBench-MO’s physicochemical property optimization subtask and MolBench-E2E Q2 both involve QED optimization. We delineate them as follows:

**MolBench-MO** evaluates *single-step* optimization capacity: given a source molecule and target property direction, the agent produces optimized molecules in a single generation pass. Performance is measured by absolute property change (Δ) and success rate. No iterative reasoning, diagnostic analysis, or strategy adaptation is required or evaluated.

**MolBench-E2E Q2** evaluates *multi-round closed-loop reasoning*: the agent must execute a structured iterative protocol with explicit diagnostic steps, maintain quantitative records across rounds, adapt strategy based on prior outcomes, enforce convergence criteria, and manage scaffold similarity constraints. The evaluation focus shifts from raw optimization magnitude to the quality of the reasoning loop—whether the agent can identify failure modes, formulate corrective hypotheses, and terminate appropriately. This mirrors the iterative nature of real medicinal chemistry campaigns, where the ability to learn from failed modifications is as important as the ability to propose improvements.

#### E2E-Q1 — Coarse-Grained Conformational Sampling and All-Atom Reconstruction

##### Target

EGFR kinase domain, PDB ID: 1M17 (chain A), resolution 2.6 Å, active conformation. Reference ligand: Erlotinib (residue name AQ4).

##### Residue Numbering Convention

PDB 1M17 uses authors’ original numbering (residues 671– 998), offset by −24 from canonical UniProt P00533 numbering. All task specifications use UniProt numbering; agents must apply the correction when working with the PDB coordinate file.

##### Workflow Specification

*Phase 1: Structure Preprocessing and CG Input Preparation*.

a. Download 1M17; extract chain A; remove ligand (AQ4), crystallographic waters, and non-standard residues. Save cleaned all-atom reference structure. Record sequence length and secondary structure distribution (helix/sheet/coil proportions).
b. Prepare inputs for two CG force fields:

- **GoCa:** Extract C*α* coordinates; construct native contact map; generate GoCa-format input files.
- **OpenAWSEM:** Use the OpenAWSEM toolchain to generate CG model input from the PDB file, including sequence file, CG coordinates, and force field parameters.

*Phase 2: CG Simulation and Conformational Sampling*.

(c) GoCa CG-MD simulation: run dynamics; save snapshots at uniform intervals; extract 10 representative conformations at equal spacing from the production trajectory (excluding equilibration).
(d) OpenAWSEM CG-MD simulation: extract 10 representative conformations in the same manner.

*Phase 3: All-Atom Reconstruction (PULCHRA)*.

(e) Reconstruct all-atom structures from GoCa C*α*-only conformations using PULCHRA. Save 10 all-atom PDB files.
(f) Reconstruct all-atom structures from OpenAWSEM CG conformations (C*α*, C*β*, O) using PULCHRA. Save 10 all-atom PDB files.

##### Estimated Tool Invocations

Structure retrieval → PDB cleaning (PDBFixer) → GoCa input preparation → GoCa simulation → trajectory extraction → OpenAWSEM input preparation → OpenAWSEM simulation → trajectory extraction → PULCHRA reconstruction (× 20): approximately 8–12 distinct MCP tool invocations.

##### Evaluation Rubric

**Evaluation Rubric.**
| Criterion | Weight | Full Credit |
| --- | --- | --- |
| Structure preprocessing | 15% | Chain extraction, ligand/water removal, secondary structure annotation |
| GoCa input preparation and simulation | 20% | Valid contact map, successful CG-MD run, 10 conformations extracted |
| OpenAWSEM input preparation and simulation | 20% | Valid CG model, successful simulation, 10 conformations extracted |
| PULCHRA reconstruction (GoCa) | 15% | 10 all-atom PDB files with correct atom counts |
| PULCHRA reconstruction (OpenAWSEM) | 15% | 10 all-atom PDB files with correct atom counts |
| Output organization and reporting | 15% | Both pipelines documented; files organized and accessible |

#### E2E-Q2 — QED-Driven Iterative Molecular Optimization (5 Rounds)

##### Starting Molecule

Butyl 2-[6-(4-chlorophenyl)-8-methoxy-1-methyl-4H-[1,2,4]triazolo[4,3-a][1,4]benzodiazepin-4-yl]acetate.

SMILES: `CCCCOC(=O)CC1N=C(c2ccc(Cl)cc2)c2cc(OC)ccc2-n2c(C)nnc21`

MW: 452.94 g/mol. Baseline QED ≈ 0.50–0.55 (unweighted).

##### Design Rationale

The triazolo-benzodiazepine scaffold contains three fused/pendant aromatic rings, imposing an inherent aromatic ring desirability ceiling (~ 0.27 for the AROM QED component). Achieving QED ≥ 0.70 from a ~ 0.55 baseline therefore requires compensatory optimization across MW, LogP, and rotatable bond dimensions—a non-trivial multi-round challenge that cannot be resolved by a single structural edit.

##### Task Specification

Iteratively optimize QED over ≤ 5 rounds. Target: QED ≥ 0.70, Tanimoto similarity ≥ 0.40 to starting molecule (Morgan fingerprints, radius 2).

##### Per-Round Protocol (Mandatory)

**Per-Round Protocol (Mandatory).**
| Step | Description |
| --- | --- |
| Assess | Calculate MW, LogP, TPSA, HBD, HBA, rotatable bonds, aromatic ring count, QED; predict ADMET profile |
| Diagnose | Identify QED-limiting features; for Rounds 2–5, explicitly state prior strategy, outcome, and remaining bottleneck |
| Design | Propose 3–5 modified molecules, each targeting 1–2 features with chemical rationale; all must maintain Tanimoto $\geq 0.40$ |
| Verify | Calculate QED and full properties for all proposals; select best as next-round seed |

##### Termination Rules

Stop early if QED ≥ 0.70 achieved. If no improvement for 2 consecutive rounds, declare convergence and report global best. Round *N* + 1 strategy must differ from Round *N* if Round *N* failed.

##### Required Output

Optimization trajectory table: Round, SMILES, QED, MW, LogP, TPSA, Tanimoto similarity, strategy label, outcome.

##### Estimated Tool Invocations

12–15 (molecular property calculation × rounds × candidates, ADMET prediction, similarity computation).

##### Evaluation Rubric

**Evaluation Rubric.**
| Criterion | Weight | Full Credit |
| --- | --- | --- |
| Property calculation accuracy | 15% | QED, LogP, MW computed correctly via validated tools |
| Diagnostic reasoning quality | 25% | Each round identifies specific structural bottleneck with quantitative justification |
| Chemical validity and similarity maintenance | 15% | All proposals valid SMILES; Tanimoto $\geq 0.40$ maintained |
| Strategy adaptation across rounds | 25% | Strategy explicitly informed by prior round’s quantitative results; pivot on failure |
| Target achievement and convergence behavior | 20% | QED $\geq 0.70$ reached, or appropriate convergence declaration |

#### E2E-Q3 — Structure-Guided Iterative Lead Optimization

##### Target and Lead Compound

EGFR kinase domain, PDB ID: 1M17 (chain A). Lead: Erlotinib. SMILES: `COCCOC1=C(C=C2C(=C1)C(=NC=N2)NC3=CC=CC(=C3)C#C)OCCOC`

##### Key Binding-Site Residues (Reference)

**Key Binding-Site Residues (Reference).**
| Functional Role | Residue (PDB#) | Residue (UniProt#) |
| --- | --- | --- |
| Hinge region | Met769 | Met793 |
| Hinge region | Leu768 | Leu792 |
| Gatekeeper | Thr766 | Thr790 |
| Catalytic lysine | Lys721 | Lys745 |
| αC-helix | Glu738 | Glu762 |
| DFG motif | Asp831 | Asp855 |
| P-loop | Gly695–Val702 | Gly719–Val726 |
| Hydrophobic pocket | Leu694, Val702, Ala719, Leu820 | Leu718, Val726, Ala743, Leu844 |

##### Optimization Objective

Achieve QuickVina docking score improvement ΔScore ≤ −2.0 kcal/mol relative to Erlotinib baseline for ≥ 2 molecules. Maximum 15 rounds.

##### Workflow

a. *Baseline Establishment*. Download 1M17 → extract AQ4 centroid (mean of all heavy-atom coordinates) before removing the ligand → remove ligand/waters → prepare receptor (PDBFixer) → dock Erlotinib (QuickVina, box center = AQ4 centroid, box size = 25 × 25 × 25 Å) → record Score_baseline and baseline molecular properties (MW, LogP, HBD, HBA, rotatable bonds, TPSA, Lipinski/Veber status). Lock docking parameters for all subsequent rounds.
b. *Modification Map*. Define quinazoline core SMARTS pattern. Enumerate ≥ 4 modifiable positions (C6/C7 methoxyethoxy side chains, C4-NH aniline moiety, C2 position on quinazoline, terminal ethynyl group). Map each position to nearby binding-site residues and hypothesized interaction improvements. Record as persistent reference.
c. *Iterative Optimization (per round):*

1. **Strategize:** Formulate modification hypothesis (target position, structural rationale, proposed change type). Record before generation.
2. **Generate:** Attempt REINVENT4 mol2mol generation; supplement with manually designed analogs if needed. Label source (REINVENT/manual). Apply scaffold check (quinazoline substructure) and drug-likeness filter (Lipinski ≤ 1 violation, Veber compliant).
3. **Dock:** Dock ≤ 10 filtered molecules with locked parameters.
4. **Evaluate:** Compute ΔScore for each molecule. If ΔScore ≤ −2.0, mark as “target met.” Write SAR summary. Select best-scoring molecule as next-round seed. Update global target tracker. Check termination.

##### Strategy Pivot Rule

If best score does not improve for 3 consecutive rounds, agent must explore a different modifiable position or attempt multi-site modification.

##### Termination Conditions

- **Success:** ≥ 2 molecules across all rounds have met ΔScore ≤ −2.0 → stop immediately.
- **Failure:** Round 15 completed with *<* 2 molecules meeting target → stop.

##### Required Output

- `run_log.md`: Per-round entries with modification hypothesis, generated molecules (with source labels), docking results table, SAR summary, round decision.
- `final_report.md`: Locked parameters table, modification map summary, optimization trajectory table, global target tracker, SAR narrative across all rounds, conclusion (SUCCESS/FAILURE, total rounds, best molecule).

##### Estimated Tool Invocations

20–50+ (receptor preparation, baseline docking, REINVENT generation × rounds, QuickVina docking × rounds × candidates, property calculation, scaffold filtering).

##### Evaluation Rubric

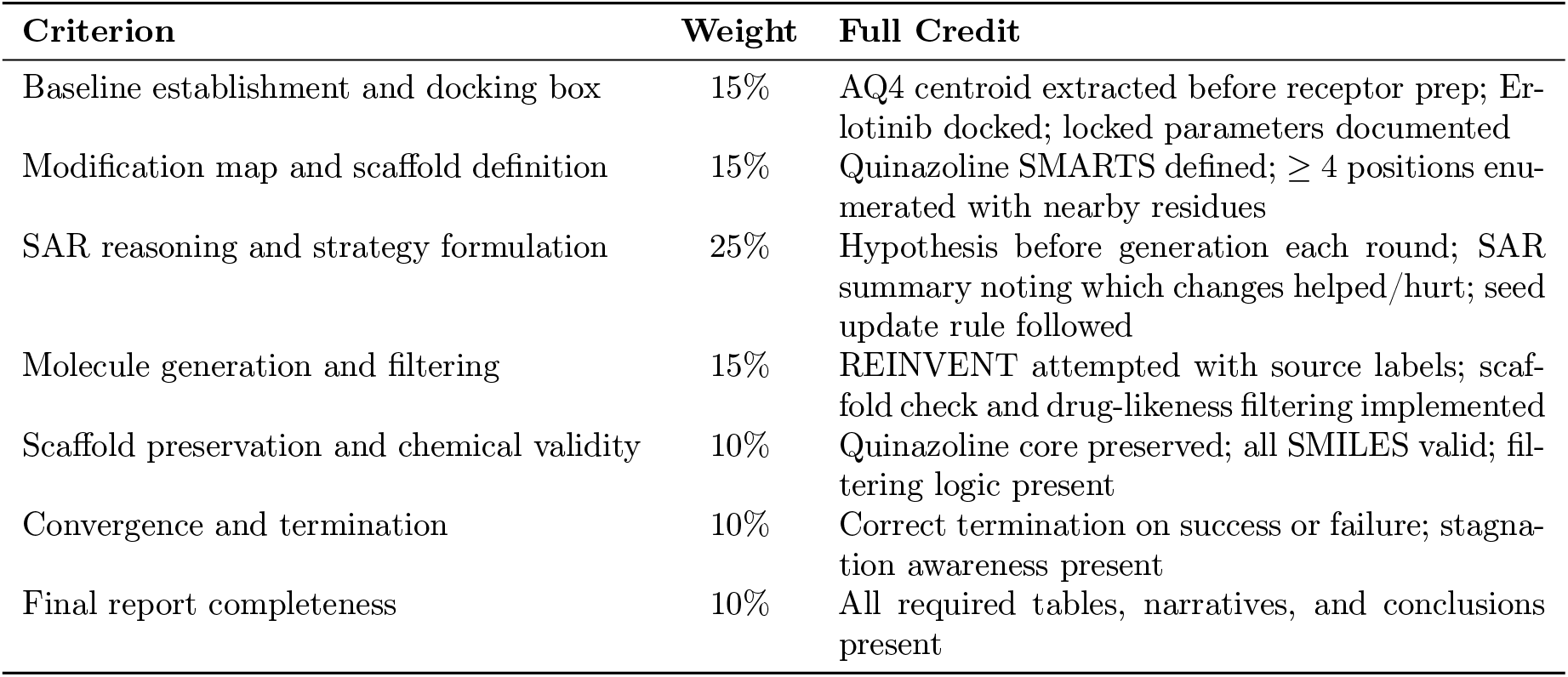

#### Evaluation Protocol

All MolBench-E2E tasks are evaluated by three independent domain-expert raters using the task-specific rubrics defined above. Each criterion is scored on a 0–2 scale (0 = not attempted or fundamentally incorrect; 1 = partially correct with significant gaps; 2 = fully correct). The weighted sum across criteria yields a normalized task score in [0, 1]. Inter-rater agreement is quantified by Krippendorff’s *α*, with disagreements resolved by majority vote. Aggregate E2E performance is reported as the mean normalized score across all 3 tasks.

#### Future Benchmark Expansion

The current MolBench-E2E comprises three tasks selected to represent maximally distinct workflow archetypes. Expanding E2E coverage to additional archetypes—including multi-target selectivity profiling, cryptic pocket discovery via conformational ensemble screening, binding free energy-guided optimization, and protein–protein interaction modulator design—is planned for future benchmark releases.

#### E2E-Q2: Design Principles and Scope

##### 1. Predicting the optimization ceiling from scaffold topology

A central finding of this study is that the triazolo-benzodiazepine scaffold imposes a hard upper bound on the achievable QED score. Here we derive this bound analytically. QED is defined as a weighted geometric mean of eight component desirability functions *d*_*i*_ (ref. [9]):

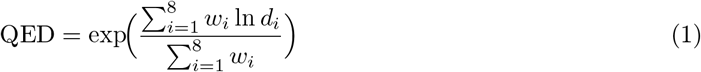

where the weights *w*_*i*_ follow the mean-weight parameterization of Bickerton et al. (MW = 0.66, ALogP = 0.46, HBA = 0.05, HBD = 0.59, PSA = 0.06, RotB = 0.65, AROM = 0.48, ALERTS = 0.95; total *W* = 3.90). Because the scaffold retains three aromatic rings in all viable derivatives—reducing to two would dismantle the triazolo-benzodiazepine pharmacophore—the AROM desirability is locked at *d*_AROM_ = 0.257. We can therefore compute the theoretical QED ceiling by assuming all other seven components are maximized. If the remaining components each reach *d* = 1.0 (the theoretical perfect score), the ceiling is:

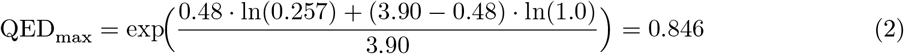

Under more realistic assumptions (*d* = 0.99 for each non-AROM component), the ceiling falls to 0.839; at *d* = 0.96, it is 0.816. The R4 molecule achieves QED = 0.799, corresponding to 94–98% of these ceiling estimates depending on the assumed maximum for the other components. By R4 the seven non-AROM desirabilities averaged 0.938, indicating that most individual dimensions were already near-optimal and little room for further improvement remained.

This ceiling prediction has practical implications: given a scaffold’s aromatic ring count, one can estimate the maximum attainable QED *before* running any optimization, providing an a priori benchmark against which to evaluate agent performance. More broadly, any scaffold feature that fixes a QED component—for example, a mandatory macrocyclic ring system that locks the rotatable bond count—will impose an analogous ceiling that can be calculated by the same formula.

##### 2. Tanimoto budget exhaustion as a convergence diagnostic

In our main Results we noted that the qualification rate—the fraction of generated molecules satisfying the Tanimoto ≥ 0.40 constraint—declined from 100% (R1–R2) to 57.6% (R5). We propose that this declining rate constitutes a generalizable convergence diagnostic that we term “Tanimoto budget exhaustion.”

The mechanism is straightforward. At each round, the agent selects the best qualifying molecule as the next seed. Because higher QED generally requires greater structural divergence (Fig. 7C), the seed’s Tanimoto to the original starting molecule decreases monotonically: 1.000 → 0.769 → 0.687 → 0.549 → 0.452 → 0.465. When the generative model is seeded with a molecule that already sits at Tanimoto = 0.452, it must simultaneously satisfy two conflicting requirements: (i) remain similar to the *seed* (a condition enforced by the generative prior), and (ii) remain similar to the *starting molecule* (the external constraint). As the seed drifts further from the starting molecule, the intersection of these two similarity spheres shrinks, reducing the fraction of valid candidates.

We observed that the qualification rate dropped below 60% at the same round (R5) at which the agent formally declared convergence on QED grounds, suggesting it can serve as a corroborating convergence signal. This suggests a complementary stopping criterion: when fewer than 50% of generated molecules satisfy the similarity constraint, the Tanimoto budget is effectively exhausted and further rounds are unlikely to yield qualifying improvements. Such a metric is scaffold-agnostic and does not require knowledge of the QED ceiling, making it applicable to any iterative constrained optimization campaign regardless of the target property.

##### 3. Phase transitions versus gradual improvement in property optimization

Not all QED components improved gradually. The structural alerts (ALERTS) desirability exhibited a discontinuous phase transition: 0.241 at R0 to 0.842 at R1, with no further change in R2–R5. This occurred because the starting molecule’s butyl ester triggered two Brenk structural alerts; once the ester was removed at R1, the alert count fell to zero and the desirability jumped to its maximum achievable value.

This phase-transition behaviour contrasts sharply with the continuous, gradual improvement of MW (five incremental reductions across R1–R4) and ALogP (four stepwise decreases). The distinction carries strategic significance: structural alerts are binary liabilities that should be targeted first, as their removal yields one-time, disproportionately large QED gains. The agent implicitly followed this strategy—the R1 modification was the most impactful single change, simultaneously clearing alerts, reducing MW, and trimming rotatable bonds—but did not explicitly articulate this prioritization. An ideal agent architecture would decompose QED into its components at the outset, identify binary versus continuous liabilities, and sequence modifications accordingly.

##### 4. The interpretability–efficiency trade-off in generative design

The evaluation question instructed the agent to “propose 3–5 modified molecules with chemical rationale.” Instead, the agent employed REINVENT4 batch generation to produce 23–54 candidates per round and selected the best by QED ranking. This substitution raises a fundamental question about AI agent design: should agents demonstrate interpretable design reasoning, or is outcome-optimal search sufficient?

The batch approach has clear advantages. It explored a far larger chemical space (182 molecules total) than the requested 3–5, enabling the discovery of modifications—such as the simultaneous OMe removal and alcohol-to-amide conversion at R3—that a human designer might not have considered in combination. It also achieved QED = 0.799, likely exceeding what hand-picked proposals would have reached.

However, interpretability was sacrificed. The agent could not explain *why* a particular molecule was generated, only why it was *selected*. For regulatory contexts that require design rationale (for example, ICH M7 mutagenicity assessments), this opacity is problematic. A hybrid architecture—using generative models for broad exploration followed by explicit decomposition of the selected molecule’s structural changes—would preserve the coverage advantage while restoring interpretability. We note that the agent’s post-hoc structural analysis (identifying the butyl ester removal, OMe deletion, and Cl removal as discrete steps) partially fulfils this role but is retrospective rather than prospective.

##### 5. Systematic blind spot detection in multi-objective optimization

The agent’s failure to detect the AMES mutagenicity deterioration (+180%, from 0.165 to 0.462) despite tracking over 13 ADMET endpoints illustrates a general vulnerability of attention-based monitoring. The agent explicitly tracked CYP3A4, hERG and DILI at each round—all of which improved—but AMES was not included in its round-by-round diagnostic summaries.

This omission likely reflects an attentional bias: the agent focused on endpoints that were problematic at baseline (CYP3A4 = 0.968, DILI = 0.839, hERG = 0.601) while neglecting endpoints that started in a safe range (AMES = 0.165). The gradual, incremental nature of the AMES increase (0.092 → 0.199 → 0.359 → 0.462 across R1–R4) further reduced its salience at any single round.

We propose that future agent architectures incorporate an “unmonitored endpoint alarm”: after each round, automatically compare all ADMET endpoints between the current best and the baseline, flagging any that have crossed a predefined deterioration threshold (for example, *>*0.15 absolute increase or *>*100% relative increase). Such a mechanism operates independently of the agent’s explicit attention allocation and would have caught the AMES trend at R3 (0.359, +118% from baseline). This is analogous to monitoring systems in clinical trials that track safety signals beyond the pre-specified primary endpoints.

##### 6. Cost-effectiveness and practical stopping rules

The diminishing returns pattern (Fig. 7H) has direct implications for computational resource allocation. Assuming roughly equal tool-call costs per round, and measuring against the R0-to-R4 improvement (+0.4216) since R4 is the recommended molecule, R1 delivers 83.0%, R1–R2 deliver 88.1%, R1–R3 deliver 94.0%, and R1–R4 deliver 100%. The R5 contribution is negligible (+0.0004, or 0.1% of the R0–R4 total).

From a cost-effectiveness perspective, a three-round campaign (R1–R3) captures 94% of the ultimate improvement at 60% of the total computational cost (3/5 rounds). The marginal cost-effectiveness ratio—defined as additional QED gain per additional round—drops from 0.350 (R1) to 0.022–0.025 (R2–R4) to 0.0004 (R5), an approximately 800-fold decrease from R1 to R5.

This suggests two practical stopping rules for production deployment: (i) an *absolute threshold rule* (stop when QED exceeds the target; here met at R1), and (ii) a *marginal improvement rule* (stop when round-over-round gain falls below a fraction of R1’s gain; for example, *<*1% of R1’s marginal gain, which would have triggered at R5). Rule (ii) is more conservative but protects against premature termination when the target has not yet been reached. Combining both with the Tanimoto budget metric from Section 2 yields a three-criterion ensemble that provides robust convergence detection without requiring knowledge of the theoretical QED ceiling.

#### E2E-Q3: Design Principles and Scope

##### 1. Multi-round iterative optimization as an emergent agent capability

The E2E-Q3 task required the AI agent to execute an iterative closed-loop optimization cycle— Strategize, Generate, Dock, Evaluate—across up to 15 rounds, with autonomous decision-making at each round boundary. This represents a fundamentally different challenge from single-step computational tasks, as it demands that the agent maintain a coherent optimization trajectory while accumulating and applying SAR knowledge from prior iterations. The agent successfully converged in 6 of the allowed 15 rounds, achieving the ΔScore ≤ −2.0 target with two structurally distinct molecules. The locked-parameter design—fixing docking box center and size across all rounds—ensures that score differences between molecules are attributable to structural modifications rather than docking-setup variability, and we propose this as a standard practice for future AI-driven lead optimization benchmarks.

##### 2. Long-range planning, self-repair, and emergent medicinal chemistry knowledge

The agent autonomously authored four pipeline versions (v1–v4, totaling 163 KB of Python), progressively diagnosing and recovering from crashes: v1 failed due to NumPy/RDKit incompatibility, v2 succeeded through R1 but crashed on an f-string bug, v3 resumed from R2 with pre-programmed R2–R5 strategies and executed through R4, and v4 resumed from R5 with pre-planned R5–R8 analog lists and achieved the second target-met molecule in R6. This iterative self-repair demonstrates not fragility, but resilience—the agent’s strategic reasoning remained intact despite repeated toolchain failures. Critically, the skills framework contained zero target-specific hints (grep for “erlotinib”, “EGFR”, “methoxy”, “fluorine” across all 58 skill files returned zero matches), confirming that the agent’s medicinal chemistry knowledge—methoxy shortening, halogen scanning, hydroxyl introduction, regioisomer exploration—derives entirely from LLM training data, representing an emergent drug design capability.

##### 3. Agent versus REINVENT: complementary collaboration rather than competition

The 3:3 tie in round winners and the non-significant pooled comparison (*p* = 0.104) mask a deeper complementarity. REINVENT excelled at creative molecular recombination: it serendipitously discovered methoxy shortening in R1 (not hypothesized by the agent), generated the F+OH+CH_3_ motif in R3, and produced the diF-OH target-met compound in R4 by recombining seed features. The agent excelled at hypothesis-driven systematic scanning: it designed the meta-F molecule that won R2 (precisely matching its “Br → F” hypothesis), and produced systematic regioisomer libraries for R5–R6 when REINVENT output dropped to zero. The agent’s most critical contribution was not hypothesis accuracy (which failed three times due to “ethynyl fixation”) but seed selection—consistently choosing the best-scoring molecule as the next round’s seed, thereby propagating productive scaffolds forward regardless of hypothesis quality.

##### 4. Optimization dynamics: stepwise improvement and performance plateau

The optimization exhibited a characteristic two-phase dynamics. In the improvement phase (R1–R4), the best score improved monotonically (−7.4 → −8.0 → −8.3 → −8.9), with per-round mean scores rising from −7.00 to −8.05 (Fig. 10B). From Round 2 onward, all rounds were significantly better than baseline (*p <* 0.01). In the plateau phase (R4–R6), the round mean stabilized at approximately −7.95 kcal/mol, and all 30 molecules scored below baseline (100% improvement rate vs. 79% in R1– R3). The R5 best score (−8.4) regressed from R4’s −8.9 before R6 recovered to −8.9, demonstrating that the agent maintained sufficient chemical diversity to escape local optima even within the plateau.

##### 5. Structural divergence–interaction conservation decoupling

Perhaps the most scientifically notable finding is the decoupling of structural novelty from interaction capacity. Tanimoto similarity decreased from 0.60 (R1) to 0.38 (R4–R6)—a highly significant trend (*ρ* = −0.691, *p* = 7.4 × 10^−9^)—while the quinazoline core was preserved in 54/54 molecules (100%). Despite this structural divergence, the H-bond residue count remained stable at 1.4–1.9 per molecule across all rounds (Kruskal–Wallis *p* = 0.89), fluctuating narrowly around the Erlotinib reference value of 2 (Fig. 10E). The more dissimilar molecules achieved significantly better scores (*ρ* = +0.650, *p <* 10^−7^; Fig. 10I), confirming that productive optimization required moving beyond conservative modifications. This decoupling—diversifying non-polar regions while preserving polar anchoring contacts—is a hallmark of rational lead optimization and suggests that the AI agent implicitly followed sound medicinal chemistry principles without explicit instruction.

At the residue level, the optimization executed a specific H-bond exchange: Erlotinib’s Asp831 (DFG) contact was lost (100% → 9%), replaced by the Met769 (hinge) contact (0% → 57%). Forest-plot analysis demonstrated that only Met769 significantly improved docking scores (Δ = −0.41 kcal/mol, *p* = 0.007), while Asp831 loss had no detectable impact (*p* = 0.362). This selective acquisition of the score-relevant H-bond and dispensation of the non-contributing one occurred without explicit interaction-level feedback—the agent had no access to per-residue interaction data during optimization, as all ProLIF attempts failed.

##### 6. Dual convergent binding modes and chemical space exploration

The two target-met molecules (R4 and R6, both −8.9 kcal/mol) achieved the same docking score through fundamentally different hydrogen-bond networks. The R4 compound (2,6-difluoro-4-hydroxyphenyl aniline) engaged four H-bonds via the DFG-adjacent pathway (Met769, Thr766, Thr830, Asp831), while the R6 compound (fluoro-hydroxy-methyl aniline) engaged three H-bonds via the catalytic-lysine pathway (Lys721, Thr766, Met769). This dual-pathway convergence demonstrates that the EGFR inhibitor optimization landscape contains multiple degenerate minima accessible from the same starting scaffold. The R6 winner is structurally a regioisomer of the R3 best compound (Tanimoto = 0.714, differing only in F/OH ring positions), yet scored 0.6 kcal/mol better (−8.9 vs. −8.3), highlighting the sensitivity of binding affinity to precise substituent placement.

##### 7. Limitations and future directions

Several limitations merit discussion. First, the agent exhibited persistent “ethynyl fixation”—hypothesizing restoration of Erlotinib’s terminal ethynyl group in Rounds 1, 3, and 4, despite the best molecules consistently lacking this group. This reflects a tendency of LLM agents to anchor on features of the parent molecule rather than learning from negative results. Second, REINVENT4 output declined progressively (5 → 4 → 2 → 11 → 0 → 0 molecules in R1–R6), with complete failure in R5–R6, suggesting that generative models trained on drug-like space may struggle when the seed molecule diverges substantially from the training distribution (Tanimoto *<* 0.4). The agent compensated through manual design, demonstrating system-level resilience even when individual components failed. Third, the agent’s SAR reasoning remained template-based (2–3 sentences per round), lacking mechanistic depth regarding conformational rigidity, solvation penalties, or entropic contributions. Fourth, the agent’s termination was driven by the predefined success criterion (≥ 2 target-met molecules) rather than an autonomous assessment of optimization convergence, though the stagnation counter in v3/v4 code indicates awareness of diminishing returns. Future work should explore integration of real-time interaction feedback (e.g., functional ProLIF or PLIP) to enable interaction-aware hypothesis generation, and investigate whether chain-of-thought prompting can improve the depth of SAR reasoning beyond template responses.

